# Somewhere I belong: phylogeny and morphological evolution in a species-rich lineage of ectoparasitic flatworms infecting cichlid fishes

**DOI:** 10.1101/2021.03.22.435939

**Authors:** Armando J. Cruz-Laufer, Antoine Pariselle, Michiel W. P. Jorissen, Fidel Muterezi Bukinga, Anwar Al Assadi, Maarten Van Steenberge, Stephan Koblmüller, Christian Sturmbauer, Karen Smeets, Tine Huyse, Tom Artois, Maarten P. M. Vanhove

## Abstract

A substantial portion of biodiversity evolved through adaptive radiation. However, the effects of explosive speciation on species interactions remain poorly understood. Metazoan parasites infecting radiating host lineages could improve our knowledge because of their intimate host relationships. Yet limited molecular, phenotypic, and ecological data discourage multivariate analyses of evolutionary patterns and encourage the use of discrete characters. Here, we assemble new molecular, morphological, and host range data widely inferred from a species-rich lineage of parasites (*Cichlidogyrus*, Platyhelminthes: Monogenea) infecting cichlid fishes to address data scarcity. We infer a multi-marker (28S/18S rDNA, ITS1, COI mtDNA) phylogeny of 58/137 species and characterise major lineages through synapomorphies inferred from mapping morphological characters. We predict the phylogenetic position of species without DNA data through shared character states, a combined molecular-morphological phylogenetic analysis, and a classification analysis with support vector machines. Based on these predictions and a cluster analysis, we assess the systematic informativeness of continuous characters, search for continuous equivalents for discrete characters, and suggest new characters for morphological traits not analysed to date. We also model the attachment/reproductive organ and host range evolution using the data of 136/137 described species and multivariate phylogenetic comparative methods (PCMs). We show that discrete characters can mask phylogenetic signals but can be key for characterising species groups. Regarding the attachment organ morphology, a divergent evolutionary regime for at least one lineage was detected and a limited morphological variation indicates host and environmental parameters affecting its evolution. However, moderate success in predicting phylogenetic positions, and a low systematic informativeness and high multicollinearity of morphological characters call for a revaluation of characters included in species characterisations.

## Introduction

### Adaptive radiations and host-parasite interactions

Adaptive radiation is one of the most important processes of species formation. These explosive speciation events might explain a substantial part of the biodiversity on the planet (Glor, 2010). Adaptive radiations are characterised by a rapid diversification resulting from adaptation to newly available ecological niches (Losos, 2010). Famous examples include Darwin’s finches (Grant, 1999), Caribbean lizards of the genus *Anolis* Daudin, 1802 (Mahler et al., 2013), cichlid fishes (Salzburger, 2018), and Hawaiian silverworths (Landis et al., 2018). Despite the attention these species have receive, few studies have investigated the evolution of ecological interactions involving these groups and other organisms of other groups beyond the feeding ecology of the former (e.g. Guerrero and Tye, 2009; Takahashi and Koblmüller, 2011; but see Karvonen and Seehausen, 2012; Blažek et al., 2018). How do organisms evolve that accompany the rapid diversification process of an adaptive radiation?

Metazoan parasites could provide answers to this question as they form often intimate relationships with their hosts. Thus, their evolutionary regime can be strongly impacted by the host’s evolutionary history (Huyse et al., 2005). Multiple evolutionary regimes might apply to parasites infecting radiating host lineages. First, parasites that are strongly dependent on their hosts, might experience a selection pressure to remain competitive in the arms race with the host’s defences (Kaltz and Shykoff, 1998). Structures relevant to this arms race such as attachment organs might experience a strong stabilising selection pressure as they are the parasite’s main physical connection to the host. In this scenario, the attachment organ evolution would be heavily impacted by host and environmental parameters (Kaltz and Shykoff, 1998). Second, parasites could follow the example of their hosts in terms of an (adaptive) radiation. Such co-radiations have previously been reported for insects parasitic on plants or other insects (Weiblen and Bush, 2002; Forbes et al., 2009) but quantitative support for these hypotheses in the form of evolutionary models has not been provided to date. Third, parasites could have structures that are not under strong selection pressure and, therefore, their morphology might randomly diverge over time following a pattern associated with a ‘random walk’, i.e. genetic drift or randomly fluctuating selection (Losos, 2008). Reproductive organ structures, e.g. in flatworms infecting the gills of cyprinid fish (Šimková et al., 2002), have been suggested to follow this pattern causing reproductive isolation between species.

### Species-rich but missing data: a case study for host-parasite systems

Extensive evolutionary analyses often require large molecular and morphological datasets. However, such datasets remain scarce for parasitic organisms. Metazoan parasites are often small, which makes identifying these organisms to species level notoriously difficult (de Meeûs et al., 2007). Furthermore, only a fraction of the known species is genetically characterised (Poulin et al., 2019) and most species are yet to be discovered and described (Poulin et al., 2020). Among the best-known models in adaptive radiation research, African cichlids (Cichliformes, Cichlidae) are possibly the best-studied system (Salzburger, 2018) with approximately 2000 (described and undescribed) species reported from Eastern Africa alone (Turner et al., 2001; Salzburger et al., 2014). This knowledge is rooted in a long and productive tradition of international cichlid research (see Ronco et al., 2020; Van Steenberge et al., 2011). Resulting from this scientific interest, one group of gill parasites (Fig. 1a) infecting these fishes, the monogenean flatworms belonging to *Cichlidogyrus* sensu Paperna, 1960 [incl. the nested genus *Scutogyrus* Pariselle & Euzet, 1995 (Wu et al., 2007), together referred to as *Cichlidogyrus* in the following] (Fig. 1b, c), has been studied in more depth than other species-rich parasite genera, in particular from the African continent (Cruz-Laufer et al., 2021). Species of *Cichlidogyrus* rival (Cruz-Laufer et al., 2021) and possibly exceed (see Poulin, 2014) their hosts in terms of species numbers: 137 parasite species have been described on 126 fish species (Cruz-Laufer et al., 2021). Monogenean flatworms have provided insight into parasite speciation (Meinilä et al., 2004; Šimková et al., 2013; Vanhove et al., 2015), population dynamics (Kmentová et al., 2021b), anthropogenic introductions (Šimková et al., 2019; Jorissen et al., 2020) as well as host biogeography (Barson et al., 2010; Pariselle et al., 2011; Vanhove et al., 2013, 2016). This variety of evolutionary research paired with the model system status of the host species of species of *Cichlidogyrus*, has led to the suggestion of the cichlid-*Cichlidogyrus* species network as model system for parasite speciation research and the evolution of host-parasite interactions (Pariselle et al., 2003; Vanhove et al., 2016). Recent advances in immunological (Zhi et al., 2018), pathological (Igeh and Avenant□Oldewage, 2020), genomic (Vanhove et al., 2018; Caña-Bozada et al., 2021), and microscopy research (Fannes et al., 2015) have brought the vision of a cichlid-*Cichlidogyrus* model system closer to reality [see Cruz-Laufer et al. (2021) for a detailed review of model system qualities].

**Figure 1.**
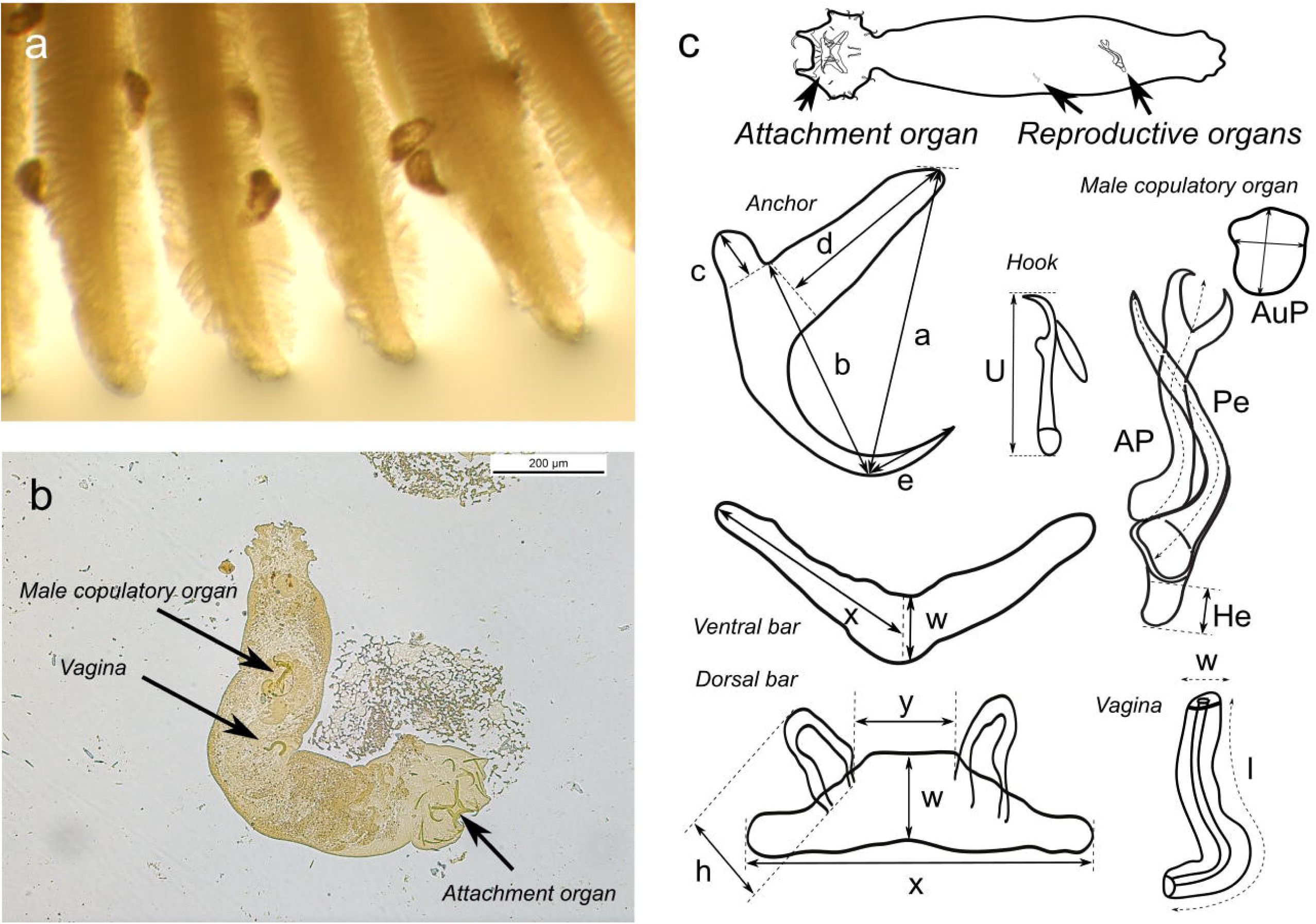
Overview of morphology of species of *Cichlidogyrus* Paperna, 1960 (Platyhelminthes: Monogenea, Dactylogyridae). (a) Multiple specimens attached to the gills of *Sarotherodon melanotheron* Rüppel, 1852 (Cichliformes: Cichlidae). (b) Microscopic image of *Cichlidogyrus agnesi* Pariselle & Euzet, 1995 with sclerotised structures of reproductive (male copulatory organ and vagina) and attachment organs indicated by arrows. (c) Overview of hard part morphology and most widely applied measurements of these structures in taxonomic literature. Abbreviations: a, anchor total length; b, anchor blade length; c, anchor shaft length; d, anchor guard length; e, anchor point length; 1–7, hook lengths; w, bar width; x, bar length; h, auricle length; y, distance between auricles; AuP, surface of auxiliary plate of male copulatory organ; AP, accessory piece length; Pe, penis length; He, heel length; l, length; w, width. Terminology and methodology of measurements according to Fannes et al. (2017).

The species-richness of *Cichlidogyrus* has previously been attributed to co-divergence with their hosts (Pariselle et al., 2003; Vanhove et al., 2015), host switching (Pariselle et al., 2003; Messu Mandeng et al., 2015), and within host speciation (Mendlová et al., 2012; Vanhove et al., 2015). Similar to other parasites, morphological characters of monogenean attachment and reproductive organ structures have formed an essential part of species descriptions. Evolutionary studies have paid considerable attention to the evolutionary patterns of these (mostly) sclerotised structures (e.g. Šimková et al., 2002; Mendlová et al., 2012). For species of *Cichlidogyrus*, the attachment organ includes a haptor with several anchors, hooks, and bars and reproductive organs with a male copulatory organ (MCO) and, at times, a sclerotised vagina (Fig. 1b, c). Based on the similar phenotypes observed in related species, some studies have suggested phylogenetic constraints, i.e. ancestry of the parasite species, as the sole determinants of the morphology of the attachment organ in *Cichlidogyrus* (Vignon et al., 2011). This claim has been questioned more recently. At least one species appears to have changed its attachment organ morphology compared to its close relatives as a response to a host switch towards non-cichlid fishes (Messu Mandeng et al., 2015). These host switches are considered rare as monogenean representatives have been reported for 27 (Carvalho Schaeffner, 2018) of 48 (Lévêque et al., 2008) fish families in Africa and most harbour some other dactylogyridean lineage of gill parasites (see Carvalho Schaeffner, 2018). Despite this discovery, no study has investigated the macroevolution of the morphological characters of the attachment and reproductive organs for almost a decade (see Mendlová et al., 2012).

The present study aims to investigate the evolutionary processes that have shaped the morphology of species of *Cichlidogyrus*. However, out of the 137 species that are currently described, DNA sequences of only 18 species were included in the most recent phylogenetic study (Messu Mandeng et al., 2015). Furthermore, morphological and ecological data have not been collected in centralised databases inhibiting the progress of large-scale meta-analytical studies (Cruz-Laufer et al., 2021). Previous studies have evaded this scarcity of information by loosely defining discrete states for the attachment organ morphology and the host repertoire base on the taxonomic literature. For instance, Vignon et al. (2011) and Mendlová et al. (2012) grouped species by the relative size of the sclerotised structures of the attachment organ (for an overview of the terminology, see Fig. 1c) using generic terms like ‘large’ and ‘small’ instead of coding these characters based on continuous measurements. Similarly, Mendlová and Šimková (2014) proposed an index of specificity (IS) to group species as species-, genus-, and tribe-specific, or generalist based on the most recent host classification at the time. Yet discretisation has been known to cause information loss as the variability of the otherwise continuous parameters is largely ignored (Altman and Royston, 2006; Goloboff et al., 2006; Parins-Fukuchi, 2018). These obstacles emphasise the need for new approaches and more extensive and accessible datasets to gain insight into the evolution of *Cichlidogyrus* and the phylogenetic information of morphological and ecological characters.

To address data scarcity, we provide new morphological, molecular, and host range data, which are made available in public databases. We perform multiple analyses to investigate the evolution of phenotypic characters to answer the following questions:

i. Which main lineages can we infer from the phylogeny? To which lineages do species without molecular data belong? Which predictions can be made through character mapping, a combined (molecular and morphological) parsimony analysis, and a machine learning algorithm?
ii. How systematically informative are the morphometric measurement most widely applied to this genus? Does the use of discrete characters lead to information loss in the morphometric and host range data?
iii. As species of *Cichlidogyrus* infect a host lineage known for its rapid speciation, which evolutionary processes have shaped the attachment and reproductive organ morphology? Are these structures under stabilising selection pressure (H_2_), do the parasites mirror the radiations of their hosts either as an early (H_3_) or a late burst (H_4_) in the character evolution, or do these structures follow a pattern associated with a random walk (H_1_)? Furthermore, do species infecting cichlids from the Eastern African radiations follow a different evolutionary regime than the species infecting other hosts (H_5_)?

## Materials and methods

### Sampling

Fish specimens belonging to Cichlidae Bonaparte, 1835 (Cichliformes), the nothobranchiid genus *Aphyosemion* Myers, 1924 (Cyprinodontiformes: Nothobranchiidae), and *Polycentropsis abbreviata* Boulenger, 1901 (Ovalentaria, *incertae sedis*: Polycentridae) were collected during various expeditions across Africa between 2008 and 2019 and from few specimens from aquaculture facilities in France (Appendix 1). As samples collected during these expeditions have been included in previous studies, sampling details can be found in the respective publications (Vanhove et al., 2011, 2013; Muterezi Bukinga et al., 2012; Pariselle et al., 2015a, 2015b; Jorissen et al., 2018a, 2018b, 2020). We dissected the gills of the fish and stored the samples in 96% ethanol to preserve the DNA of the parasites attached to the gills. Then, we screened the gills for parasitic infections of flatworm species belonging to *Cichlidogyrus*, removed the parasites from the gills using dissection needles under a stereomicroscope, mounted them temporarily in water for species-level identification at a magnification of 1000x (100x magnification with oil immersion and 10x ocular) and stored the parasite specimens individually in 96% ethanol for DNA extraction. As mostly entire specimens were used for DNA extraction, we refer to the above-mentioned articles for type and voucher specimens from the same host individuals deposited in curated collections (see Appendix 1).

### DNA extraction, amplification, and sequencing

For the DNA extraction of recent samples, we used the Nucleospin Kit (Macherey-Nagel, USA) following manufacturer guidelines but with 60 µl instead of 100 µl of elution buffer added in the final step. For some samples processed at an earlier stage, we suspended DNA samples through homogenisation with no added extraction steps following the procedures described by Marchiori et al. (2015). We amplified and sequenced three partial nuclear ribosomal genes including a fragment of the large subunit ribosomal DNA (28S rDNA) with the primers C1 (5’-ACCCGCTGAATTTAAGCAT-3’) and D2 (5’-TGGTCCGTGTTTCAAGAC-3’) (Hassouna et al., 1984), a fragment of the small subunit ribosomal DNA (18S rDNA) and the internal transcribed spacer 1 (ITS1) with the primers S1 (5′-ATTCCGATAACGAACGAGACT-3′) (Matejusová et al., 2001) and IR8 (5′-GCAGCTGCGTTCTTCATCGA-3′) (Šimková et al., 2003), and a fragment of the mitochondrial gene coding for the cytochrome oxidase *c* subunit 1 protein (COI mtDNA) with the primers ASmit1 (5’-TTTTTTGGGCATCCTGAGGTTTAT-3’) (Littlewood et al., 1997), Cox1_Schisto_3 (5’-TCTTTRGATCATAAGCG-3’) (Lockyer et al., 2003), and ASmit2 (5’-TAAAGAAAGAACATA ATGAAAATG-3’) (Littlewood et al., 1997). Reaction protocols followed the procedures of Messu Mandeng et al. (2015) for 28S rDNA, Mendlová et al. (2012) for 18S rDNA, and Vanhove et al. (2015) for COI mtDNA. We purified PCR products with a NucleoFast 96 PCR kit (Macherey-Nagel, USA) or with a GFX PCR DNA kit and Gel Band Purification kit (GE Healthcare, USA) following manufacturer guidelines. Bidirectional Sanger sequencing was conducted according to the Big Dye Terminator v3.1 sequencing protocol (Applied Biosystems, USA) at a 1:8 dilution with an ABI PRISM 3130 Avant Genetic Analyser automated sequencer (Applied Biosystems, USA) and the primers included in the PCR protocols.

### Morphological and host range data collection

We assembled morphometric (Fig. 1c) and host range data for 136 species belonging to *Cichlidogyrus.* Raw measurements taken through light microscopy were assembled from AP’s personal morphometric database and from the raw data of previous publications kindly provided by the authors of the cited studies (Pariselle and Euzet, 2003; Vanhove et al., 2011; Gillardin et al., 2012; Muterezi Bukinga et al., 2012; Pariselle et al., 2013; Řehulková et al., 2013; Van Steenberge et al., 2015; Messu Mandeng et al., 2015; Kmentová et al., 2016a, 2016b, 2016c; Rahmouni et al., 2017, 2018; Igeh et al., 2017; Geraerts et al., 2020; Gobbin et al., 2021). These raw data were deposited at MorphoBank (www.morphobank.org, project XXXXX). We calculated the standard errors and standard deviations for each measurement from the raw data. However, for species where no or only partial raw data were available, we used the mean values provided in the literature (Douëllou, 1993; Pariselle and Euzet, 2003; Řehulková et al., 2013; Rahmouni et al., 2017; Jorissen et al., 2018b, 2018a) without errors. If no mean was reported, we inferred the measures from drawings in the literature (Dossou, 1982; Dossou and Birgi, 1984; Birgi and Lambert, 1986; Muterezi Bukinga et al., 2012) and calibrating measurements through the included scale bars. Measurements of *C. dionchus* Paperna. 1968 were not included as no mean values were provided in respective publications (Paperna, 1960, 1968; Paperna and Thurston, 1969) and drawings were incomplete with some structures, e.g. the second marginal hook, entirely missing. For the auxiliary plate (AuP) of the male copulatory organ, we used the surface area and its relative error resulting of the sum of relative errors of the original measurements inferred from the plate length and width assuming an ellipsoid shape.

Beyond these continuous characters, we assigned all species to previously suggested discrete characters for the haptor morphology and host range, which include the configuration of the hooks (Vignon et al., 2011), the similarity of the anchors, the shape of the ventral bar (Mendlová et al., 2012), and the index of specificity (IS) (Mendlová and Šimková, 2014). Hook configurations were coded according to the secondary growth in the first and third hook pair (‘small-small’, ‘small-large’, ‘large-small’, ‘large-large’) relative to the second pair, which retains it embryonal size in adult specimens (Llewellyn, 1963). The anchor pairs were categorised as ‘similar’ or ‘dissimilar’ in shape and size based on the original species descriptions. The ventral bar shape was defined according to the presence of membranous extensions (see Mendlová et al., 2012) and size (‘with membranous extensions’, ‘without membranous extensions’, ‘massive with membranous extension’, ‘bar supports large extended plate (*Scutogyrus*)’). We inferred IS data from the host-parasite interactions listed by Cruz-Laufer et al. (2021) with species exclusively reported from hosts belonging to a single species, genus, or tribe classified as species-, genus-, and tribe-specific, or generalist if hosts belonging to two different tribes (or even families) were infected by the same parasite species. For the classification of cichlid species into tribes, we followed Dunz and Schliewen (2013). All discrete and continuous morphometric and host specificity data are made available at TreeBase alongside the DNA sequence alignments and tree topologies (www.treebase.org, accession number: XXXXX). Raw morphometric data are made available at MorphoBank (www.morphobank.org, project XXXXX).

We also surveyed the taxonomic literature to detect additional features used to recognise groups of related species. First, we attempted to find equivalents for characters and character states frequently mentioned in species descriptions among the morphometrics. For instance, the root lengths of the anchors in the attachment organ are frequently mentioned as ‘large’ or ‘small’, which can be reflected through the ratio of the two roots in each anchor pair. If no measurement reflected the described features, we developed new discrete variables. Detailed information on the continuous and proposed new discrete characters and their character states can be found in Appendix 2.

### Phylogenetic analyses

We assembled a four-locus concatenated multiple alignment from the sequences generated during this study and sequences available at GenBank (see Appendix 3). The alignment includes partial ITS1, 18S rDNA, 28S rDNA, and COI mtDNA sequence data. Partial DNA sequence data (i.e. specimens with less than those four loci sequenced) were included with the lacking fragments coded as missing data. We aligned the sequences of each locus using the algorithm *L-INS-i* in *MAFFT* v.7.409 (Katoh and Standley, 2013) as recommended for ribosomal DNA by the *MAFFT* manual, and removed poorly aligned positions and divergent regions using the options for less stringent parameters in *Gblocks* v0.91b (Talavera and Castresana, 2007).

We estimated tree topologies under maximum parsimony (MP) through *TNT* v1.5 (Goloboff et al., 2008b; Goloboff and Catalano, 2016) using extended implied weighting (Goloboff, 2014) in a range of values for the concavity constant K (20, 21, 23, 26, 30, 35, 41, 48, 56). Extended implied weighting reduces the impact of characters with missing data that were weighted artificially high in the original implied weighting method (Goloboff, 1993). For DNA sequence data, collectively weighting all sites in a partition (gene or codon) was suggested as a more appropriate method (Goloboff et al., 2008a). Therefore, we explored three weighting schemes proposed by Mirande (2019): all characters weighted separately (SEP), all characters weighted according to the average homoplasy of the marker (BLK), and characters in the protein-coding marker, i.e. COI, weighted according to the average homoplasy of their position with other characters were weighted as in BLK (POS). Similar to Mirande (2009), we selected the parameter combinations (k value and weighting scheme) that produced the most stable topology to compute a strict consensus tree. The distortion coefficient and subtree pruning and regrafting (SPR) distance were used as selection criteria by computing the similarity of each consensus tree obtained under the different parameters to the rest. MP tree searches involved rounds of tree fusing, sectorial searches, tree drifting, and tree ratchet (Goloboff, 1999; Nixon, 1999) under default settings and each round was stopped following three hits of the same optimum. Gaps were treated as missing data. As bootstrapping and jackknifing are reported to be distorted by differently weighted characters (Goloboff, 2003), branch support was estimated through symmetric resampling with a probability of change of 0.33 and values expressed as differences in frequencies (GC: ‘Groups present/Contradicted’). *Cichlidogyrus berrebii* Pariselle & Euzet, 1994, *C. kothiasi* Pariselle & Euzet, 1994, and *C. pouyaudi* Pariselle & Euzet, 1994, parasites of tylochromine cichlids, were used to root the phylogenetic trees due to the well-documented early diverging phylogenetic position of these species (Mendlová et al., 2012; Messu Mandeng et al., 2015).

To infer the phylogenetic position of species of *Cichlidogyrus* without DNA sequence data, we performed a second phylogenetic analysis including the morphometric measurements of the attachment and reproductive organs as a separate block in *TNT* and extended implied weighting using the same set of k values and the weighting scheme that produced the most stable tree topologies for the DNA data. The homoplasy of each morphometric character was estimated independently as recommended by Goloboff et al. (2006) for continuous characters. All other settings in *TNT* remained the same as above. The full *TNT* data matrix is provided in Supporting Information (Table S1).

For downstream analyses of character evolution, we also estimated phylogenies through Bayesian inference (BI) and maximum likelihood (ML) methods using *MrBayes* v3.2.6 (Ronquist and Huelsenbeck, 2003) on the CIPRES Science Gateway online server (Miller et al., 2010) and *IQ-Tree* v1.6.12 (Nguyen et al., 2015) respectively. We used these model-based approaches (BI and ML) to provide a consistent approach to the downstream multivariate phylogenetic comparative analyses, which are themselves model-based. DNA sequence data were partitioned by gene and, for the COI mtDNA, by codon position. We selected the substitution models for each partition according to the Bayesian information criterion (BIC) as rendered by *ModelFinder* in *IQ-TREE* (Kalyaanamoorthy et al., 2017) using partition merging (Chernomor et al., 2016) (Appendix 4). For BI analyses, we selected only models implemented in MrBayes (Appendix 4). We used two parallel runs and four chains of Metropolis-coupled Markov chain Monte Carlo iterations. We ran 20 million generations with a burn-in fraction of 0.25 and sampled the trees every 1000^th^ generation. We checked convergence criteria by assessing the average standard deviation of split frequencies (< 0.01 in all datasets) and the effective sample size (> 200) using *Tracer* v1.7 (Rambaut et al., 2018). For ML analyses, we estimated branch support values using both ultrafast bootstrap approximation (Hoang et al., 2018) and Shimodaira-Hasegawa-like approximate likelihood ratio tests (SH-aLRT) (Guindon et al., 2010) with 1000 replicates as recommended by the *IQ-Tree* manual. We considered a BI posterior probability (PP) ≥ 0.95, an ultrafast bootstrap values ≥ 95, and SH-aLRT statistic ≥ 80 as well-supported (Hoang et al., 2018). We plotted the graphs and phylogenetic trees using the *R* packages *ggplot2* v3.3.5 (Wickham, 2016) and *ggtree* v3.13 (Yu et al., 2017, 2018).

To verify congruence of the final hypothesis of the parsimony analysis (molecular data) with the BI and ML consensus trees, we analysed the congruence of the MP, BI, and ML tree topologies using the Congruence Among Distance Matrices (CADM) tests (Legendre and Lapointe, 2004; Campbell et al., 2011). We used the package *ape* v5.3 (Paradis and Schliep 2019) in *R* v4.1.0 (R Core Team, 2021) to calculate phylogenetic pair-wise distance matrices and to conduct the CADM test.

### Clade affiliation and discriminative power of morphometrics: statistical classification, literature review, and cluster analysis

Beyond the clades supported in the combined MP analysis, we also used two additional approaches to assess the phylogenetic position of species of *Cichlidogyrus* that have not been sequenced to date. First, we characterised the morphology and host repertoires of the clades inferred from the molecular phylogeny (Fig. 2) based on character maps of the continuous and discrete morphological characters and the host repertoires surveyed in the literature. To map continuous characters on the consensus MP tree, we estimated ancestral character states through the function *anc.ML* in the *R* package *phytools* v0.7-80 (Revell, 2012). To map the discrete characters, we estimated ancestral states under maximum parsimony with all rates set to equal as we could not make any assumption on the transition costs between character states. This analysis was implemented in the function *asr_max_parsimony* in the *R* package *castor* v1.6.9 (Louca and Doebeli, 2018). All character maps were plotted using the *ggplot2* and *ggtree*. Species of *Cichlidogyrus* not included in molecular phylogenies were assigned to the clades based upon the measurements and their character states while taking host repertoires (by tribe of cichlids or family of non-cichlids) into consideration. These last criteria (natural and invasive host repertoires) were taken into account as we expected the host environment to be a relevant factor in speciation processes of these parasites (see McCoy, 2003; Huyse et al., 2005). We used the host-parasite list and literature databases provided by Cruz-Laufer et al. (2021) to summarise the host repertoires.

**Figure 2.**
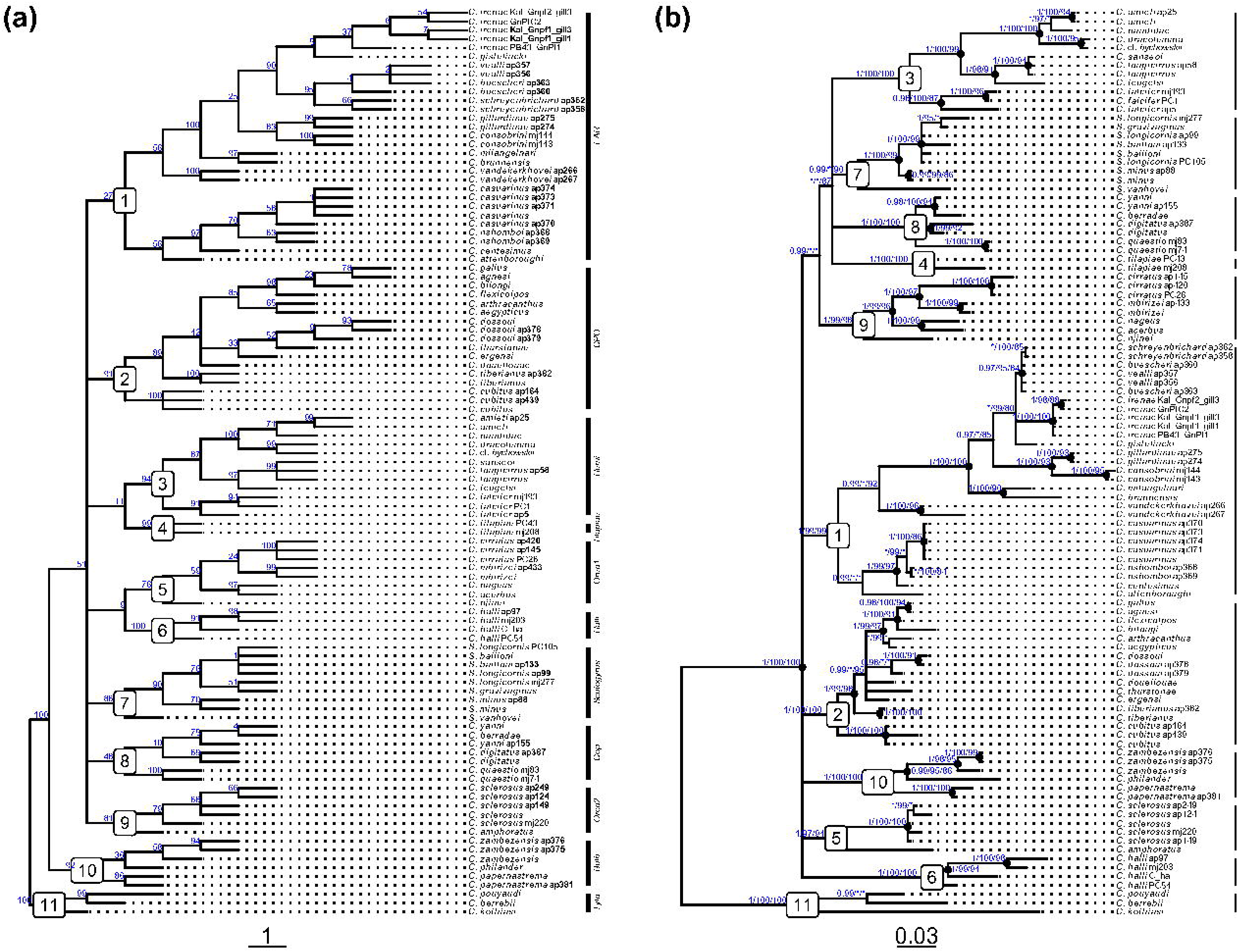
Phylograms of three monogenean flatworms belonging to *Cichlidogyrus* and *Scutogyrus* (Platyhelminthes, Monogenea, Dactylogyridae) based on three nuclear (18S, 28S, and ITS rDNA) and one mitochondrial (CO1 mtDNA) DNA sequence markers. Sequences of specimens in bold have been generated for this study. (a) Final hypothesis of analysis under maximum parsimony and extended implied weighting (k = 48, weighting scheme BLK) with node support estimated through symmetric resampling (p = 0.33). (b) Bayesian phylogram with Bayesian posterior probabilities (PP) followed by ultrafast bootstrap values (UFBoot) and Shimodaira-Hasegawa-like approximate likelihood ratios (SH-aLRT) inferred from maximum likelihood estimation indicated at nodes; asterisk (*) indicates low or moderate support below the threshold (PP < 0.95, UFBoot < 95, SH-aLRT < 80); black dots, internal nodes with strong support across all support values. Node labels (1-11), monophyletic clades considered strongly supported: species infecting (mostly) hemichromine cichlids (*Hemi*), species belonging to *Scutogyrus* (*Scutogyrus*), species infecting (mostly) coptodonine cichlids among others (*Cop*), the first species group infecting oreochromine cichlids among others (*Oreo1*), species infecting cichlids belonging to the East African Radiation (*EAR*), species infecting coptodonine, pelmatolapiine oreochromine, tilapiine, heterotilapiine, and gobiocichline cichlids (*CPO*), species from Southern Africa with a bulbous penis (*Bulb*), the second species group infecting mainly oreochromine cichlids among others (*Oreo2*), *C. halli*– clade (*Halli*), and species infecting tylochromine cichlids (*Tylo*). Abbreviation: *C. schreyenbrichard.*, *C. schreyenbrichardorum*.

Second, we predicted clade affiliation through a supervised machine learning algorithm using support vector machines (SVMs). Machine learning can be used to assess of the predictive power of morphometric measurements in taxonomic studies, in particular with support vector machines (SVMs) (e.g. Zischke et al., 2016; Fang et al., 2018). In these supervised machine learning approaches, a dataset is provided to train the learning algorithm to classify individuals in distinct groups. The predictions generated by the algorithm are then validated against a set of samples with a known affiliation. The performance can then provide an insight into the predictive power of the input data. Subsequently, the optimised algorithm can be applied to a test dataset of samples with unknown group affiliation. Here, we carried out the SVM analysis with a radial basis kernel function as implemented in the method *svmRadial* in the *R* package *caret* (Kuhn, 2008; Meyer et al., 2020). Missing data in the measurements were imputed through k-nearest neighbour imputation, scaled, and centred as implemented in the function *preProcess*. The *C* and σ parameters of the kernel function were optimised through a grid-search with exponentially growing sequences as suggested by Hsu et al. (2003) and through tenfold cross-validation with ten repetitions. As specimens belonging to different clades were observed in unequal numbers (class imbalance), we optimised the parameters based on Cohen’s κ a multiclass performance metric accounting for class imbalance (Landis and Koch, 1977). We considered κ < 0.2 a *slight*, κ between 0.2 and 0.4 as *fair*, κ between 0.4 and 0.6 as *moderate*, κ between 0.6 and 0.8 as *substantial*, and κ above 0.8 as *almost perfect* agreement (Landis and Koch, 1977). Prediction provided by the optimised algorithm were compared to the clade affiliations provided the combined molecular-morphological parsimony analyses and the literature survey. Finally, we inferred the variable importance for SVM predictions through a pairwise receiver operating characteristic (ROC) curve analysis as implemented in the function *filtervarImp*. We considered an area under the curve (AUC) between below 0.7 *poor*, between 0.7 and 0.8 *acceptable*, between 0.8 and 0.9 *excellent*, and above 0.9 *outstanding*.

To further assess the systematic informativeness of the morphometrics, we investigated the amount of possible redundancies by identifying clusters of multicollinear morphometric measurements. We used a Pearson pairwise correlation matrix (Dormann et al., 2013) and the *ward.d2* clustering algorithm (Murtagh and Legendre, 2014) to detect clustered variables in the *R* package *ComplexHeatmap* v2.8.0 (Gu et al., 2016). If the absolute values of the Pearson correlation coefficients (|r|) exceeded 0.7 (Dormann et al., 2013), we considered measurements multicollinear. Hence, we split the variables into clusters along the dendrograms provided by *ward.d2* until all coefficients within the same cluster exceeded this threshold. Heatmaps were plotted using the *R* package *ComplexHeatmap* v2.8.0 (Gu et al., 2016).

### Character evolution of the attachment and reproductive organs and phylogenetic signal

Using the morphometric measurements, we tested for possible patterns of a random walk (H_1_), stabilising selection (H_2_), and adaptive radiation with a decelerating (H_3_) and accelerating divergence of characters (H_4_) on the evolution of the attachment and reproductive organs. We also tested if species from the East African lakes followed a different evolutionary regime (H_5_) as a result of the multiple host radiations in this region. To detect these patterns, we employed multivariate phylogenetic comparative methods as implemented in the *R* package *mvmorph* v1.1.4 (Clavel et al., 2015) to account for potential interactions between characters. This software package addresses the sensitivity of previous multivariate approaches (e.g. Khabbazian et al., 2016; Goolsby et al., 2017) to trait covariation, data orientation, and trait dimensions (Adams and Collyer, 2018).

We fitted a range of multivariate models of continuous character evolution on a single sample of 100 randomly selected tree topologies drawn from the post–burn-in phase of the BI analysis of the molecular markers. Morphometric measurements were averaged by species. Furthermore, we excluded all but one specimen per species from the phylogenetic trees to avoid that specimens of the same species are assigned identical values, which might otherwise artificially decrease the estimated variability of the measurements per taxon affecting model performance. Specimens included in the subset trees, were chosen at random and are highlighted in Appendix 3. The tested models include:

- The Brownian motion model (BM) (Felsenstein, 1973) simulates a ‘random walk’, i.e. genetic drift or randomly fluctuating selection (see Hansen and Martins, 1996) (H_1_). In this case, the parasite morphology would not be affected by strong selection pressures and would randomly diverge over time.
- The Ornstein-Uhlenbeck model (OU) (Butler and King, 2004) approximates a character evolution under stabilising selection (H_2_). Here, the parasite morphology experiences a selection pressure towards a selective optimum due to a high host-dependence.
- The Early-Burst (EB) (Harmon et al., 2010) or accelerate-decelerate model (ACDC) (Blomberg et al. 2003) models an early rapid evolution of characters followed by a slow-down such as might occur during adaptive radiation events (H_3_). Here, the parasite morphology has rapidly diverged possibly mirroring the explosive speciation events reported for some of the host lineages.
- The Late-Burst model (LB), a variation of the ACDC model with an accelerating divergence of characters (Blomberg et al., 2003) (H_4_), simulates evolution as expected under a recent, ongoing radiation event. The speciation of the parasites would mirror the host radiations but would still be ongoing.
- The multi-rate Brownian motion model (BMM) represents two different multi-selective and multi-rate Brownian motion regimes for East African parasite lineages and other lineages (O’Meara et al., 2006). Here, the parasite morphology would have undergone an evolutionary rate shift because species infecting the host radiations in Eastern African have developed differently than their congeners in the rest of Africa. The regimes were defined using the function *paintSubTree* in the *R* package *phytools* v0.7-80 (Revell, 2012).

We fitted all models using the *mvgls* function (Clavel et al., 2019) in *mvmorph* as recommended for independently evolving measurements with a natural orientation. Therefore, we used a restricted maximum likelihood (REML) estimation with a leave-one-out cross-validation of the penalized log-likelihood (PL-LOOCV), an estimated within-species variation, an archetypal ridge estimator, and a diagonal unequal variance target matrix (Clavel et al., 2019). To run the LB model with *mvgls*, we set the upper bound of the EB search algorithm to 10. For the BMM model, we defined a separate regime for the *EAR* clade (Fig. 2), which encompasses species infecting cichlids of the East African radiation (Fig. 3b). As traditional information criteria such as the Akaike information criterion (AIC) are not applicable to penalised likelihoods (Clavel et al., 2019), we assessed the model performance with the extended information criterion (EIC) (Ishiguro et al., 1997; Kitagawa and Konishi, 2010) using the function *EIC* in *mvmorph* with 100 bootstrap replicates. Smaller EIC values were considered indicative of a better model performance. To test for phylogenetic signal in the datasets, we also inferred Pagel’s λ (Pagel, 1999) through the *lambda* option in *mvgls*.

**Figure 3.**
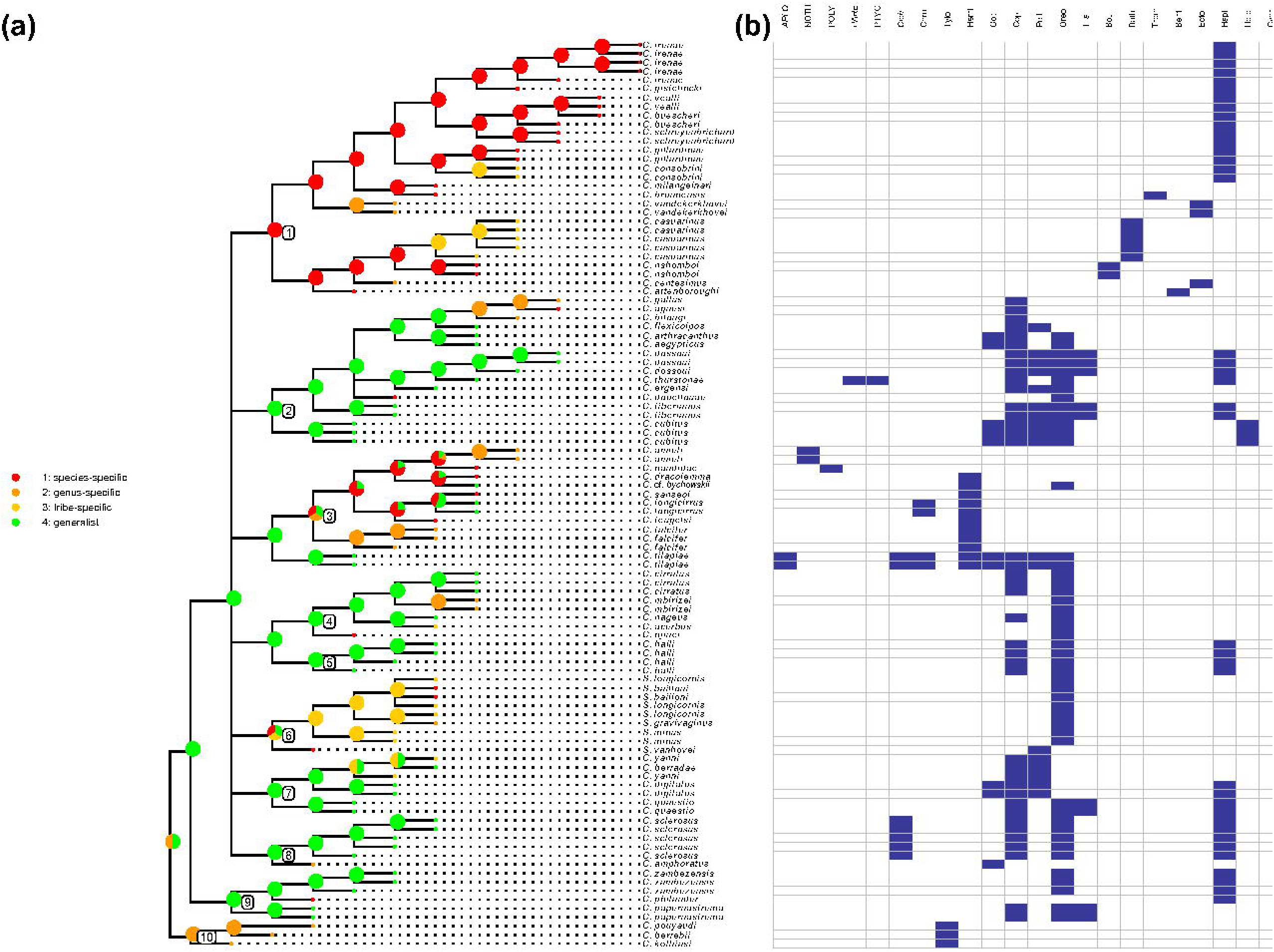
Host repertoire of species of *Cichlidogyrus.* (a) Character map of index of specificity according to Mendlová and Šimková (2014). (b) Host range matrix of the species by tribe or subfamily of cichlid hosts and family of non-cichlid hosts (recent anthropogenic host range expansions italicised). APLO, Aplocheilidae; NOTH, Nothobranchiidae; POLY, Polycentridae; Pare, Paretroplinae; Ptyc, Ptychochrominae; Cich, Cichlasomatini; Chro, Chromidotilapiini; Tylo, Tylochromini; Hemi, Hemichromini; Gobi, Gobiocichlini; Copt, Coptodonini; Hete, Heterotilapiini, PelL, Pelmatolapiini; Oreo, Oreochromini; Tila, Tilapiini; Boul, Boulengerochromini; Bath, Bathybatini; Trem, Trematocarini; Bent, Benthochromini; Cypr, Cyprichromini; Ecto, Ectodini; Hapl, Haplochromini.

Finally, we modelled the character evolution of discrete characters proposed in previous studies (see *Morphological and host range data collection*) under a univariate Pagel’s λ model (Pagel, 1999) using the *fitDiscrete* function in the *R* package *geiger* v2.0 (Pennell et al., 2014) to assess the effects of data discretisation. We applied this function to the same set of 100 randomly selected BI subset trees fitted to the morphometric data to account for phylogenetic uncertainty. Model performance was assessed through the sample size–corrected Akaike information criterion in relation to a white noise model (ΔAICc) that assumes absolute phylogenetic independence (Pennell et al., 2014). We also produced character maps for the discrete morphological characters (HC, VBS, AS) and the equivalent sets of continuous characters using *asr_max_parsimony*, *anc.ML*, *ggplot2*, and *ggtree* as mentioned above.

## Results

### Phylogenetic analyses: Molecular data

We generated 60 sequences for 33 species and 64 specimens of *Cichlidogyrus* and *Scutogyrus* including 47 28S, 14 combined 18S and ITS1 rDNA, and five COI mtDNA sequences. For 21 of these species, these sequences are the first to be published representing a 57% increase in taxon coverage to 42% of all currently described species, 45% of described species of *Cichlidogyrus* and 71% of *Scutogyrus*. The alignments include 93 sequences/820 base pairs for 28S rDNA, 54 sequences/481 base pairs for 18S rDNA, 57 sequences/383 base pairs for ITS rDNA, and 19 sequences/489 base pairs for COI mtDNA. The combined data set includes sequences of 103 specimens belonging to 58 species (see Appendix 3), and has a length of 2173 base pairs following the removal of poorly aligned positions and divergent regions (missing sequence data were treated as gaps). We deposited sequences generated for this study in GenBank (XXXXXX – XXXXXX). All sequences are listed in Appendix 3 including the respective GenBank accession numbers, the host species, and the sampling locations. The concatenated alignments and MP, BI, and ML tree topologies can be accessed at Treebase (www.treebase.org, accession number: XXXXX). Substitution models used for the different partitions are provided in Appendix 4.

The BLK weightings scheme produced the most stable tree topologies for the molecular markers (distortion coefficients and SPR distances approximately 1.000) for all k values. For the combined parsimony analysis, k = 48 produced the most stable tree topology (distortion coefficient: 0.901; SPR distance: 0.743 with 46 SPR moves). Thus, the final hypotheses (molecular and combined trees) were inferred from the strict consensus the trees produced under these parameters (BLK; k = 48) (Fig. 2a; Fig. 4). Model selection for BI and ML analyses resulted in the merging of all COI codon positions. The BI and ML concatenated tree topologies were congruent with the final MP consensus tree (Kendall’s W = 0.76, χ² = 12066, p < 0.01). Thus, we display support values of the ML analysis (UFBoot) alongside the posterior probabilities in the BI phylogram (Fig. 2b).

**Figure 4.**
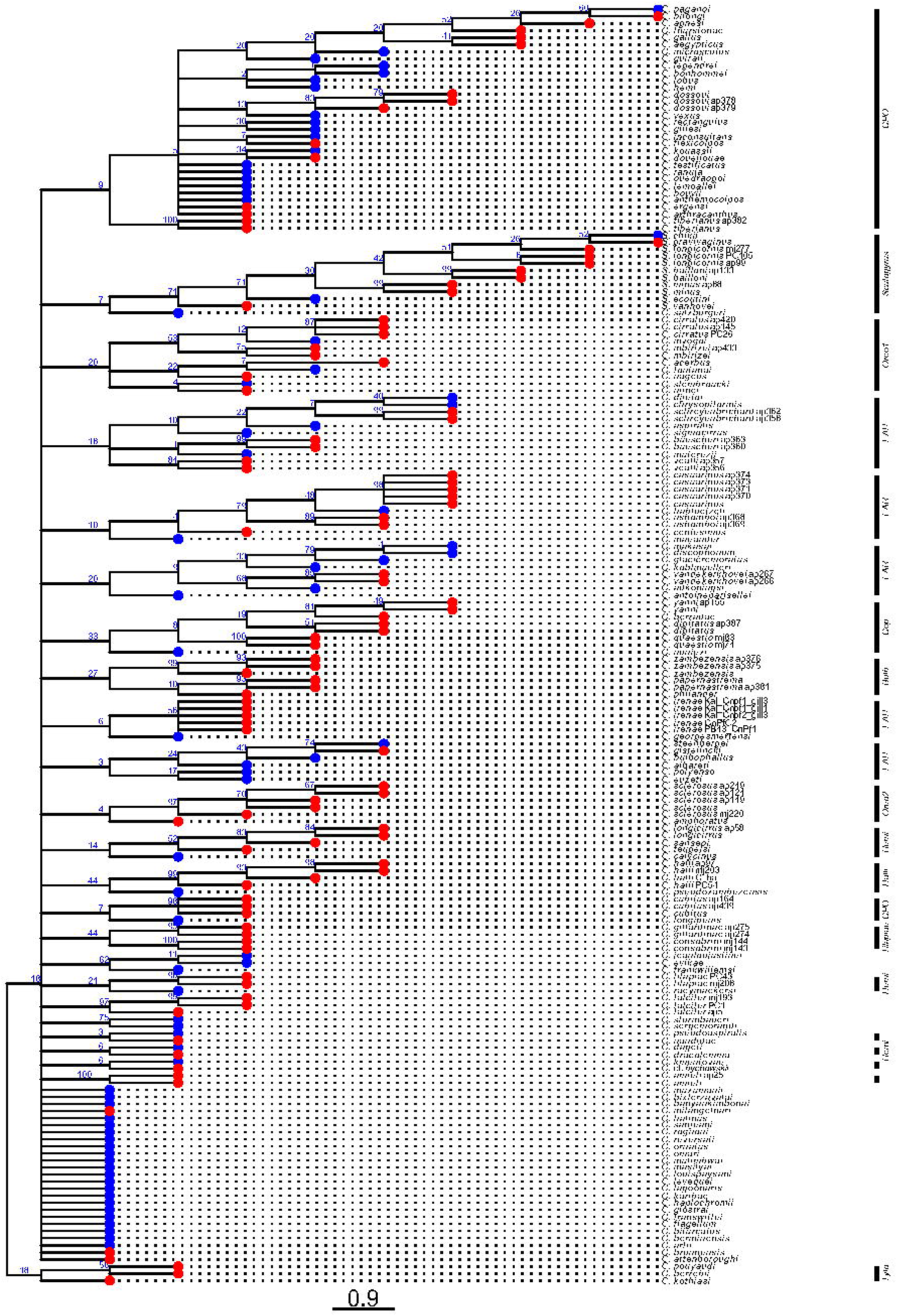
Consensus tree of phylogenetic analysis under maximum parsimony and extended implied weighting (k = 48, BLK weighting scheme) combining molecular and morphometric data; red tip points indicate species with and blue without DNA sequence data; node support values constitute GC values inferred from symmetric resampling. Compared to the molecular tree (Fig. 2a), this tree is less resolved and less supported.

We found 11 well-supported clades within *Cichlidogyrus* based on the MP analysis of the molecular dataset (Fig. 2a). These clades were also supported the BI and ML analyses suggesting a high stability of the phylogenies produced by the molecular dataset provided here. The clades and the their respective node support values (GC value/PP/UFBoot/SH-aLRT) are (Fig. 2a): the *‘EAR’* clade (27/1/99/99) (#1), the *‘CPO’* clade (91/1/100/100) (#2), the *‘Hemi’* clade (94/1/100/100) (#3), the ‘*Tilapiae*’ clade (99/1/100/100) (#4), the *‘Oreo1’* clade (76/1/99/98) (#5), the *‘Halli’* clade (100/1/100/100) (#6), *Scutogyrus* (86/0.99/*/90) (#7), the *‘Cop’* clade (48/1/100/100/*) (#8), the *‘Bulb’* clade (92/1/100/100) (#9), the *‘Oreo2’* clade (81/1/97/94) (#10), and the basal *‘Tylo’* clade (100/1/100/100) (#11) (see *Characterisation of species groups* for details of morphology and host range). Beyond the well-supported monophyly of all species excluding the *Tylo* group (100/1/100/100), the MP tree shows the *Bulb* clade as sister group to all the other clades (51/*/*/*). Furthermore, *C. tilapiae* and *C. halli* are positioned as sister clades of *Hemi* and *Oreo1* respectively albeit with moderate support (GC values of 11 and 9). The BI and MP analyses also provides partial support for two main lineages containing *Oreo1*, *Cop*, *Scutogyrus*, and *Hemi* (0.99/*/*/*) or *Cop*, *Scutogyrus*, and *Hemi* (*/*/87/*). Both main lineages would also include *C. tilapiae* Paperna, 1960.

### Literature survey and character mapping

For a more extensive classification of species of *Cichlidogyrus* into the proposed species groups, we developed four new discrete characters for the reproductive organs: the shape of the penis, its diameter, the shape of the accessory piece, and the shape of the sclerotised vagina (for the respective character states, see Appendix 2). We mapped all morphometric and the newly proposed discrete characters on the molecular MP phylogeny (Fig. 5). Out of the 11 species groups (Fig. 2a), we found shared morphological characteristics in eight. Species of the *Oreo2* and the *EAR* groups shared no apparent features in their attachment and reproductive organs with species of the same group apart from the characteristics mentioned in the genus diagnosis (Pariselle and Euzet, 2009). We were able to affiliate all but 12 species without available DNA sequences to these clades (Appendix 5) based on character states shared with species included in the molecular phylogeny. Some morphological structures were particularly relevant as synapomorphies of the species groups (Appendix 6; Appendix 7; also Fig. 5a and Fig. 6). Based on the number of species groups defined by a structure, the male copulatory organ (MCO) was the most distinctive structure as phenotypes were specific to eight species groups (and several subgroups) (Fig. 5b; Fig. 6; Appendix 6). The first hook pair (U1) was enlarged in four species groups (Fig. 6) and one subgroup of the morphologically diverse *EAR* group. The length of the auricles of the dorsal bar (DB h) was distinct in two species groups (Fig. 6) and one subgroup of the *EAR* group. Hook pairs 3–7 were enlarge in two groups (Fig. 6). For the detailed morphological characterisations of all mentioned groups and a discussion on the host and geographical ranges, see *Characterisation of species groups* as well as several character maps (Fig. 5) and an overview of the morphological features (Fig. 6) (full species names are cited in Appendix 5 and reported alongside host names; for abbreviations of morphometric measures, see Fig. 1c).

**Figure 5.**
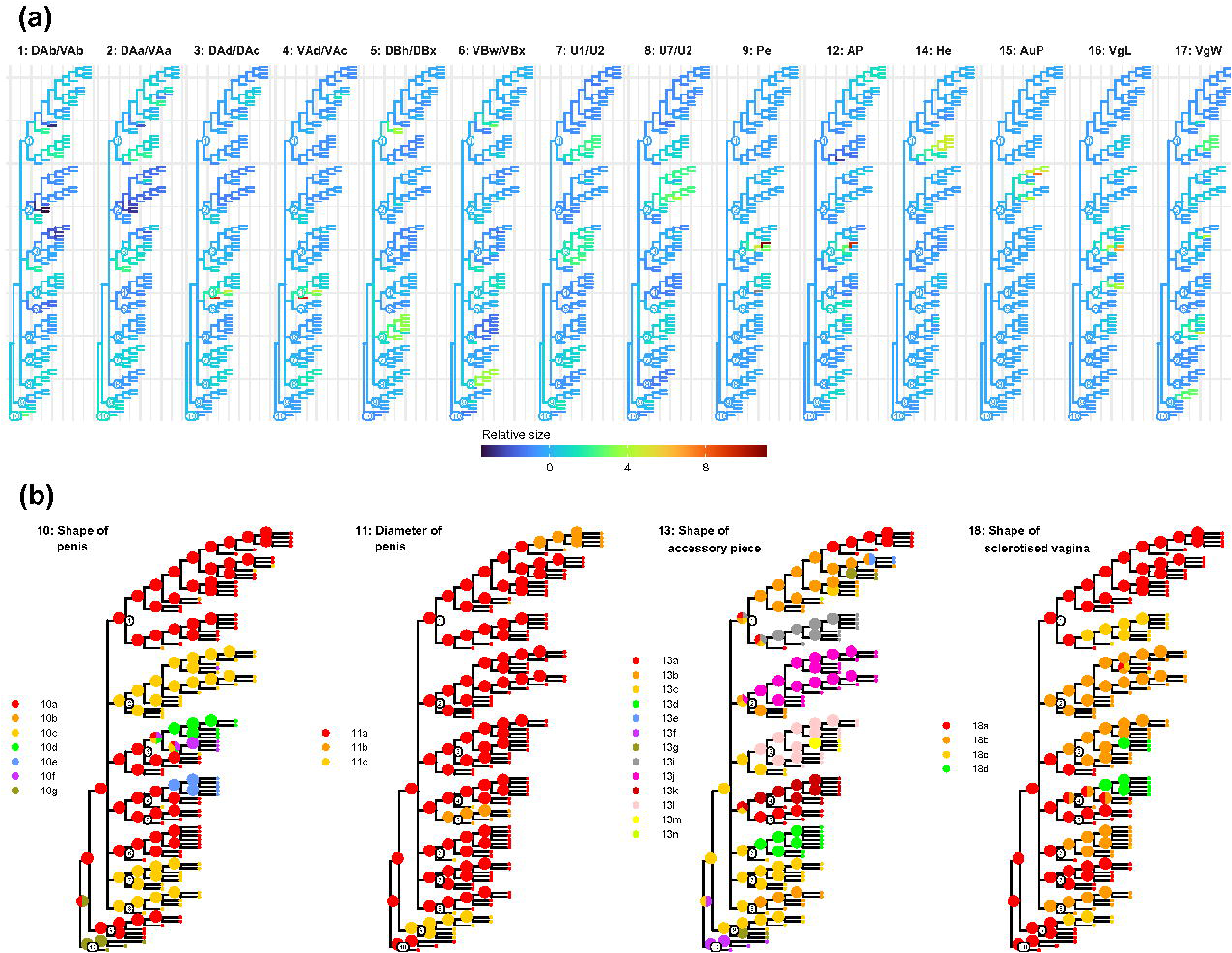
Character maps of morphological characters. (a) Scaled means of morphological continuous characters with abbreviations referring to measurements of sclerotised structures in the attachment and reproductive organ used to characterise species belonging to *Cichlidogyrus* (see Fig. 1). (b) Character maps of newly proposed discrete characters for the reproductive organs; indices in legend refer to character states suggested in Appendix 2. For details on all characters, see numbers in Appendix 2. Numbers on nodes refer to clades in Fig. 2a.

**Figure 6.**
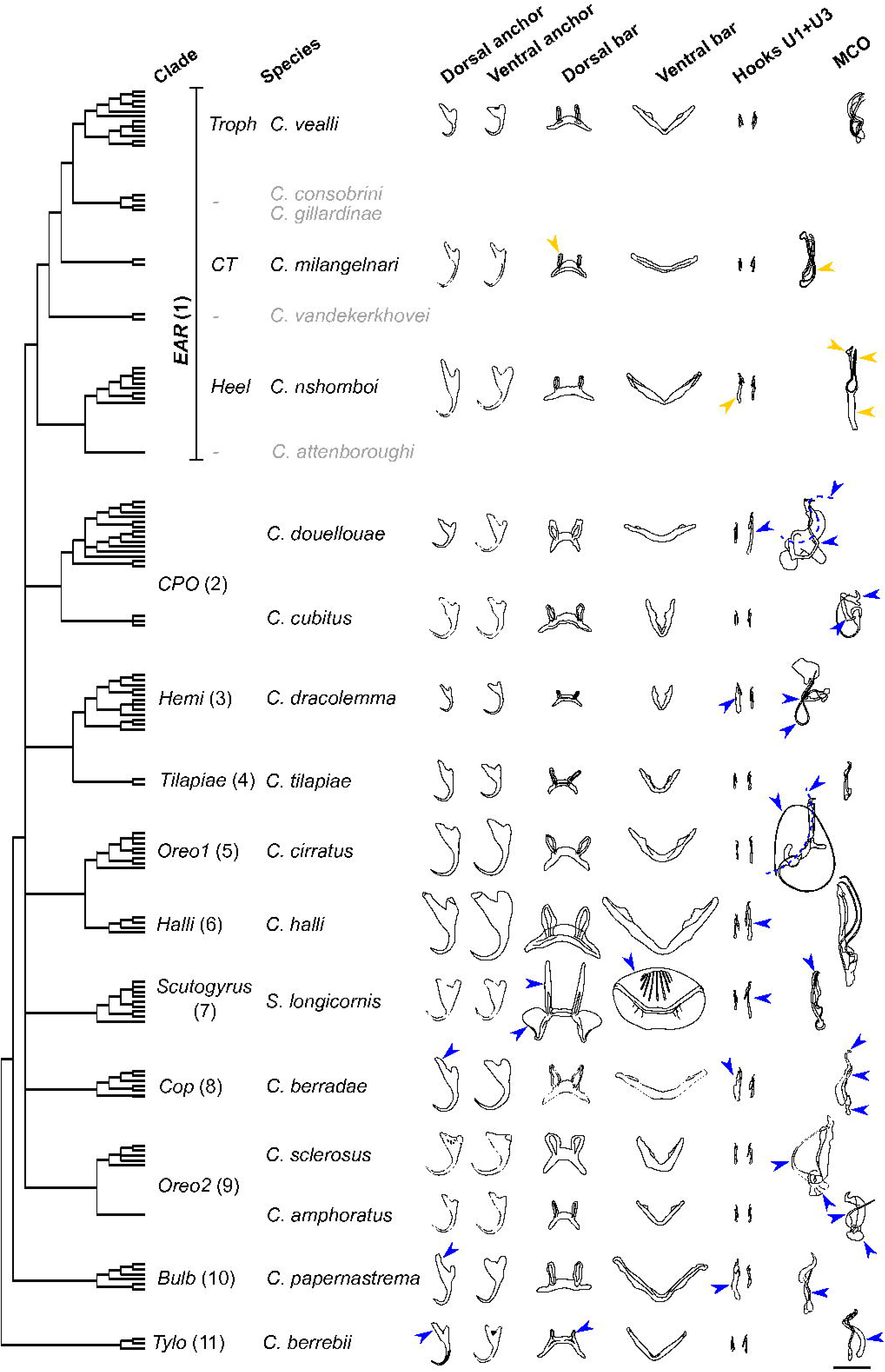
Morphology of attachment and reproductive organs of representative species belonging to proposed species groups and subgroups (Fig. 2) of *Cichlidogyrus* (incl. *Scutogyrus*) including cladogram and illustrations of sclerotised structures of attachment organ (anchors, bars, hooks) and the MCO of selected species (vagina morphology not shown as little unifying or contrasting morphological patterns were detected between groups, see “Characterisation of species groups”). Arrows indicate key features of each species group (blue) and subgroup of the *EAR* group (yellow), dashed lines indicate typical shapes. Species of the *EAR* group that are not displayed are labelled in grey. Scale = 30 µm.

### Characterisation of species groups

In the following section we summarise the results of the literature survey. All continuous and discrete characters and their states are explained in Appendix 2. Ancestral character states for the different species groups (and subgroups) are listed in Appendix 6 and Appendix 7. The following characterisations also discuss morphological differences to species groups with similar characteristics. We also refer to publications that have previously mentioned or discussed shared characteristics of these species groups. For 12 species, we found no morphological similarities that could place them near any of the species groups characterised above.

1. *EAR: Species infecting cichlids from the East African Radiation*. The *EAR* species group comprises a large heterogeneous group of species that infect cichlids from East African lineages, i.e. the East African Radiation (Schedel et al., 2019). The *EAR* clade consists of multiple potential subgroups based on morphological features (Fig. 6) and node support values from the phylogenetic analyses. The subgroups of the *EAR* group include species with a long heel infecting non-haplochromine cichlids such as bathybatine, ectodine, boulengerochromine, and possibly cyphotilapiine cichlids (*Heel*), species infecting cyprichromine and trematocarine and potentially eretmodine cichlids (*CT*), and species infecting tropheine cichlids (*Troph*) (see Fig. 6). The morphology of the attachment and reproductive organs vary substantially between and within the subgroups (Fig. 5b). Hence, we observed no characteristic features for the whole group with rather general and simple ancestral features (e.g. small marginal hooks; straight, simple penis; no sclerotised vagina, see Appendix 6 and Appendix 7). However, some subgroups display the following shared morphological features.

- *Troph* (90/*/99/80). These species share few similarities and were placed in a subgroup due to the host species, which all belong to the tropheine radiation. Species without available DNA sequences, infecting tropheines, and sharing morphological features with species in this group, were also added to this subgroup. Dorsal anchor unlike ventral with elongated inner root (except for *C. antoineparisellei*) and deeper indentation between the roots. However, indentation generally shallow, sometimes with fused roots (see parasites of *Interochromis loocki* including *C. antoineparisellei*, *C. buescheri*, *C. schreyenbrichardorum*, *C. vealli* on the ventral and *C. antoineparisellei* and *C. buescheri* on the dorsal anchor) (Fig. 6). Dorsal bar with well-developed but not elongated auricles. Hooks generally small but can be slightly more developed (e.g. *C. irenae, C. gistelincki*, *C. masilyai, C. salzburgeri*). The MCO consists of penis and accessory piece, both variably shaped. Penis thin and arched (*C. antoineparisellei*, *C. buescheri*) or looped (*C. schreyenbrichardorum*) or slightly broadened (*C. vealli*) (Fig. 5b; Fig. 6) for parasite of *Interochromis loocki* or thin and slightly sinuous for *C. gistelincki*. Penis broadened to wide and short (reminiscent of *C. halli*) for other species. Accessory piece simple, elongated with few remarkable structures, sometimes with cap-like (*C. gistelincki*), forked (*C. antoineparisellei, C. buescheri, C. masilyai, C. salzburgeri*), or hook-shaped (*C. raeymaekersi*) distal end.
- *Heel* (97/1/99/97). Dorsal anchor unlike ventral with elongated inner root. Dorsal bar with short auricles. Well-developed first hook pair. Male copulatory organ with short penis sometimes with spirally screw thread–like thickened wall (*C. casuarinus, C. centesimus*, *C. nshomboi*), characteristic elongated heel (Fig. 5a) of one third to twice the length of the penis, and reduced filiform or sometimes missing (only *C. centesimus*) accessory piece (Fig. 5b; Fig. 6). The species of the *Heel* subgroup have previously been grouped together (Muterezi Bukinga et al., 2012), most recently by Rahmouni et al. (2018b).
- *CT* (97/1/100/90). Dorsal and ventral anchors similar in shape but can vary in size (see *C. brunnensis*) with deep to shallow (*C. brunnensis*) indentation. Dorsal bar with short auricles. Hooks generally short. Male copulatory organ consists of penis and accessory piece and sometimes a marked heel (*C. brunnensis*). Penis medium-length, broadened, and mostly straight (*C. brunnensis, C. evikae, C. jeanloujustinei*, *C. milangelnari*, *C. sturmbaueri*) with thickened wall. Accessory piece with bifurcate distal end (*C. brunnensis*), or in two-parts (*C. evikae, C. jeanloujustinei, C. milangelnari*). Similarities of some species of this subgroup with species infecting cyphotilapiine and ectodine cichlids with elongated auricles (*C. adkoningsi*, *C. discophonum*, *C. glacicremoratus*, *C. koblmuelleri*, *C. makasai*, *C. vandekerkhovei*) or elongated marginal hooks (*C. rectangulus*, *C. sturmbaueri*) have previously been discussed (Rahmouni et al., 2017, 2018) but the phylogenetic analysis provided no support for a monophyletic group including species of the *CT* subgroup and *C. vandekerkhovei* (Fig. 2a).
- *Other species*. The remaining species of the *EAR* species group shared no evident features with any of the proposed subgroups. *Cichlidogyrus attenboroughi* appears to be the sister taxon to the *Heel* group with moderate support (56/0.99/*/*) but shares no notable characteristics with these species. Several species infecting cyphotilapiine and ectodine cichlids display a broadened (*C. glacicremoratus*, *C. koblmuelleri*) or straight (*C. adkoningsi*, *C. rectangulus, C. sturmbaueri*) penis with an accessory piece in two-parts (*C. glacicremoratus*, *C. koblmuelleri*) or with a bifurcated distal end (*C. makasai*, *C. rectangulus, C. sturmbaueri*, *C. vandekerkhovei*) similar to species of the *CT* group. Yet many of these species also display elongated auricles at the dorsal bar (*C. adkoningsi*, *C. discophonum*, *C. glacicremoratus*, *C. koblmuelleri*, *C. makasai*) similar to *C. vandekerkhovei*, which appears to be unrelated to species of the *CT* group (Fig. 2a) but also displays a bifurcate distal end. According to the phylogenetic analyses (Fig. 2a), *C. consobrini* and *C. gillardinae* form a monophyletic group. However, we found few shared characteristics except for a simple tubular penis with a small heel and a simple accessory piece. These characteristics might place these two species close to *C. haplochromii* and *C. longipenis*, both of which have been reported from haplochromine cichlids in Lake Victoria (Pariselle and Euzet, 2009). Yet these simple features might also reflect ancestral character states of the *EAR* group (Appendix 6; Appendix 7).
2. *CPO: Species infecting coptodonine, pelmatolapiine, oreochromine and other cichlids*. Dorsal anchor with inner root longer than outer root and V-shaped indentation between the roots but inner root of *C. arthracanthus* considerably longer than of other species. Roots of *C. cubitus* and *C. louipaysani* fused. Ventral anchor similar shape to dorsal anchor but slightly larger. However, roots of ventral anchor of *C. tiberianus* fused. Dorsal bar slightly arched with large medium-sized to large auricles. Ventral bar always with membranous extensions. First hook pair (U1) small and hook pairs 3–7 (U3–7) very long, i.e. more than triple the size of first pair (Fig. 5a; Fig. 6) but U1 of *C. arthracanthus* large and U3–7 of *C. cubitus* and *C. louipaysani* short (Fig. 6). The MCO consists of long, arched, or sometimes spiralled (*C. arthracanthus*), tubular penis with a well-marked irregularly shaped heel, and a massive, roughly S-shaped accessory piece (Fig. 5b; Fig. 6) that is frequently connected to the heel. The accessory piece has an extension or thickening at the first turn in the proximal half and frequently displays a folded back (*C. paganoi, C. vexus*), straight and pointy (*C. guirali*), or hook-like (*C. bilongi, C. douellouae, C. ergensi, C. gallus, C. legendrei, C. microscutus*) distal end, or sometimes additional terminations resulting in a furcate ending with two (*C. aegypticus, C. agnesi, C. anthemocolpos, C. bouvii, C. cubitus, C. flexicolpos, C. lemoallei, C. louipaysani, C. ouedraogoi, C. testificatus, C. thurstonae, C. tiberianus*) or three (*C. bonhommei, C. hemi, C. kouassii*) digitations. However, the first turn is never V-shaped or knee-like as in species of the *Hemi* clade and the hook-shaped termination is never sickle-like such as in species of the *Cop* clade. Several species display an auxiliary plate (AuP) close to the distal end of the MCO (Fig. 5a) including *C. aegypticus, C. agnesi, C. bilongi, C. gallus, C. guirali* (two pieces), *C. microscutus, C. paganoi,* and *C. thurstonae*. A sclerotised vagina is mostly present but has not been reported for *C. arthracanthus* (Paperna, 1960; Ergens, 1981). Other species with more developed hook pairs 3–7 include *Cichlidogyrus bulbophallus, C. flagellum, C. lobus,* and *C. ranula,* some species of the *Hemi* clade, and all species of the *Heel* subgroup of the *EAR* clade. *Cichlidogyrus bulbophallus* and species of the *Hemi* clade and the *Heel* subgroup have been likened to *C. arthracanthus* because of the additional well-developed first hook pair (U1) (Geraerts et al., 2020). However, all of these species differ from the confirmed clade members as follows.

- The dorsal and ventral anchors differ in shape and size but are similar for species of the *CPO* clade.
- The penis is never long, arched, and tubular but short (*Heel*), bulbous (*C. bulbophallus, C, flagellum, C. ranula*), or draws a loop (Fig. 6) or long spiral (*Hemi*).
- The accessory piece is not complex and S-shaped with an extension in the proximal half but mostly reduced (*Heel*), simple (*C. bulbophallus, C, flagellum, C. ranula*), or has a sharp knee-like bend (*Hemi*). Previous phylogenetic studies (Mendlová et al., 2012; Messu Mandeng et al., 2015) have placed *C. cubitus* alongside species of the *CPO* clade with long marginal hooks (“*tiberianus*” group, see Pouyaud et al., 2006) but morphological similarities were not further discussed.
3. *Hemi: Species infecting hemichromine cichlids and non-cichlid fishes*. Dorsal anchor with elongated narrow inner root and rounded outer root. Ventral anchor slightly larger with short and robust inner root. Dorsal bar slightly arched with short auricles. Ventral bar arched with small membranous extensions. First hook pair (U1) large and well-developed (Fig. 5a, Fig. 6) except for *C. amieti*. Hook pairs 3–7 medium-sized to large with bases more developed than for second hook pair. The MCO consists of thin and long penis that frequently forms loops (*C. amieti, C.* cf. *bychowskii*, *C. dracolemma, C. inconsultans, C. kmentovae, C. nandidae*) (Fig. 6) or spirals with one (*C. calycinus, C. teugelsi*) to multiple turns (*C. euzeti, C. longicirrus, C. polyenso, C sanseoi*) and re-joins the accessory piece in its distal portion, and an accessory piece of two distinct portions. These portions can be shaped like a V or a simple spiral with an expanded knee-like bend (*C. amieti, C. calycinus, C.* cf. *bychowskii*, *C. dageti, C. dionchus, C. dracolemma, C. falcifer, C. kmentovae, C. nandidae, C.teugelsi*) (see Dossou and Birgi, 1984; Pariselle and Euzet, 2004), or a large spiral followed by a non-spiralled distal portion (*C. euzeti, C longicirrus, C. polyenso, C. sanseoi*). In the presence of the knee-like bend a heel is visible. The sclerotised vagina is long, sinuous (except for *C. dageti, C. falcifer, C. kmentovae*) or spiralled (*C. longicirrus, C. sanseoi*). Species of the *Cop* and *Tylo* groups share some characteristics with the species of the *Hemi* group but have either a substantially shorter penis or do not display the characteristic knee-like bend in the accessory piece. Previous studies have grouped species of the *Hemi* species group together based on phylogenetic (Mendlová et al., 2012; Řehulková et al., 2013) and morphological (Dossou and Birgi, 1984; Pariselle and Euzet, 2004; Jorissen et al., 2018a) analyses.
4. *Tilapiae: The species complex of Cichlidogyrus tilapiae*. For a detailed characterisation, see species descriptions and characterisations by Paperna (1960) and Douëllou (1993). *C. tilapiae* have both been hypothesised to form a species complex (Pouyaud et al., 2006).
5. *Oreo1: Species infecting oreochromine and coptodonine cichlids*. Anchor pairs similar in shape and size but dorsal anchors somewhat smaller. Anchors with short and broadened outer root and long inner root. Dorsal bar arched with two broad and large auricles. Ventral bar with thin mid-portion, and thickened towards the ends includes triangular membranous extensions. Hooks small with small shaft. The MCO consists of long tubular (*C. cirratus, C. giostrai, C. mbirizei, C. mvogoi, C. ornatus*) (Fig. 6) or short and thickened (*C. acerbus, C. lagoonaris, C. nageus, C. njinei, C. slembroucki*) penis (Fig. 5b) sometimes with bulbous extension at the base of the tube (*C. njinei, C. ornatus, C. giostrai*) and complicated roughly C-shaped (Fig. 6) accessory piece often with finger or hook-shaped outgrowths and marked heel. Proximal end of accessory piece attached to penis bulb and distal end holds the distal portion or a mid-portion of the penis. Vagina simple (*C. acerbus, C. lagoonaris, C. slembroucki*), sinuous (*C. giostrai, C. njinei, C. ornatus*), or coiled (*C. cirratus, C. mbirizei, C. mvogoi*) tube and only slightly sclerotised.
6. *Halli*: *The species complex of Cichlidogyrus halli*. For detailed characterisation see original species description by Price and Kirk (1967). *Cichlidogyrus halli* has both been hypothesised to be a species complex (Douëllou, 1993; Jorissen et al., 2018b).
7. *Scutogyrus: Species infecting oreochromine and pelmatolapiine cichlids*. This group has been diagnosed as the separate genus *Scutogyrus* including a detailed morphological characterisation (Pariselle and Euzet, 1995). Concerning the haptor morphology, species of *Scutogyrus* display well-developed hook pairs 3–7, elongated auricles at the dorsal bar (Fig. 5a; Fig. 6) and a ventral bar that supports a large plate (Fig. 6). The accessory piece presents a distal flap (Fig. 5b; Fig. 6).
8. *Cop: Species infecting coptodonine cichlids*. Anchor pairs dissimilar in shape. Dorsal anchor with V-shaped indentation between the roots and an elongated inner root at least four times longer (Fig. 5a) than the outer root. Ventral anchor with shallower indentation, and relatively short roots including a considerably shorter inner root. Dorsal bar thick with large and broad auricles. Ventral bar V-shaped and truncated at the ends and thinner towards the mid-portion, displays membranous extensions. First hook pair (U1) well-developed (Fig. 5a, Fig. 6). Hook pairs 3–7 (U3–7) small. The MCO consists of short tubular penis with a marked heel and an accessory piece with a sickle-like terminal hook connected to the basal bulb of the penis. No sclerotised vagina has been observed for species belonging to this group except for *C. digitatus* in the original description (Dossou 1982). However, none of the more recent reports of species of the *Cop* clade (Pariselle and Euzet, 1996, 2003) remark on this report. Previous studies have grouped species of the *Hemi* clade, e.g. *C. dionchus* (Pariselle and Euzet, 2003), and the *Tylo* clade, e.g. *C. kothiasi* (Pariselle et al., 2013), with the species of the *Cop* clade because of the similar morphology of the hook pairs, i.e. large U1 and small U3–7. However, none of these species displays an accessory piece with a sickle-like terminal hook. Species of the *Cop* clade have previously been grouped together (Pariselle and Euzet, 2003; Jorissen et al., 2018b) but without *C. berminensis*.
9. *Oreo2: Two species infecting oreochromine and coptodonine cichlids*. Both species display simple features such as similar anchor pairs and small hook pairs. Most other features differ considerably between the species. Dorsal and ventral anchors are similar in size and shape, the roots are variable in length, i.e. a long inner root in *C. amphoratus* (see drawing Pariselle and Euzet, 1996) or barely to no distinct roots in *C. sclerosus* (Douëllou, 1993). Dorsal bar X-shaped with two auricles of variable shape and length. Ventral bar variable in shape and size with membranous extensions (Mendlová et al., 2012). Hooks small with developed but small shaft except for the second pair (Douëllou, 1993; Pariselle and Euzet, 1996). The MCO consists of arched, tubular penis (Fig. 5b) with large heel (‘subspherical’ in *C. amphoratus* or shaped as a ‘serrated plate’ in *C. sclerosus*), and associated relatively simple accessory piece (Douëllou, 1993; Pariselle and Euzet, 1996) (Fig 5b). Sclerotised vagina present. *Cichlidogyrus amphoratus* presents a swollen middle portion of the penis reminiscent of species of the *Bulb* clade (Pariselle and Euzet, 1996). However, in the latter group, the dorsal anchors have long inner roots and the first hook pair is well developed (Fig. 6). The close relationship of these species has previously been reported but was not further discussed (see Messu Mandeng et al., 2015).
10. *Bulb: Species from Southern Africa with a bulbous penis*. Dorsal and ventral anchors dissimilar. Dorsal anchors with deep indentation between short outer root and a long inner root (Fig. 6). Dorsal bar short and stout with auricles medium-sized to large (average > 20 µm for *C. zambezensis*). First hook pair (U1) well-developed (Fig. 5a; Fig. 6) albeit somewhat less in *C. zambezensis*. Hook pairs 3–7 small to slightly developed. Male copulatory organ consists of a broad penis with a swollen, bulbous middle portion, a small heel (Fig. 6), and a sharp tube-like (*C. philander*) or filiform (*C. maeander, C. papernastrema*, *C. pseudozambezensis, C. zambezensis*) termination, and a simple accessory piece of similar length, which is articulated to the base of the penis bulb (Price et al., 1969; Douëllou, 1993; Jorissen et al., 2018b). *Cichlidogyrus amphoratus*, *C. bulbophallus*, *C. flagellum*, *C. giostrai*, *C. karibae*, *C. lagoonaris*, *C. njinei*, *C. ornatus*, and *C. sanjeani* also present a swollen portion in the penis but differ from the species of the group as follows.

- *C. amphoratus* has similar dorsal and ventral anchors with no well-developed inner root of the dorsal anchor (Fig. 6).
- The first hooks (U1) of C. amphoratus (Fig. 6), C. flagellum, C. giostrai, C. karibae, C. lagoonaris, C. njinei, C. ornatus, C. ranula, and C. sanjeani are considerably shorter and less developed.
- The hooks U3–U7 of *C. bulbophallus* and *C. ranula* are considerably longer and resemble the marginal hooks of the species of the *CPO* species group.
- The tubular distal portion of the penis of *C. amphoratus* (Fig. 6)*, C. bulbophallus, C. flagellum, C. giostrai, C. njinei*, and *C. ornatus* exceed the length of the swollen portion of the penis considerably.
- *C. amphoratus, C. giostrai, C. lagoonaris, C. njinei*, *C. ornatus,* and *C. sanjeani* have only been reported from Western Africa not Southern Africa (see Pariselle and Euzet 2009).
- *C. amphoratus* and *C. njinei* belong to the *Oreo2* and *Oreo1* species groups respectively.
11. *Tylo*: *Species infecting tylochromine cichlids.* Dorsal and ventral anchors dissimilar. Roots of anchors frequently fused forming fenestrae (windows), e.g. for *Cichlidogyrus berrebii, C. chrysopiformis, C. dijetoi, C. kothiasi, C. pouyaudi* (Fig. 6), and *C. sergemorandi* (Pariselle and Euzet, 1994; Pariselle et al., 2014; Rahmouni et al., 2018). Dorsal anchors with short outer root and a long inner root, median portion of blade hollow in *C. berrebii, C. pouyaudi* (Pariselle and Euzet, 1994), and *C. sigmocirrus* (Pariselle et al., 2014). Dorsal bar with two generally short auricles fused to the bar (Fig. 6; Fig. 7a) is one of the most distinctive features of species of this group (Pariselle et al., 2014; Rahmouni et al., 2018). The hooks are generally small but the first pair is developed in *C. bixlerzavalai, C. chrysopiformis*, *C. dijetoi*, *C. kothiasi*, and *C. muzumanii* (Fig. 5a). The MCO consists of a coiled tubular penis, a heel, and a flat ribbon-like (*C. berrebii, C. kothiasi, C. pouyaudi, C. sigmocirrus*) (Pariselle and Euzet, 1994; Pariselle et al., 2014), drape-like (*C. chrysopiformis, C. dijetoi, C. mulimbwai, C. omari, C. sergemorandi*) (Muterezi Bukinga et al., 2012; Pariselle et al., 2014; Jorissen et al., 2018a; Rahmouni et al., 2018), or reduced (*C. bixlerzavalai, C. muzumanii*) (Muterezi Bukinga et al., 2012; Jorissen et al., 2018a) accessory piece. Unlike other species of *Cichlidogyrus*, the penis and accessory piece of the MCO are separated at the base but might be connected through a filament in *C. chrysopiformis* and *C. sigmocirrus* (Pariselle et al., 2014). Species of the *Tylo* group have regularly been grouped together since the descriptions by Pariselle and Euzet (1994) with most recent additions provided by Jorissen et al. (2018a) and Rahmouni et al. (2018b).

**Figure 7.**
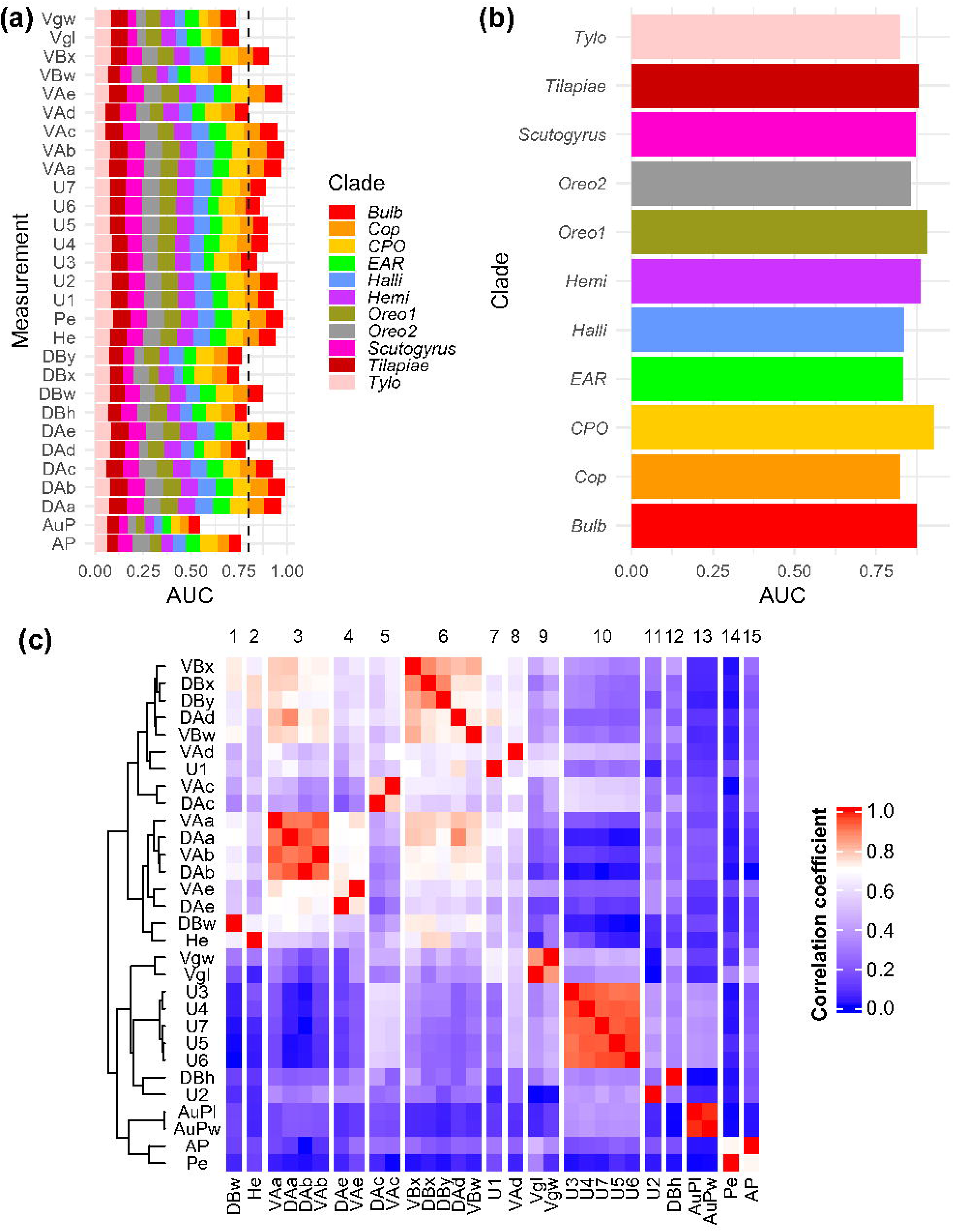
Systematic informativeness of morphometric measurements (a) Variable importance (AUC) according the optimised support vector machines (SVM) classifying specimens belonging to *Cichlidogyrus* into proposed species groups; colours indicate contribution by species group; AUC values > 0.8 are considered excellent discriminators; AUC values between 0.7 and 0.8 are considered acceptable discriminators; AUC values < 0.7 are considered poor discriminators. (b) AUC values clustered by clade showing that the SVMs could distinguish species groups with different accuracy. (c). Cluster analysis of morphometric measurements of attachment and reproductive organs; clusters were detected using the *ward.d2* clustering algorithm. For abbreviation of measurements, see Fig. 1. Species groups were proposed based on phylogenetic analysis (Fig. 2).

### Phylogenetic position of ‘unsequenced’ species and systematic informativeness: literature survey, combined phylogeny, machine learning algorithm, and cluster analysis

The addition of the morphometric data to the parsimony analysis resulted in an overall less resolved and supported tree (Fig. 4). Overall, 30% (24/79) of the ‘unsequenced’ species formed well-supported clades with sequenced species allowing us to affiliate these with the respective clades from the molecular phylogeny. Phylogenetic positions inferred from the combined MP consensus tree were in moderate accordance (κ = 0.48) with the classifications in species groups inferred from the character maps. Meanwhile, the optimised SVMs (C = 2^-7^, σ= 0.5) predicted species group affiliation of the sequenced species with moderate success (κ = 0.68) with 1035 observations included. For the unsequenced species (833 observations), accordance with the parsimony analysis (κ = 0.36) and literature survey (κ = 0.31) was comparatively low.

Variable importances of the optimised SVMs indicated an acceptable to excellent discriminatory power on average (Fig. 7a) with exception of the surface of the auxiliary plate (AuP) in the MCO that performed poorly (mean AUC < 0.6). Collapsing the AUC values by species groups (Fig. 7b), indicates an uneven distribution with some groups such as *CPO* being easier to discriminate against their congeners than others such as *Tylo*. Lastly, based on the Pearson pairwise correlation matrix (Fig. 7c), we detected 15 clusters of multicollinear measurements indicating that almost half of the measurements (14) are multicollinear.

### Character evolution of the attachment and reproductive organs: morphometrics and discretisation

For the attachment organ measurements, we detected a strong phylogenetic signal (λ = 0.91; CI: 0.77–1.00). Brownian Motion (BM), Ornstein-Uhlenbeck (OU), and Early-Burst (EB) models all performed equally (Fig. 4b). The Late Burst (LB) model performed worse. Only the multi-rate BM model (BMM) performed better albeit with a slight overlap in EIC estimates. For the reproductive organ measurements, the phylogenetic signal was considerably weaker (λ = 0.73; CI: 0.43–1.00) and no model outperformed the BM model.

Out of the discrete characters, only the hook configuration (HC) and ventral bar shape (VBS) produced strong phylogenetic signals (Fig. 4c). In contrast, we detected a relatively weak phylogenetic signal for the anchor similarity (AS) and no signal for the index of specificity (IS). Character maps (Fig. 9a) illustrate these phylogenetic pattern. Based on the literature survey, we suggested a series of measurements that are equivalents of HC, VBS, and AS. The HC summarises the lengths of the marginal hooks (U1–U7), VBS is described the width of the ventral bar in relation to its size (‘massive bar’ in Mendlová et al., 2012) but also includes information on the presence of membranous extensions, and AS likely refers to a difference in lengths of the inner roots of the dorsal vs. the ventral anchor pair and possibly a difference in size between the anchors pairs based on a drawing in Mendlová et al., (2012) but the publication only mentions a ‘difference in shape’. Character maps of these continuous measurements are plotted next to the equivalent discrete characters (Fig. 9b).

**Figure 8.**
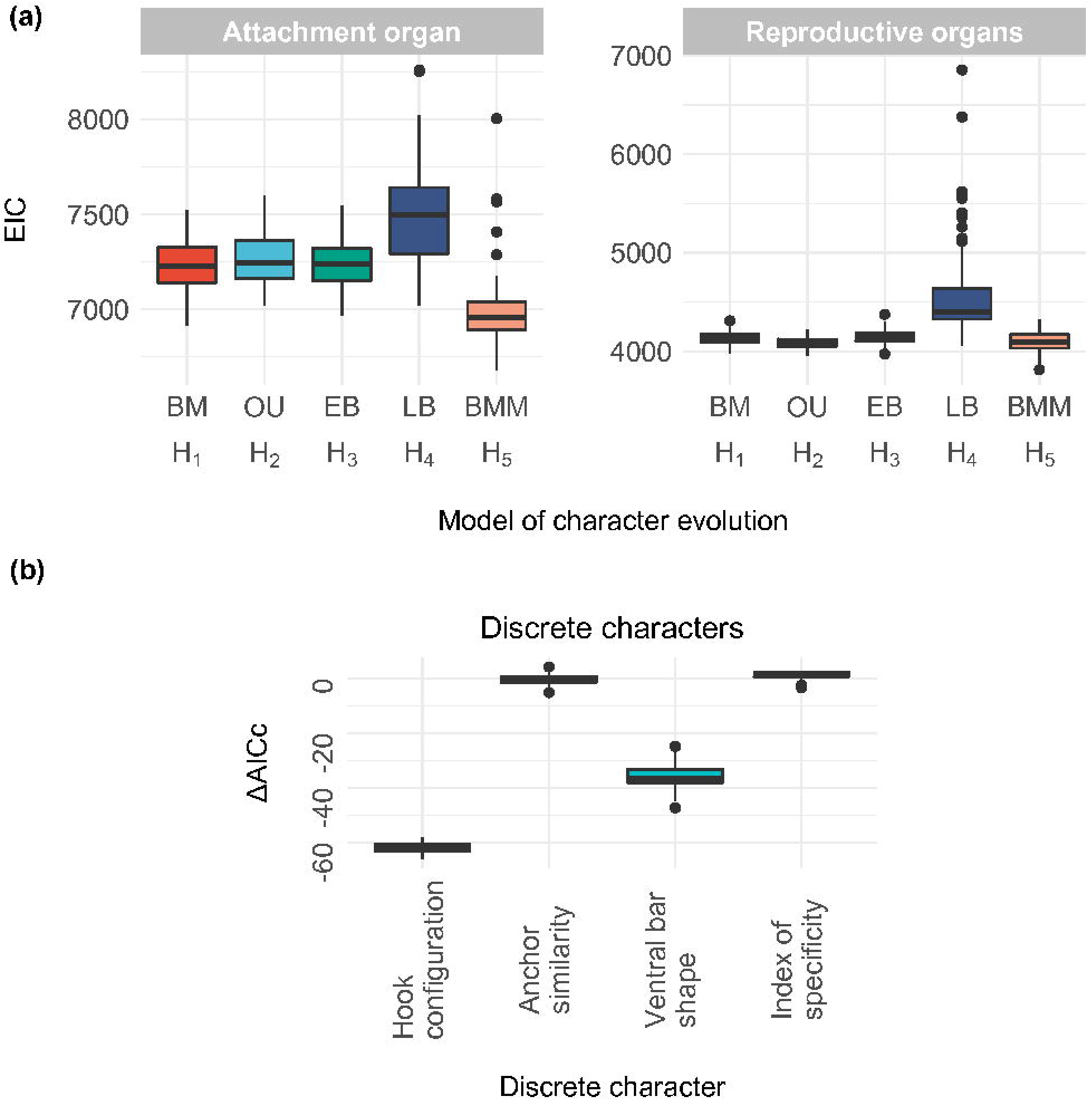
Results of phylogenetic comparative analysis applied to one set of 100 randomly sampled tree topologies from the Bayesian post–burn in phase. Fitted continuous models include Brownian Motion (BM), Ornstein-Uhlenbeck (OU), Early-Burst (EB), Late-Burst (LB), and multi-rate Brownian Motion (BMM) models simulating a random walk (H_1_), stabilising selection (H_2_), adaptive radiation with a decelerating (H_3_) and accelerating (H_4_) divergence of characters, and divergent BM regimes in parts of the phylogeny (H_5_). (a) Model fits were assessed through the Extended Information Criterion (EIC) for multivariate analysis. All models assuming a single evolutionary regime across the phylogeny performed similarly (LB performed worse). However, the BMM model suggesting a different regime for the EAR clade outperformed the latter. (b) Univariate PCMs for discrete characters including the hook configuration (HC) (Vignon et al. 2011), anchor similarity (AS), ventral bar shape (VBS) (Mendlová et al. 2012), and index of (host) specificity (IS) (Mendlová and Šimková 2014). HC and VBS show a strong phylogenetic signal, AS and IS shows no detectable phylogenetic signal. Model performance is assessed through difference in the sample size–corrected Akaike Information Criterion (ΔAICc) compared to a white noise model under the assumption of a star-like phylogeny (absolute phylogenetic independence).

**Figure 9.**
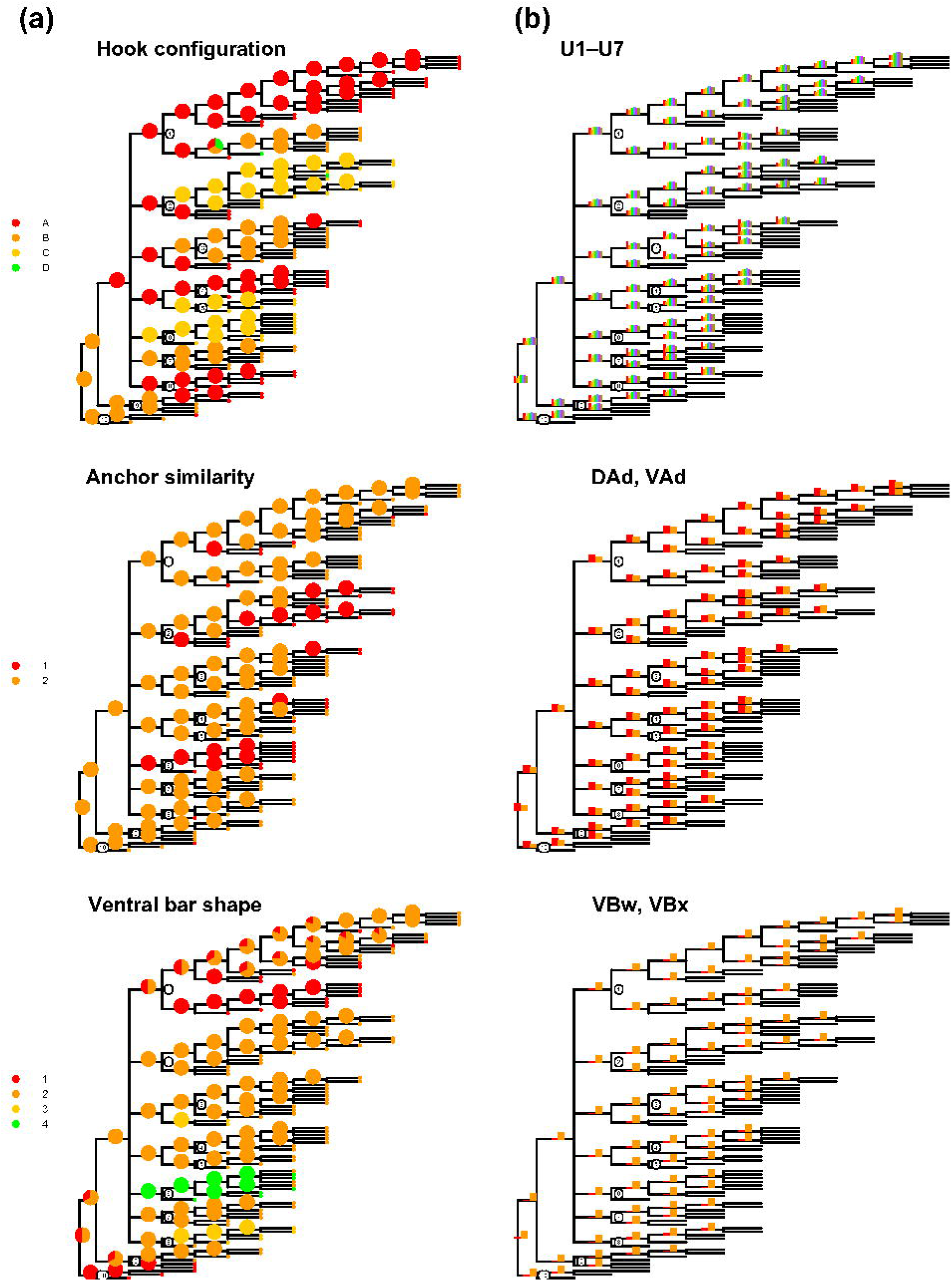
Character maps of discrete characters for the attachment organ morphology as defined by Vignon et al. (2011) and Mendlová et al. (2012) vs. the respective set of continuous measurement suggested in this study. The character maps indicate that summarising multiple variables into a single discrete character might cause the information loss observed as a lack of a phylogenetic signal in Fig. 8b. Numbers on nodes refer to clades in Fig. 2a. Hook configuration A, B, and C assigned according to criteria by Vignon et al. (2011) with relative size of hook pair 1 vs pairs 3–7; haptor configuration D equates to the configuration of *C. arthracanthus* with large hooks 1 and 3–7. Anchor similarity either similar (1) or dissimilar (2). Ventral bar shape with (1) or without (2) membranous extensions, massive with extensions (3) or support large plate (4).List of appendices (in print)

## Discussion

We investigated the evolution of the attachment and reproductive organs of parasites infecting a host lineage that has undergone multiple adaptive radiation events (Seehausen, 2006). Species of *Cichlidogyrus* (incl. the nested *Scutogyrus*) form a morphologically diverse lineage of parasitic flatworms infecting the gills of African cichlid fishes. This diversity alongside the model system status of the hosts led to the suggestion to use these parasites as a model system for parasite macroevolution (Pariselle et al., 2003; Vanhove et al., 2016). In this context, the present study is the most extensive reconstruction of intra-generic evolutionary relationships of African metazoan parasites to date providing new morphological, molecular, and host range data, which are made available in public databases. We were able to identify and characterise 11 different species groups (Fig. 2a; Fig. 6) amongst the 137 species that are currently known (Cruz-Laufer et al., 2021). For some of these groups, member species have previously been hypothesised to be closely related, e.g. species within the *Tylo, Hemi*, *CPO*, and ‘*Scutogyrus*’ groups. However, this study is the first to summarise all these groups and to analyse the evolution of the attachment and reproductive organ morphology using sequence data of 58 species and morphometric data of 136 species. We here hypothesise the phylogenetic position of described species of *Cichlidogyrus* and *Scutogyrus* without available molecular data through a synthesis of morphological, molecular, and host range data. However, due to missing data, e.g. a limited coverage of the four gene regions and a limited taxon coverage with the majority of species yet undiscovered (Cruz-Laufer et al., 2021), the relationship between the proposed species groups remains mostly unresolved with only moderate support for a few larger clades (Fig. 2a, b). In this light, we opted against taking any nomenclatural act such as splitting the genus into several new genera or synonymising *Scutogyrus* with *Cichlidogyrus* [for a more detailed discussion of this matter and similar cases within Dactylogyridae, see Kmentová et al. (2021a)].

### Systematic informativeness of morphometric measurements and host range data

To identify and classify species of *Cichlidogyrus,* previous publications have applied a specific set of measurements of the sclerotised attachment and reproductive organs (Cruz-Laufer et al., 2021). We investigated the systematic informativeness of the 29 measurements most frequently provided in the literature (Fig. 1c). Generally, the measurements appear to have a low systematic informativeness as a substantial portion of unsequenced species (70%) could not be placed in any of the clades inferred from the molecular phylogeny through a parsimony analysis combining molecular and morphometric data. Phylogenetic positions inferred from the combined MP consensus tree were in moderate accordance with the classifications based on character maps that considered additional discrete characters for the reproductive organs proposed here (Fig. 5b). The limited systematic informativeness of the measurements is also reflected by the performance of the classification analysis using support vector machines (SVMs). The optimised machine learning algorithm predicted species group affiliation within the sequenced species with moderate success (κ = 0.48). Yet for the unsequenced species, accordance of the SVM-based classification with the parsimony analysis (κ = 0.36) and literature survey (κ = 0.31) was comparatively low. One cause for this performance might be the low number of systematically informative measurements in the dataset as ten of these showed a moderate or low discriminatory power on average (AUC < 0.8, see Fig. 7a). Furthermore, the discriminative power of the measurements varied considerably between the different clades (Fig. 7b).

Notably, the phylogenetic signal was low for any of the evolutionary models applied to the reproductive organ in the context of the phylogenetic comparative method. This lack of information presents a stark contrast to the importance of the reproductive organ morphology to the character map–based classification. The phenotype of the male copulatory organ was crucial for characterising groups of related species in the form of several additional discrete characters proposed here (Appendix 2). These characters allowed us to consider information on the shape and diameter of the reproductive organs not included in the parsimony analysis and SVM-based approach. Therefore, we could provide hypotheses for all but 12 species regarding their phylogenetic position. This relative success highlights the value of phylogenetic information conveyed by reproductive organ structures; e.g. Rahmouni et al. (2017) noted that the heel of the male copulatory organ was shown to be similarly shaped in possibly related Eastern African species. Similar conclusions have been drawn for other dactylogyrid monogeneans such as species of *Characidotrema* Paperna & Thurston, 1968 (Řehulková et al., 2019). Regarding the lack of a phylogenetic signal of reproductive organ measurements, Pouyaud et al. (2006) proposed that this issue might be explained by a fast evolutionary rate but we suggest that this observation might be rather linked to a limited taxon coverage as more species have been described since. Instead, the low systematic informativeness indicated here more likely reflects a lack of systematically informative metrics as, for example, the number of reproductive organ measurements is low compared to the attachment organ (5 vs. 24). The measurements that are currently employed, fail to incorporate the systematic information offered by the attachment and reproductive organ morphology.

The low predictive power of the morphometrics has encouraged the use of loosely defined discrete characters in the past (Fig. 9). Several studies have categorised species according to their attachment organ morphology (Vignon et al., 2011; Mendlová et al., 2012) and host repertoires (Mendlová and Šimková, 2014). Here, we compared the systematic informativeness of these discrete characters against their continuous counterparts (Fig. 9). Two of the discrete characters lacked a phylogenetic signal (Fig. 8b) despite the moderate discriminatory power of the continuous characters suggested by the machine learning algorithm (Fig. 7a), a problem possibly caused by merging the information of multiple continuous variables into a single discrete character (Fig. 9). Discretisation can also introduce a researcher bias and produce misleading results. For instance, Mendlová et al. (2012) proposed that similar anchor shapes were an ancestral state of the binary character ‘shape of anchors’ (‘anchor similarity’ in the present study) based on a character map but the phylogenetic informativeness of the ‘shape of anchors’ character was never investigated further. In fact, the cluster analysis of the present study shows that dorsal anchor measurements and the respective ventral anchor measurements are highly multicollinear (Fig. 7c) with the exception of the inner roots (DAd and VAd). The ventral bar shape was also introduced as a variable for character mapping but no ancestral state could be inferred (Mendlová et al., 2012). Yet species of *Scutogyrus* have a unique phenotype with an extended plate associated to the bar structure (Pariselle and Euzet, 1995). These inconsistent results between continuous characters and their discretised counterparts underline that interpretations based on these discretised characters should be treated with caution. For instance, the hook configuration character states introduced by Vignon et al. (2011) are founded on gaps detected between the hook measurements of different species (see Pariselle and Euzet, 2009). New species discoveries in the last decade demonstrate that these character states might not be as clear-cut (see Rahmouni et al., 2017; Geraerts et al., 2020). Nonetheless, hook measurements were highly relevant in a machine learning context (Fig. 7a) indicating that, while an update to the coding of this character might be required, some character states might still prove useful for characterising species groups of *Cichlidogyrus*, e.g. the monophyletic *CPO* group species with well-developed hook pairs 3–7. If researchers wish to use discretised morphological characters in the future, character states should be redefined through quantitative character coding techniques such as proposed by Garcia-Cruz and Sosa (2006). The resulting character states would be more independent from researcher bias, which could increase the repeatability of such studies.

A discrete variable proposed to represent host specificity has also led to doubtful conclusions. Mendlová and Šimková, (2014) used the index of specificity (IS) to summarise the phylogenetic specificity of species of *Cichlidogyrus* (and *Scutogyrus*) into four categories including species-, genus-, and tribe-specific species, and generalists (Fig. 3a). Their study concluded that host specificity is related to parental care of the cichlid hosts. However, this observation was made due to an inherent sampling bias towards hosts raising their offspring through mouthbrooding (Cruz-Laufer et al., 2021), a specific type of parental care. Mendlová and Šimková (2014) also observed a correlation of the IS with the host phylogeny. However, this observation might again reflect a study bias in cichlid parasitology, in this case towards large economically relevant host species (Cruz-Laufer et al., 2021). Because of their importance to fisheries, these fishes unlike others have been introduced to various new habitats. Anthropogenic introductions might also increase the chances of co-introduction for their parasites and create ecological opportunities to expand host repertoires (e.g. Jorissen et al., 2020). To avoid these biases, future studies should treat host specificity as a complex parameter encompassing structural, phylogenetic, and geographical aspects (Poulin et al., 2011) as well as temporal fluctuations (Brooks et al., 2019). We suggest that these parameters could be included, e.g., in a genus-wide comparative study in the future.

In general, the size of morphological structures should first be treated as continuous, unmodified characters before further processing to avoid losing systematic information. Discretisation should only be applied with sufficient statistical evidence to back up character coding. We suggest increasing sampling and sequencing efforts to address data deficiency.

### Attachment organ evolution: rate heterogeneity, co-divergence, and host switching

To infer evolutionary patterns from the attachment organ morphology, we applied a penalised likelihood framework for multivariate phylogenetic comparative methods (PCMs) to the continuous character data. The evolutionary models representing different evolutionary hypotheses performed almost equally well (Fig. 4b). However, we found no significantly increased performance of the Ornstein-Uhlenbeck (OU), Early Burst (EB), and Late Burst (LB) models compared to the Brownian Motion (BM) model. Only the multi-rate BM (BMM) model performed slightly but significantly better than the other models (Fig. 4b). The OU, EB, and LB models present extensions of the Brownian Motion (BM) model (Butler and King 2004; Harmon et al. 2010). Therefore, the character evolution of the attachment organs measurements might simply follow a pattern associated with genetic drift or randomly fluctuating natural selection (Losos, 2008). This pattern might also be reflected in morphological features that are frequently shared by related species (Fig. 6). The relatedness of the parasite species could determine the evolution of attachment organ as suggested by a previous study (Vignon et al. 2011). Studies on a related lineage of monogeneans (*Ligophorus* Euzet & Suriano, 1977) also indicate a correlation of morphometrics and phylogenetic relationships (Rodríguez-González et al. 2017). However, the increased support for the BMM model suggests that the evolution of these parasites might have been shaped by different evolutionary regimes. This rate heterogeneity appears logical as host lineages outside the Great African lakes have not experienced the same explosive speciation as their East African relatives (see Brawand et al., 2015). In fact, purely single-rate models appear increasingly unlikely to capture the real evolutionary history of *Cichlidogyrus* because of its pan-African distribution (Cruz-Laufer et al., 2021) and ecological diversity of hosts (Burress, 2014). In the present study, we only modelled a multi-rate BM model because a lack of software packages suited for high-dimensional dataset with pre-set hypothesis. Yet other multi-rate models including OU, EB, and LB processes could provide better fits. In particular, species of the *EAR* group (Fig. 2a) that infect cichlid radiations in East Africa (Fig. 3) might mirror the explosive speciation of their hosts (see Vanhove et al., 2015). In recent years, multiple tools have been developed to investigate these more complex models but often these methods represent maximum likelihood (ML) approaches (e.g. Beaulieu et al., 2012; Ingram and Mahler, 2013; Puttick, 2018) that are sensitive to trait covariation, data orientation, and trait dimensions, particularly for high-dimensional datasets (Adams and Collyer, 2018). Other methods were developed to detect evolutionary rate shifts (e.g. Ingram and Mahler, 2013; Uyeda and Harmon, 2014; Khabbazian et al., 2016) rather than to test hypotheses based on previous knowledge (such as for the *EAR* clade in the present case). Therefore, penalised log-likelihood approaches that address these issues remain currently limited to multi-rate BM models but future updates will most likely close this technological gap.

Highly similar morphological characteristics can be found in multiple species groups (Fig. 5a; Fig. 6) such as a well-developed outer root of the dorsal anchor (e.g. species of the *Bulb* and *Cop* groups), large first hooks (e.g. species of *Tylo*, *EAR – Heel*, and *Hemi* groups), and long third to seventh hooks (e.g. species of the *EAR – Heel*, *CPO* groups). Character shifts observed within species groups appear to be adaptations to divergent host repertoires or environments (see Messu Mandeng et al., 2015). Host- and environmentally induced shifts in the attachment organ morphology have also been observed amongst other dactylogyrids, e.g. species of *Kapentagyrus* (Kmentová et al., 2018) and *Thaparocleidus* (Šimková et al., 2013), and other parasite lineages, e.g. cymothoid isopods (Baillie et al., 2019). For *Cichlidogyrus*, host species infected by parasites from the same species group frequently belong to the same tribe or related tribes of cichlids (Fig. 3; Appendix 5). Consequently, the phylogenetic congruence amongst some clades of *Cichlidogyrus* and their hosts is strong (Vanhove et al., 2015). Examples for host-induced character shifts amongst species of *Cichlidogyrus* can be found across the genus:

- *Cichlidogyrus amieti* has undergone a recent character shift as it displays small first hooks unlike other species of the *Hemi* clade, where they are well-developed. This change might result from a host switch (Messu Mandeng et al., 2015).
- *Cichlidogyrus sclerosus,* a generalist infecting fishes belonging to various cichlid tribes (Fig. 3), shares almost no attachment organ features with its sister species *C. amphoratus*, a specialist infecting only coptodonine cichlids (Fig. 3), beyond those shared by all species of the genus, e.g. auricles associated with the dorsal bar of the attachment organ (see Pariselle and Euzet, 2009).
- The diverse morphological features of species of the *EAR* group might relate to the ecological and morphological diversity of the hosts, e.g. related to feeding behaviour (Burress, 2014). The parasites, which we grouped into subgroups of the *EAR* clade, might have adapted to these ecologically diverse hosts resulting in divergent attachment and reproductive organ morphologies.
- Co-infections of different species of *Cichlidogyrus* can result in niche segregation on the gills of a host such as in species infecting Lake Victoria cichlids (Gobbin et al., 2020). At least some co-infecting species can be told apart based on their attachment organ morphology (see Gobbin et al., 2021). Thus, phenotypic differences might be linked to specific infection sites. Morphological adaptations to gill microhabitats have been reported for other monogeneans (Rohde, 1976; Ramasamy et al., 1985) but no study has so far investigated these mechanisms for species of *Cichlidogyrus*.

The host repertoires of the parasites offer clues for evolutionary processes frequently associated with host-induced effects in the evolutionary history of *Cichlidogyrus* including co-divergence and host switching. For instance, co-divergence might be reflected in the host (Fig. 3; Appendix 5) and geographical repertoires of species of *Cichlidogyrus* (Vanhove et al., 2013) (e.g. *EAR* group species are exclusive to Eastern Africa and *Bulb* group species to Southern Africa). Furthermore, a small yet increasing number of natural and invasion-induced host switches are reported in the literature from cichlids to non-cichlids, e.g. in the *Hemi* clade (Birgi and Euzet, 1983; Birgi and Lambert, 1986), and from African cichlids to cichlids and non-cichlids in the Americas, the Levant, and Madagascar (*C. arthracanthus, C. sclerosus, C. tiberianus, C. tilapiae*, *C. thurstonae,* and *Scutogyrus longicornis*) (Paperna, 1960; Jiménez-Garcia et al., 2001; Šimková et al., 2019). While the literature suggests that most African fish families have their own monogenean lineages (Carvalho Schaeffner, 2018) and the number of similar host switches might, hence, be limited, more host switches might be discovered in the future if more (also non-cichlid) host species are studied.

Despite these host-induced character shifts, the evolutionary models employed for multivariate PCMs detected no changes in the evolutionary rate of the attachment organ measurements. Apart from the limitation regarding multi-rate models, mentioned above, we suggest that this observation might be a result of incomplete sampling, which has affected similar studies in the past. For instance, previous studies were limited to single structures, i.e. the anchors (Rodríguez-González et al., 2017), or to the much lower number of species of *Cichlidogyrus* described at the time (77 vs. 130 today), which comprised less phenotypical diversity than is currently known to exist (Vignon et al., 2011). In fact, the phylogenetic position of some species with features diverging from their respective species groups (e.g. *C. amieti* from other *Hemi* group species) were not known. Observed similarities of the attachment organ morphology in related species reflected the specificity of the parasites to the respective host lineage rather than only the relatedness of the monogeneans. While the present analysis includes more species, DNA sequence data of less than half of all described species are available at GenBank and many species are likely to be discovered in the future (Cruz-Laufer et al., 2021). This data deficiency could affect the performance of the models implemented through PCMs as, e.g., the lack of lineages might lower the resolution of phylogenetic inference and increase phylogenetic uncertainty. A more complete taxon coverage of morphological, molecular, and ecological data will provide more insight into the evolutionary patterns of the genus. Furthermore, a closer look into the evolution of single species groups might reveal different patterns than for a genus-wide study. For instance, the high variability in the attachment and reproductive organ morphology of the *EAR* group might indicate a late burst in the divergence of characters induced by phenotypic adaptation to the rapidly speciating host lineages. A slightly better performance of the multi-rate model provides reason to further investigate the evolutionary history of this group, as it might fall under a divergent evolutionary regime. Lastly, convergent evolution of morphological characters might also have affected PCM model performance. Convergence can mask evolutionary processes that have occurred in the past as phenotypes become more similar (Losos, 2008). Monogenean parasites infecting the haplochromine radiations in the Eastern African lakes might be particularly impacted by this effect because of the convergent evolution of their hosts. These fishes have evolved via replicated adaptive radiation events resulting in similar yet unrelated species occupying the same niches in different lakes (McGee et al., 2016). This setting makes their parasites also potential targets to study convergent evolution in parasites. Conversely, the Eastern African diversity of *Cichlidogyrus* has mostly been investigated in Lake Tanganyika, the oldest lake, limiting the current possibilities to study parasite evolution in the context of replicate host radiations in this region [but see Gobbin et al. (2021) for a study focusing on Lake Victoria].

### Reproductive organ evolution

Unlike the attachment organ, the reproductive organs show no character changes correlated to host relatedness. Reproductive organ characters are evolutionarily stable within the different clades as evidenced by the role of the male copulatory organ in the morphological characterisation of the species groups. The evolution of the reproductive organs, unlike the attachment organ, has no apparent connection to the host range and seems to be determined mostly by phylogenetic constraints. Differences within species groups seem to reflect phylogenetic positions, e.g. the short penis of *C. falcifer* (*Hemi*) and simple accessory piece of *C. falcifer* and *C. cubitus* (*CPO*) are ancestral states (Fig. 5) of the respective clade. We detected a marginally better support for the BMM model (Fig. 8a) suggesting that the *EAR* group, with its host lineages that have resulted from adaptive radiation events, might have experienced an evolutionary regime slightly divergent from its congeners. However, the low phylogenetic signal in these data provides far less confidence in this result. Previous studies on the reproductive organs of monogenean flatworms have suggested that the complex surface of the reproductive organ might correlate with environmental factors, e.g. to avoid separation during copulation through the water flow (Kearn and Whittington, 2015). This hypothesis would suggest that the sclerotised structures evolve under selective pressure. However, while the screw thread–like thickening of the penis wall for some species (*C. casuarinus, C. centesimus*, *C. nshomboi*) (Fig. 6, *Heel* subgroup) might present such an example, we see no general trend to confirm this hypothesis. Neither generalist species such as *C. halli, C. tilapiae*, and *C. sclerosus* nor species infecting potentially more motile hosts living in fast-flowing waters, e.g. *C. consobrini, C. maeander,* and *C. ranula* infecting species of *Orthochromis*, display more complex MCOs than their congeners. Consequently, the diversification of the reproductive organs might be explained by reproductive isolation such as previously suggested for other dactylogyrid monogeneans (Šimková et al., 2002). A quantitative approach to measure reproductive organ shape could provide a more detailed answer to this question by capturing the shape variation currently only recorded as qualitative information (Fig. 5b) in taxonomic literature (see below for proposals).

### Classification, prediction, and research prospects

We were able to demonstrate the existing yet limited predictive power of the measurements currently employed by most taxonomists through a phylogenetic analysis under maximum parsimony accompanied by a machine learning approach. To improve the accuracy of these predictions overall, datasets with a more complete taxon coverage and higher number of observations are needed. The results of the character map–based classification vs. the parsimony analysis and the statistical classification approaches have highlighted a discrepancy between the systematic informativeness of the attachment and reproductive organ morphology in taxonomic literature and the low number of phylogenetically informative measurements. Future studies should address this discrepancy by increasing the number of systematically informative metrics.

Several possible solutions have been proposed to increase the number of metrics. First, some studies included novel measures such as the lengths of the shaft of the hooks, the curved length of the accessory piece, or the width of the penis tube and the bulbous portion of the penis (e.g. Geraerts et al., 2020) to describe species of *Cichlidogyrus* in more detail. Second, geomorphometric analyses have been employed for the anchors of some species of *Cichlidogyrus* (Rahmouni et al., 2021) as well as other monogeneans (Vignon and Sasal, 2010; Rodríguez-González et al., 2017; Kmentová et al., 2020) to infer the most systematically informative structures. Third, 3D imaging techniques as previously developed for monogenean sclerotised structures (e.g. García-Vásquez et al., 2012) have been proposed to improve characterisations of species of *Cichlidogyrus* (Cruz-Laufer et al., 2021) using methods employed for shape analyses (Klingenberg, 2010; Dryden and Mardia, 2016). Key characters that could be inferred using this approach are for example elongation (long vs round), torsion (straight vs coiled), and complexity (few vs many extensions). However, all of these solutions remain currently unavailable for comparative analyses. Data resulting from the proposed additional measures, geomorphometric measurements, and 3D imaging techniques are available for only few morphological structures, species, and studies. Future research should increase the taxon coverage for proposed methods and improve quantification methods to address these limitations. Future datasets should encompass an increasing number of DNA sequences and morphometric measurements from all known species as this study highlights the limitations of using morphological characters to infer the evolution of monogenean parasites. Nonetheless, raw data should regularly be published in supplements or data repositories (e.g. Kmentová et al., 2016b) to improve the accessibility of morphometric data. This approach could enhance information flow and academic capacity for research into species of monogenean parasites in the future (Cruz-Laufer et al., 2021).

## Conclusion

Due to the species-richness and the model system status of their cichlid hosts, the parasite lineage including species of *Cichlidogyrus* and *Scutogyrus* could be an ideal macroevolutionary model system for speciation in parasitic organisms in the future. Notably, these species belong to one of the more extensively investigated lineages in close interaction with an adaptive radiation of other species, namely cichlid fishes. As adaptive radiations might explain a substantial part of the biodiversity on the planet, many species potentially interact with radiating lineages. Therefore, understanding evolutionary history in this context could be key for understanding the diversity of species interactions in general.

Similar to many parasite systems, the limited availability of phenotypic and genotypic data for the 137 species of *Cichlidogyrus* currently described has hampered investigations into the evolutionary history of the lineage. This study provides an example of how an increased availability of morphological and DNA sequence data can promote computational approaches such as parsimony analyses, phylogenetic comparative methods, and machine learning classification to answer basic questions of evolutionary research. The phylogenetically informative measurements (continuous and discrete, established and newly proposed) and characterisation of multiple clades within *Cichlidogyrus* have highlighted key information required to describe new species as well as the shortcomings of the morphological characters that are currently used. Features of the attachment organ used to characterise the proposed species groups remain key descriptors alongside the features of the reproductive organs. However, they only serve as predictors in a multivariate context. This conclusion presents a stark contrast to previous classification systems for this taxon, which used to predict species groups based on single discrete morphological characters.

Using multivariate phylogenetic comparative methods, we were able to model evolutionary mechanisms of the attachment and reproductive organs suggested by previous studies. No support for alternative models suggests that the variation of the morphology of these organs across the genus has been shaped by different evolutionary regimes. Morphological patterns in different species groups further indicate that the attachment organ morphology can be shaped by host parameters. In contrast, the reproductive organs appear to follow a more random evolutionary regime. Other mechanisms might play a role in the evolution of this parasite lineage, e.g. within-host speciation as suggested by other studies. Therefore, more morphometric and DNA sequence data will be needed to provide a more detailed picture of the evolution of the genus. We encourage researchers to publish morphometric raw data regularly to improve the data accessibility for the scientific community, and to sequence systematically informative DNA regions of the remaining described or undiscovered species.

## Acknowledgements

We thank J.-F. Agnèse (Institut de Recherche pour le Développement & Université de Montpellier) for taking responsibility for part of the molecular work, and F. A. M. Volckaert (KU Leuven) and J. Snoeks (Royal Museum for Central Africa & KU Leuven) for their guidance since the early stages of this research. J. Bamps, A. F. Grégoir, L. Makasa, J. K. Zimba (Department of Fisheries), C. Katongo (University of Zambia), T. Veall, O. R. Mangwangwa (Rift Valley Tropicals), V. Nshombo Muderhwa, T. Mulimbwa N’sibula, D. Muzumani Risasi, J. Mbirize Ndalozibwa, V. Lumami Kapepula (Centre de Recherche en Hydrobiologie-Uvira), the Schreyen-Brichard family (Fishes of Burundi), F. Willems (Kasanka Trust), S. Dessein (Botanic Garden Meise), A. Chocha Manda, G. Kapepula Kasembele, E. Abwe, B. Katemo Manda, C. Mukweze Mulelenu, M. Kasongo Ilunga Kayaba and C. Kalombo Kabalika (Université de Lubumbashi), M. Collet and P. N’Lemvo (Institut Congolais pour la Conservation de la Nature), D. Kufulu-ne-Kongo (École Muilu Kiawanga), L. Matondo Mbela (Université Kongo), S. Wamuini Lunkayilakio, P. Nguizani Bimbundi, B. Boki Fukiakanda, P. Ntiama Nsiku (Institut Supérieur Pédagogique de Mbanza-Ngungu), P. Nzialu Mahinga (Institut National pour l’Etude et la Recherche Agronomiques— Mvuazi/Institut Supérieur d’études agronomiques—Mvuazi) and M. Katumbi Chapwe are thanked for administrative, field and lab support, making this study possible. We thank A. Avenant-Oldewage, M. Geraerts, T. Gobbin, P. C. Igeh, N. Kmentová, M. Mendlová, and C. Rahmouni for providing raw species measurements used for their respective publications. Furthermore, we thank J. M. Mirande and T. Moons for technical advice on the use of *TNT*. Field work was supported by travel grants V.4.096.10.N.01, K.2.032.08.N.01 and K220314N from the Research Foundation—Flanders (FWO-Vlaanderen) (to MPMV), two travel grants from the King Leopold III Fund for Nature Conservation and Exploration (to MPMV and MVS), FWO-Vlaanderen Research Programme G.0553.10, the University Development Cooperation of the Flemish Interuniversity Council (VLIR-UOS) South Initiative ZRDC2014MP084, the OCA type II project S1_RDC_TILAPIA and the Mbisa Congo project (2013–2018), the latter two being framework agreement projects of the RMCA with the Belgian Development Cooperation. When data collection for this study commenced, MVS and MPMV were PhD fellows, and TH a post-doctoral fellow, of FWO-Vlaanderen. Part of the research leading to results presented in this publication was carried out with infrastructure funded by the European Marine Biological Research Centre (EMBRC) Belgium, Research Foundation – Flanders (FWO) project GOH3817N. MWPJ was supported by the Belgian Federal Science Policy Office (BRAIN-be Pioneer Project (BR/132/PI/TILAPIA), and a BOF Reserve Fellowship from Hasselt University. AJCL is funded by Hasselt University (BOF19OWB02), and MPMV receives support from the Special Research Fund of Hasselt University (BOF20TT06). We thank the reviewers including J. M. Mirande and an anonymous reviewer for suggesting improvements to the manuscript.

## Conflict of interest

The authors declare that they have no conflict of interest.

## Data availability statement

The morphological data that support the findings of this study are openly available in MorphoBank at www.morphobank.org, at https://dx.doi.org/XXXXXXXX. The DNA sequence data are openly available in the GenBank Nucleotide Database at https://www.ncbi.nlm.nih.gov/genbank, accession numbers XXXXXX–XXXXXX. Phylogenetic trees and data matrices are openly available in TreeBase at https://treebase.org, accession number XXXXXX.

---

**Appendix 1.** Sampling localities and dates of specimens belonging to *Cichlidogyrus* and *Scutogyrus* collected for this study including a reference for the samples previously included in taxonomic studies. Freshwater ecoregions are assigned according to Thieme et al. (2005).

**Appendix 2.** List of continuous and discrete characters and character states inferred from the taxonomic literature and used for character mapping in Fig. 1. All characters represent characteristics that are commonly used to described species of *Cichlidogyrus* and *Scutogyrus*.

**Appendix 3.** Specimen data of cichlid parasites of the genera *Cichlidogyrus* and *Scutogyrus* used for phylogenetic analyses including host species, GenBank accession numbers, locality by country, and reference. Voucher/isolate ID and accession numbers in italics indicate specimens not included in subset trees used for phylogenetic comparative methods.

**Appendix 4.** Substitution models of molecular evolution and partitions for Bayesian inference (BI) and maximum likelihood estimation (ML) of phylogeny of species of *Cichlidogyrus* and *Scutogyrus*. Models include the general time reversible model (GTR), the Kimura 1980 model (K80), the transitional model 3 with unequal base frequencies (TIM3e), the Tamura-Nei model (TN), and the three-parameter model 2 (TPM2) plus empirical base frequencies (+ F), a proportion of invariable sites (+ I), a discrete Γ model with four rate categories (Γ4), or a FreeRate model with three categories (+ R3). For model specification see the IQ-TREE ModelFinder manual (Kalyaanamoorthy et al., 2017).

**Appendix 5.** Species groups of *Cichlidogyrus* with species included, species potentially included, and the respective host ranges reported in the taxonomic literature including host species of candidate species.

**Appendix 6.** Ancestral states of continuous characters inferred from a literature survey and used for character mapping. Character IDs refer to numbers in Appendix 2.

**Appendix 7.** Ancestral states of discrete characters inferred from a literature survey and used for character mapping. Values represent probabilities for different character states. Character IDs refer to numbers in Appendix 2.

## Supporting Information

**File S1.** Input data matrix for use in *TNT* including morphometric and DNA sequence data.

**File S2.** Raw morphometric measurements for species of *Cichlidogyrus* and *Scutogyrus*.

## Notes

### Competing Interest Statement

The authors have declared no competing interest.

### Summary of Updates

Whole manuscript revised, stronger focus on parasite evolution and parsimony analyses, character maps added

