## Appendices for "Somewhere I belong: phylogeny and morphological evolution in a species-rich lineage of ectoparasitic flatworms infecting cichlid fishes"

Appendices (in print)

**Appendix 1.** Sampling localities and dates of specimens belonging to *Cichlidogyrus* and *Scutogyrus* collected for this study including a reference for the samples previously included in taxonomic studies. Freshwater ecoregions are assigned according to Thieme et al. (2005).

| **Species** | **Host species** | **Host family** | **Country** | **Freshwater Ecoregion** | **Locality** | **Date** | **Lon.** | **Lat.** | **Reference** |
| --- | --- | --- | --- | --- | --- | --- | --- | --- | --- |
| *Cichlidogyrus amieti* Birgi & Euzet, 1983 | *Aphyosemion cameronense* (Boulenger, 1903) | Notho-branchiidae | Cameroon | Southern Gulf of Guinea Drainages | swampy area on the road from Edéa to Kribi | 2015 | 3.35 | 10.08 |  |
| *Cichlidogyrus berrebii* Pariselle & Euzet, 1994 | *Tylochromis jentinki* (Steindachner, 1894) | Cichlidae | Côte d'Ivoire | Eburneo | Fresco Lagoon, Bolo | 17/01/2010 | 5.10 | -5.58 | Bitja-Nyom et al. (2021) |
| *Cichlidogyrus buescheri* Pariselle & Vanhove, 2015 (n = 2) | *Interochromis loocki* (Poll, 1949) | Cichlidae | Zambia | Lake Tanganyika | Kalambo Lodge | Apr 2008 | -8.62 | 31.20 | Pariselle et al. (2015b) |
| *Cichlidogyrus casuarinus* Pariselle, Muterezi Bukinga & Vanhove, 2015  (n = 4) | *Bathybates minor* Boulenger, 1906 | Cichlidae | DR Congo | Lake Tanganyika | Mpala | 21/04/2010 | -6.75 | 29.53 | Pariselle et al. (2015a) |
| *Cichlidogyrus centesimus* Vanhove, Volkaert & Pariselle, 2011 | *Ophthalmotilapia ventralis* (Boulenger, 1898) | Cichlidae | DR Congo | Lake Tanganyika | Kikoti | Apr 2010 | -7.19 | 30.07 | Vanhove et al. (2011) |
| *Cichlidogyrus* cf. *bychowskii* | *Hemichromis bimaculatus* Gill, 1862 | Cichlidae | Côte d'Ivoire | Eburneo | Kossou, Bandama | 21/01/2010 | 7.03 | -5.48 | Bitja-Nyom et al. (2021) |
| *Cichlidogyrus cirratus* Paperna, 1964 | *Oreochromis aureus* (Steindachner, 1964) | Cichlidae | Morocco | Atlantic Northwest Africa | Guelta Zerga | Oct 2019 | 28.43 | -10.80 |  |
| *Cichlidogyrus cirratus* Paperna, 1964 | *Oreochromis niloticus* (Linnaeus, 1758) | Cichlidae | France | NA | Montpellier, Aquaculture facilities | 01/06/2011 | 43.63 | 3.87 |  |
| *Cichlidogyrus consobrini* Jorissen, Pariselle & Vanhove in Jorissen et al. 2018b | *Sargochromis mellandi* (Boulenger, 1905) | Cichlidae | DR Congo | Bangweulu-Mweru | Kipopo Inera station | Aug 2014 | -11.56 | 27.35 | Jorissen et al. (2018b) |
| *Cichlidogyrus cubitus* Dossou, 1982 | *Coptodon guineensis* (Günther, 1862) | Cichlidae | Morocco | Atlantic Northwest Africa |  | Dec 2018 | 27.93 | -11.42 |  |
| *Cichlidogyrus cubitus* Dossou, 1982 | *Pelmatolapia mariae* (Boulenger, 1899) | Cichlidae | Côte d'Ivoire | Eburneo | Fresco Lagoon, Bolo | 18/01/2010 | 5.10 | -5.58 | Bitja-Nyom et al. (2021) |
| *Cichlidogyrus digitatus* Dossou, 1982 | *Pelmatolapia mariae* (Boulenger, 1899) | Cichlidae | Cameroon | Southern Gulf of Guinea Drainages | Edéa, Lake Ossa | 30/12/2015 | 3.78 | 10.02 | Bitja-Nyom et al. (2021) |
| *Cichlidogyrus dossoui* Douëllou, 1993 (n = 2) | *Coptodon rendalli* (Boulenger, 1897) | Cichlidae | Zambia | Bangweulu-Mweru | Bangweulu Wetlands | July 2010 | -11.95 | 30.25 | Vanhove et al. (2013) |
| *Cichlidogyrus falcifer* Dossou & Birgi, 1984 | *Hemichromis fasciatus* Peters, 1857 | Cichlidae | Ivory Coast | Eburneo | Fresco Lagoon, Bolo | 18/01/2010 | 5.10 | -5.58 | Bitja-Nyom et al. (2021) |
| *Cichlidogyrus gillardinae* Muterezi Bukinga, Vanhove, Van Steenberge & Pariselle, 2012 (n = 2) | *Astatotilapia burtoni* (Günther, 1894) | Cichlidae | DR Congo | Lake Tanganyika | Kalemie fish market | Apr 2010 | -5.90 | 29.18 | Muterezi Bukinga et al. (2012) |
| *Cichlidogyrus halli* (Price & Kirk, 1967) | *Oreochromis niloticus* (Linnaeus, 1758) | Cichlidae | Mali | Upper Niger | Bamako, River Niger | 01/02/2010 | 12.63 | -7.98 | Bitja-Nyom et al. (2021) |
| *Cichlidogyrus irenae* Gillardin, Vanhove, Pariselle, Huyse & Volckaert, 2012 (n = 2) | *Gnathochromis pfefferi* (Boulenger, 1898) | Cichlidae | Zambia | Lake Tanganyika | Kalambo Lodge | Apr 2008 | -8.62 | 31.20 | Gillardin et al. (2012) |
| *Cichlidogyrus kothiasi* Pariselle & Euzet, 1994 | *Tylochromis jentinki* (Steindachner, 1894) | Cichlidae | Côte d'Ivoire | Eburneo | Fresco Lagoon, Bolo | 17/01/2010 | 5.10 | -5.58 | Bitja-Nyom et al. (2021) |
| *Cichlidogyrus longicirrus* Paperna, 1965 | *Hemichromis elongatus* (Guichenot, 1861) | Cichlidae | Cameroon | Northern Gulf of Guinea Drainages | Idénao, unnamed river | 24/02/2008 | 4.20 | 8.98 | Bitja-Nyom et al. (2021) |
| *Cichlidogyrus mbirizei* Muterezi Bukinga, Vanhove, Van Steenberge & Pariselle, 2012 | *Oreochromis tanganicae* (Günther, 1894) | Cichlidae | DR Congo | Lake Tanganyika |  | Apr 2010 |  |  | Muterezi Bukinga et al. (2012) |
| *Cichlidogyrus nandidae* Birgi & Lambert, 1986 | *Polycentropsis abbreviata* (Boulenger, 1901) | Poly-centridae | Cameroon | Southern Gulf of Guinea Drainages | swampy area on the road from Edéa to Kribi | 2015 | 3.35 | 10.08 |  |
| *Cichlidogyrus nshomboi* Muterezi Bukinga, Vanhove, Van Steenberge & Pariselle, 2012 (n = 2) | *Boulengerochromis microlepis* (Boulenger, 1899) | Cichlidae | DR Congo | Lake Tanganyika | Mulembwe | 09/04/2010 | -6.12 | 29.27 | Muterezi Bukinga et al. (2012) |
| *Cichlidogyrus papernastrema* Price, Peebles & Bamford, 1969 | *Tilapia sparrmanii* Smith, 1840 | Cichlidae | DR Congo | Bangweulu-Mweru | Jardin Zoologique de Lubumbashi | Apr 2016 | -11.65 | 27.47 | Jorissen et al. (2018b) |
| *Cichlidogyrus quaestio* Price, Peebles & Bamford, 1969 | *Coptodon rendalli* (Boulenger, 1897) | Cichlidae | DR Congo | Bangweulu-Mweru | Luapula River off Kashobwe | 08/09/2014 | -9.67 | 8.62 | Jorissen et al. (2018b) |
| *Cichlidogyrus sanseoi* Pariselle & Euzet 2004 | *Hemichromis fasciatus* Peters, 1857 | Cichlidae | Senegal | Senegal-Gambia | Djoudj jetty, Senegal River | 05/03/2008 | 16.40 | -16.30 | Bitja-Nyom et al. (2021) |
| *Cichlidogyrus schreyenbrichardorum* Pariselle & Vanhove, 2015 (n = 2) | *Interochromis loocki* (Poll, 1949) | Cichlidae | Zambia | Lake Tanganyika | Kalambo Lodge | Apr 2008 | -8.62 | 31.20 | Pariselle et al. (2015b) |
| *Cichlidogyrus sclerosus* Paperna & Thurston, 1969 | *Oreochromis mossambicus* (Peters, 1852) | Cichlidae | DR Congo | Lake Tanganyika | Uvira, Ruzizi | 2013 | -3.33 | 29.17 |  |
| *Cichlidogyrus sclerosus* Paperna & Thurston, 1969 | *Oreochromis niloticus* (Linnaeus, 1758) | Cichlidae | France | NA | Montpellier, Aquaculture facilities | 01/06/2011 | 43.63 | 3.87 |  |
| Cichlidogyrus sclerosus Paperna & Thurston, 1969 | *Oreochromis mossambicus* (Peters, 1852) | Cichlidae | Mozambique | Limpopo | Linlangalinwe | 2010 |  |  | Firmat et al. (2016) |
| *Cichlidogyrus teugelsi* Pariselle & Euzet 2004 | *Hemichromis fasciatus* Peters, 1857 | Cichlidae | Ivory Coast | Eburneo | Fresco Lagoon, Bolo | 18/01/2010 | 5.10 | -5.58 | Bitja-Nyom et al. (2021) |
| *Cichlidogyrus thurstonae* Ergens, 1981 | *Oreochromis niloticus* (Linnaeus, 1758) | Cichlidae | DR Congo | Lower Congo | Monzi | Apr 2016 | -5.62 | 13.23 |  |
| *Cichlidogyrus tiberianus* Paperna, 1960 | *Coptodon rendalli* (Boulenger, 1897) | Cichlidae | Zambia | Bangweulu-Mweru | Bangweulu Wetlands | July 2010 | -11.95 | 30.25 | Vanhove et al. (2013) |
| *Cichlidogyrus vandekerkhovei* Vanhove, Volkaert & Pariselle, 2011 (n = 2) | *Ophthalmotilapia ventralis* (Boulenger, 1898) | Cichlidae | DR Congo | Lake Tanganyika | Kikoti | Apr 2010 | -7.19 | 30.07 | Vanhove et al. (2011) |
| *Cichlidogyrus vealli* Pariselle & Vanhove, 2015 (n = 2) | *Interochromis loocki* (Poll, 1949) | Cichlidae | Zambia | Lake Tanganyika | Kalambo Lodge | Apr 2008 | -8.62 | 31.20 | Pariselle et al. (2015b) |
| *Cichlidogyrus yanni* Pariselle & Euzet, 1996 | *Sarotherodon melanotheron* Rüppel, 1852 | Cichlidae | France | NA | Montpellier, Aquaculture facilities | 01/06/2011 | 43.63 | 3.87 |  |
| *Cichlidogyrus zambezensis* Douëllou, 1993 | *Serranochromis macrocephalus* (Boulenger, 1899) | Cichlidae | DR Congo | Bangweulu-Mweru | Lake Kipopo | 08/09/2014 | -11.57 | 27.35 | Jorissen et al. (2018b) |
| *Cichlidogyrus zambezensis* Douëllou, 1993 | *Serranochromis macrocephalus* (Boulenger, 1899) | Cichlidae | Zambia | Bangweulu-Mweru | Bangweulu Wetlands | July 2010 | -11.95 | 30.25 | Vanhove et al. (2013) |
| *Scutogyrus bailloni* Pariselle & Euzet, 1995 | *Sarotherodon galilaeus* (Linnaeus, 1758) | Cichlidae | Côte d'Ivoire | Eburneo | Comoé, Bridge close to Anekouadiokro | 27/01/2010 | 7.12 | -6.32 | Bitja-Nyom et al. (2021) |
| *Scutogyrus gravivaginus* (Paperna & Thurston 1969) | *Oreochromis mweruensis* Trewavas, 1983 | Cichlidae | DR Congo | Bangweulu-Mweru | Kishwishi River near Futuka Farm | Apr 2016 | -11.48 | 27.65 | Jorissen et al. (2018b) |
| *Scutogyrus longicornis* (Paperna & Thurston 1969) | *Oreochromis niloticus* (Linnaeus, 1758) | Cichlidae | Mali | Upper Niger | Bamako, River Niger | 01/02/2010 | 12.63 | -7.98 | Bitja-Nyom et al. (2021) |
| *Scutogyrus longicornis* (Paperna & Thurston 1969) | *Oreochromis niloticus* (Linnaeus, 1758) | Cichlidae | Madagascar | North-western Madagascar | Lake Andranotapahina | Apr 2016 | -18.39 | 47.42 | Jorissen et al. (2018b) |
| *Scutogyrus minus* (Dossou, 1982) | *Sarotherodon melanotheron* Rüppel, 1852 | Cichlidae | Côte d'Ivoire | Eburneo | Layo, Ebrié Lagoon | 23/01/2010 | 5.31 | -4.30 | Bitja-Nyom et al. (2021) |
| *Scutogyrus vanhovei* Pariselle, Bitja Nyom & Bilong Bilong, 2013 | *Pelmatolapia mariae* (Boulenger, 1899) | Cichlidae | Cameroon | Southern Gulf of Guinea Drainages | Edéa, Lake Ossa | 30/12/2015 | 3.78 | 10.02 | Bitja-Nyom et al. (2021) |

**Appendix 2.** List of continuous and discrete characters and character states inferred from the taxonomic literature and used for character mapping in Fig. 1. All characters represent characteristics that are commonly used to described species of *Cichlidogyrus* and *Scutogyrus*.

Anchors

The size of the anchors relative of each other is often reported in species descriptions.

1.	*Size of the anchors without taking the anchor roots into account: DAb/VAb.*

Mendlová et al. (2012) classified species according to the ‘shapes’ of the dorsal and ventral anchors. Based on the drawing in Figure 6 (1.1 and 1.2) in this publication, we hypothesis were mostly referring to the elongated inner root (VAd or DAd) in some species. Thus, we think that the ‘shape of anchors’ is best represented by the measurements of the entire anchor from the tip of the inner root to the beginning of the point.

2.	*Size of the anchors taking the anchor roots into account: DAa/VAa.*

The literature often reports the lengths of the roots in the anchors relative to each other. Thus, we suggest the following two measurements:

3.	*Dorsal anchor root ratio: DAd/DAc*.

4.	*Ventral anchor root ratio: VAd/VAc*.

Dorsal bar

In species description, one of the most reported characteristics is the length of the auricles, which can be elongated such as in species of *Scutogyrus* (Pariselle and Euzet, 1995) or short such as in species of *Cichlidogyrus* infecting tylochromine cichlids (Pariselle and Euzet, 1994). Mendlová et al. (2012) suggested three discrete states based on the length of the auricles but here we reflect the auricle length simply as continuous measurement in relation to the bar size.

5.	*Relative length of auricles: DBh/DBx*

Ventral bar

Mendlová et al. (2012) provide four character states for the shape of the ventral bar. One aspect of this classification, the presence of membranous extensions, cannot be reflected through the morphometric measurements that are currently used. However, some species are reported to possess a ‘massive’ ventral bar, which we here reflect as the width of the bar relative to its size.

6.	*Relative width: VBw/VBx*

Hook configuration

The hook configuration is one of the most widely used characters to group species of *Cichlidogyrus* and *Scutogyrus*. In its current form, it was first proposed by Vignon et al. (2011) but continues to be used also in the most recent species description (Geraerts et al., 2020). The hook configuration is a representation of the length of the hooks in relation to each other. The second hook pair remains constant across all species as it maintains its embryonal size (Llewellyn, 1963). Thus, we use its size to standardise measurements of the remaining marginal hooks. Furthermore, pairs 3–7 are often considered as single block (e.g. Gillardin et al., 2012) with little difference observed between them (see also Fig. 7). While this observation might reflect a sampling bias towards species with similarly sized U3–7, we opted to represent these hooks through the measurement of U7 as the most common hook to be measured (Table S2) most likely because of its less peripheral position on mounted specimens.

7.	*Relative size of first hook pair: U1/U2*

8.	*Relative size of third to seventh hook pair: U7/U2*

Male copulatory organ (MCO)

The shapes and length of the MCO are rarely used to group species. However, they are provided in species descriptions. Here, we opt to keep the length of the penis, accessory piece, and heel as measurements but add two additional discrete characters to described their shapes. The length and width of the auxiliary plate are summarised as surface area assuming an ellipsoid shape.

9.	*Length of penis: Pe*

10.	*Shape of penis*

a.	Straight: penis more or less straight with no strong arching, twisting, looping, or spiralling but can be slightly sinuous or arched.

b.	Straight, thick-walled: same as before but wall of penis present thickening.

c.	Arched: penis strongly arched in one direction, distal portion often held in position by accessory piece.

d.	Looped: penis draws a loop in the shape of a G.

e.	Large loop: penis draws large circle ending in distal portion of accessory piece.

f.	Spiralled: penis draws spiral in large radius.

g.	Spirally coiled: penis draws spiral in small radius in the shape of a helix

11.	*Diameter of penis*

a.	Tubular: penis in the shape of a simple tube.

b.	Widened: penis widened.

c.	Bulbous: penis presents a bulbous portion (outside the basal bulb).

12.	*Length of accessory piece: AP*

13.	*Shape of accessory piece:*

a.	Simple: elongated accessory piece without additional structures mentioned in the other character states but species with more unique structures such as connecting stalks and caps are also included here.

b.	Furcated: accessory piece present one or more furcations.

c.	Distal hook: Accessory piece ends in a single distal hook.

d.	Distal flap: Accessory piece ends in a single distal flap.

e.	Gutter-like: Accessory piece in the shape of a gutter guiding the penis.

f.	Ribbon-like: Accessory piece is a flattened structure in the shape of a ribbon or drape.

g.	Spirally coiled: Accessory piece in the shape of simple helix.

h.	Looped: Accessory piece draws a loop in the shape of a G.

i.	Reduced: Accessory piece reduced to a thin, string-like structure or absent.

j.	Complex, S-shaped: massive, roughly S-shaped accessory piece (Fig. 6: *CPO*) that is frequently connected to the heel. The accessory piece has an extension or thickening at the first turn in the proximal half and frequently displays a folded back, straight and pointy, or hook-like distal end, or sometimes additional terminations resulting in a furcate ending with two or three digitations. However, the first turn is never V-shaped or knee-like such as in (l) and the hook-shaped termination is never sickle-like such as in (c).

k.	Complex, C-shaped: complicated roughly C-shaped accessory piece often with finger or hook-shaped outgrowths and marked heel (Fig. 6: *Oreo1*).

l.	Two portions, V-shaped: accessory piece consists of two distinct portions shaped like a V with an expanded knee-like bend (Fig. 6: *Hemi*)

m.	Two portions, spiralling: accessory piece consists of two distinct portions, large spiral followed by non-spiralled distal portion.

n.	In two parts: accessory piece consists of two distinct, superimposed parts.

14.	*Size of heel: He*

15.	*Surface area of auxiliary plate: AuP*

Sclerotised vagina

Similar to the MCO the sclerotised vagina is often measured and described but rarely used to group species. We keep both measurements here and add a discrete character to depict the shape of the structure.

16.	*Length of vagina: Vgl*

17.	*Width of vagina: Vgw*

18.	*Shape of vagina:*

a.	Non-sclerotised: Vagina not sclerotised.

b.	Tubular: Vagina in the shape of a simple tube.

c.	Bulbous: Vagina widened in at least one portion.

d.	Spiralled: tubular vagina that draws a spiral.

**Appendix 3.** Specimen data of cichlid parasites of the genera *Cichlidogyrus* and *Scutogyrus* used for phylogenetic analyses including host species, GenBank accession numbers, locality by country, and reference. Voucher/isolate ID and accession numbers in italics indicate specimens not included in subset trees used for phylogenetic comparative methods.

| **Parasite** | **Host** | **Isolate/ Voucher** | **GenBank accession numbers** | | | | **Locality** | **Reference** |
| --- | --- | --- | --- | --- | --- | --- | --- | --- |
|  |  |  | 28S rDNA | 18S rDNA | ITS1 rDNA | CO1 mtDNA |  |  |
| *Cichlidogyrus acerbus*  Dossou, 1982 | *Sarotherodon galilaeus*  (Linnaeus, 1758) | PC75 | HQ010036.2 | HE792780.2 | HE792780.2 |  | Senegal | Mendlová et al. (2012); Mendlová et al. (2010) |
| *Cichlidogyrus aegypticus*  Ergens, 1981 | *Coptodon guineensis*  (Günther, 1862) | PC16 | HQ010021 | HE792781 | HE792781 |  | Senegal | Mendlová et al. (2012); Mendlová et al. (2010) |
| *Cichlidogyrus agnesi*  Pariselle & Euzet, 1995 | *Coptodon guineensis*  (Günther, 1862) |  |  | AJ920286 | AJ920286 |  | Ivory Coast | Šimková et al. (2006) |
| *Cichlidogyrus amieti*  Birgi & Euzet, 1983 | *Aphyosemion cameronense* (Boulenger, 1903) | AP25 | XXXXXX |  |  |  | Cameroon | This study |
| *Cichlidogyrus amieti*  Birgi & Euzet, 1983 | *Aphyosemion cameronense*  (Boulenger, 1903) | *MRAC:37784* | *KT945076* |  |  |  | Cameroon | Messu Mandeng et al. (2015) |
| *Cichlidogyrus*  *amphoratus*  Pariselle & Euzet, 1996 | *Coptodon guineensis*  (Günther, 1862) | PC51/PC35^1^ | HE792772 | HE792782 | HE792782 |  | Senegal | Mendlová et al. (2012) |
| *Cichlidogyrus arthracanthus*  Paperna, 1960 | *Coptodon guineensis*  (Günther, 1862) | PC60 | HQ010022 | HE792783 | HE792783 |  | Senegal | Mendlová et al. (2010); Mendlová et al. (2012) |
| *Cichlidogyrus attenboroughi* Kmentová, Gelnar, Koblmüller & Vanhove, 2016 | *Benthochromis tricoti* (Poll, 1948) | PB46_CiAt | MH708146 | MH708153 | MH708153 |  | Burundi | Kmentová et al. (2018) |
| *Cichlidogyrus berradae* Pariselle & Euzet, 2003 | *Coptodon rendalli* (Boulenger, 1897) | 182 | XXXXXX | XXXXXX | XXXXXX |  | DR Congo | Jorissen et al. (2021) |
| *Cichlidogyrus berrebii* Pariselle & Euzet, 1994 | *Tylochromis jentinki* (Steindachner, 1894) | AP320 | XXXXXX |  |  |  | Ivory Coast | This study |
| *Cichlidogyrus bilongi* Pariselle & Euzet, 1995 | *Coptodon guineensis* (Günther, 1862) |  |  | AJ920287 | AJ920287 |  | Ivory Coast | Šimková et al. (2006) |
| *Cichlidogyrus brunnensis* Kmentová, Gelnar, Koblmüller & Vanhove, 2016 | *Trematocara unimaculatum* Boulenger, 1901 | PB13_CiBr | MH708144 | MH708152 | MH708152 |  | Burundi | Kmentová et al. (2018) |
| *Cichlidogyrus buescheri* Pariselle & Vanhove, 2015 | *Interochromis loocki* (Poll, 1949) | AP360 | XXXXXX |  |  |  | Zambia | This study |
| *Cichlidogyrus buescheri* Pariselle & Vanhove, 2015 | *Interochromis loocki* (Poll, 1949) | *AP363* | *XXXXXX* | *XXXXXX* | *XXXXXX* |  | Zambia | This study |
| *Cichlidogyrus casuarinus* Pariselle, Muterezi Bukinga & Vanhove, 2015 | *Bathybates minor* Boulenger, 1905 | PB3 | KX007796 | KX007775 | KX007775 | KX007823 | Burundi | Kmentová et al. (2016a) |
| *Cichlidogyrus casuarinus* Pariselle, Muterezi Bukinga & Vanhove, 2015 | *Bathybates minor* Boulenger, 1906 | AP370 | XXXXXX |  |  |  | DR Congo | This study |
| *Cichlidogyrus casuarinus*  Pariselle, Muterezi Bukinga & Vanhove, 2015 | *Bathybates minor* Boulenger, 1906 | *AP371* | *XXXXXX* | *XXXXXX* | *XXXXXX* |  | DR Congo | This study |
| *Cichlidogyrus casuarinus*  Pariselle, Muterezi Bukinga & Vanhove, 2015 | *Bathybates minor* Boulenger, 1906 | *AP373* | *XXXXXX* | *XXXXXX* | *XXXXXX* |  | DR Congo | This study |
| *Cichlidogyrus casuarinus*  Pariselle, Muterezi Bukinga & Vanhove, 2015 | *Bathybates minor* Boulenger, 1906 | *AP374* | *XXXXXX* | *XXXXXX* | *XXXXXX* |  | DR Congo | This study |
| *Cichlidogyrus centesimus*  Vanhove, Volkaert & Pariselle, 2011 | *Ophthalmotilapia ventralis*  (Boulenger, 1898) | AP264 | XXXXXX |  |  |  | DR Congo | This study |
| *Cichlidogyrus* cf. *bychowskyii* | *Hemichromis bimaculatus*  Gill, 1862 | AP171 | XXXXXX |  |  |  | Ivory Coast | This study |
| *Cichlidogyrus cirratus* Paperna, 1964 | *Oreochromis niloticus* (Linnaeus, 1758) | *AP145* | *XXXXXX* |  |  |  | France | This study |
| *Cichlidogyrus cirratus* Paperna, 1964 | *Oreochromis aureus* (Steindachner, 1964) | AP420 | XXXXXX |  |  |  | Morocco | This study |
| *Cichlidogyrus cirratus* Paperna, 1964 | *Oreochromis niloticus* (Linnaeus, 1758) | PC26 | HE792773 | HE792784 | HE792784 |  | Senegal | Mendlová et al. (2012) |
| *Cichlidogyrus consobrini* Jorissen, Pariselle & Vanhove 2017 | *Sargochromis mellandi*  (Boulenger, 1905) | 143 | XXXXXXXX |  |  | XXXXXX | DR Congo | Jorissen et al. (2021); this study |
| *Cichlidogyrus consobrini* Jorissen, Pariselle & Vanhove 2017 | *Sargochromis mellandi*  (Boulenger, 1905) | *144* | *XXXXXXXX* | *XXXXXXXX* | *XXXXXXXX* |  | DR Congo | Jorissen et al. (2021) |
| *Cichlidogyrus cubitus* Dossou, 1982 | *Pelmatolapia mariae* (Boulenger, 1899) | AP164 | XXXXXX |  |  |  | Ivory Coast | Jorissen et al. (2021) |
| *Cichlidogyrus cubitus* Dossou, 1982 | *Coptodon guineensis* (Günther, 1862) | *AP439* | *XXXXXX* |  |  |  | Morocco | Jorissen et al. (2021) |
| *Cichlidogyrus cubitus* Dossou, 1982 | *Coptodon guineensis* (Günther, 1862) | *PC55* | *HQ010037* | *HE792785* | *HE792785* |  | Senegal | Mendlová et al. (2012); Mendlová et al. (2010) |
| *Cichlidogyrus digitatus* Dossou, 1982 | *Pelmatolapia mariae* (Boulenger, 1899) | *AP387* | *XXXXXX* |  |  |  | Cameroon | This study |
| *Cichlidogyrus digitatus* Dossou, 1982 | *Coptodon guineensis* (Günther, 1862) | PC10 | HQ010023 | HE792786 | HE792786 |  | Senegal | Mendlová et al. (2012); Mendlová et al. (2010) |
| *Cichlidogyrus dossoui* Douëllou, 1993 | *Coptodon rendalli* (Boulenger, 1897) | *85* | *XXXXXXXX* | *XXXXXXXX* | *XXXXXXXX* | *XXXXXXXX* | DR Congo | Jorissen et al. (2021) |
| *Cichlidogyrus dossoui* Douëllou, 1993 | *Coptodon rendalli* (Boulenger, 1897) | *AP378* | *XXXXXX* | *XXXXXX* | *XXXXXX* |  | Zambia | This study |
| *Cichlidogyrus dossoui* Douëllou, 1993 | *Coptodon rendalli* (Boulenger, 1897) | AP379 | XXXXXX |  |  |  | Zambia | This study |
| *Cichlidogyrus douellouae* Pariselle, Bilong & Euzet, 2003 | *Sarotherodon galilaeus*  (Linnaeus, 1758) | PC8 | HE792774 | HE792787 | HE792787 |  | Senegal | Mendlová et al. (2012) |
| *Cichlidogyrus dracolemma*  Řehulková, Mendlová & Šimková, 2013 | *Hemichromis letournaeuxi* Sauvage, 1880 | PC6 | HQ010027 | HE792794 | HE792794 |  | Senegal | Mendlová et al. (2012); Mendlová et al. (2010) |
| *Cichlidogyrus ergensi* Dossou, 1982 | *Coptodon guineensis* (Günther, 1862) | PC56 | HQ010038 | HE792788 | HE792788 |  | Senegal | Mendlová et al. (2012); Mendlová et al. (2010) |
| *Cichlidogyrus falcifer* Dossou & Birgi, 1984 | *Hemichromis stellifer* Loiselle, 1979 | *193* | *XXXXXXXX* | *XXXXXXXX* | *XXXXXXXX* | *XXXXXXXX* | DR Congo | Jorissen et al. (2021) |
| *Cichlidogyrus falcifer* Dossou & Birgi, 1984 | *Hemichromis fasciatus*  Peters, 1857 | *AP5* | *XXXXXX* |  |  |  | Ivory Coast | This study |
| *Cichlidogyrus falcifer* Dossou & Birgi, 1984 | *Hemichromis fasciatus*  Peters, 1857 | PC1 | HQ010024 | HE792789 | HE792789 |  | Senegal | Mendlová et al. (2012); Mendlová et al. (2010) |
| *Cichlidogyrus flexicolpos*  Pariselle & Euzet, 1995 | *Coptodon guineensis* (Günther, 1862) |  |  | AJ920283 | AJ920283 |  | Ivory Coast | Šimková et al. (2006) |
| *Cichlidogyrus gallus* Pariselle & Euzet, 1995 | *Coptodon guineensis* (Günther, 1862) |  |  | AJ920285 | AJ920285 |  | Ivory Coast | Šimková et al. (2006) |
| *Cichlidogyrus gillardinae* Muterezi Bukinga, Vanhove, Van Steenberge & Pariselle, 2012 | *Astatotilapia burtoni* (Günther, 1894) | *AP274* | *XXXXXX* |  |  |  | DR Congo | This study |
| *Cichlidogyrus gillardinae* Muterezi Bukinga, Vanhove, Van Steenberge & Pariselle, 2012 | *Astatotilapia burtoni* (Günther, 1894) | AP275 | XXXXXX |  |  |  | DR Congo | This study |
| *Cichlidogyrus halli*  (Price & Kirk, 1967) | *Oreochromis niloticus* (Linnaeus, 1758) | *203* | *XXXXXXXX* | *XXXXXXXX* | *XXXXXXXX* | *XXXXXXXX* | DR Congo | Jorissen et al. (2021) |
| *Cichlidogyrus halli*  (Price & Kirk, 1967) | *Oreochromis niloticus* x *mweruensis* | *C_ha* | *MG973075* | *MG973075* | *MG973075* | *MG970255* | DR Congo | Vanhove et al. (2018) |
| *Cichlidogyrus halli*  (Price & Kirk, 1967) | *Oreochromis niloticus* (Linnaeus, 1758) | *AP97* | *XXXXXX* |  |  |  | Mali | This study |
| *Cichlidogyrus halli*  (Price & Kirk, 1967) | *Sarotherodon galilaeus*  (Linnaeus, 1758) | PC54 | HQ010025 | HE792790 | HE792790 |  | Senegal | Mendlová et al. (2012); Mendlová et al. 2010 |
| *Cichlidogyrus irenae* Gillardin, Vanhove, Pariselle, Huyse & Volckaert, 2012 | Gnathochromis pfefferi  (Boulenger, 1898) | *GnPfC2* |  |  | *KT037169* | *KT037339* | Zambia | Vanhove et al. (2015) |
| *Cichlidogyrus irenae* Gillardin, Vanhove, Pariselle, Huyse & Volckaert, 2012 | *‘Gnathochromis’ pfefferi*  (Boulenger, 1898) | *Kal_Gnpf2_*  *gill3* |  |  | *KT037172* | *KT037340* | Zambia | Vanhove et al. (2015) |
| *Cichlidogyrus irenae* Gillardin, Vanhove, Pariselle, Huyse & Volckaert, 2012 | *‘Gnathochromis’ pfefferi*  (Boulenger, 1898) | *PB43_GnPf1* | *MH708145* | *KT692939* | *KT692939* |  | Burundi | Kmentová et al. (2016b); Kmentová et al. (2018) |
| *Cichlidogyrus irenae* Gillardin, Vanhove, Pariselle, Huyse & Volckaert, 2012 | *‘Gnathochromis’ pfefferi*  (Boulenger, 1898) | *T08_Kal_*  *Gnpf1_gill1* |  | *XXXXXX* | *XXXXXX* |  | Zambia | Vanhove et al. (2015); this study |
| *Cichlidogyrus irenae* Gillardin, Vanhove, Pariselle, Huyse & Volckaert, 2012 | *‘Gnathochromis’ pfefferi*  (Boulenger, 1898) | T08_Kal_  Gnpf1_gill3 |  | XXXXXX | XXXXXX |  | Zambia | Vanhove et al. (2015); this study |
| *Cichlidogyrus kothiasi* Pariselle & Euzet, 1994 | *Tylochromis jentinki* (Steindachner, 1894) | AP325 | XXXXXX |  |  |  | Ivory Coast | This study |
| *Cichlidogyrus longicirrus*  Paperna, 1965 | *Hemichromis elongatus* (Guichenot, 1861) | AP58 | XXXXXX |  |  |  | Cameroon | This study |
| *Cichlidogyrus longicirrus*  Paperna, 1965 | *Hemichromis fasciatus* Peters, 1857 | *PC37* | *HQ010026* | *HE792791* | *HE792791* |  | Senegal | Mendlová et al. (2012); Mendlová et al. (2010) |
| *Cichlidogyrus mbirizei* Muterezi Bukinga, Vanhove, Van Steenberge & Pariselle, 2012 | *Oreochromis niloticus* x *mweruensis* | *C_mb* | *MG973076* | *MG973076* | *MG973076* | *MG970257* | DR Congo | Vanhove et al. (2018) |
| *Cichlidogyrus mbirizei* Muterezi Bukinga, Vanhove, Van Steenberge & Pariselle, 2012 | *Oreochromis tanganicae* (Günther, 1894) | AP433 | XXXXXX |  |  |  | DR Congo | This study |
| *Cichlidogyrus nageus* Řehulková, Mendlová & Šimková, 2013 | *Sarotherodon galilaeus* (Linnaeus, 1758) | PC13 | HQ010028 | HE792795 | HE792795 |  | Senegal | Mendlová et al. (2012); Mendlová et al. (2010) |
| *Cichlidogyrus nandidae* Birgi & Lambert, 1986 | *Polycentropsis abbreviata* (Boulenger, 1901) | AP23 | XXXXXX |  |  |  | Cameroon | This study |
| *Cichlidogyrus njinei* Pariselle, Bilong & Euzet, 2003 | *Sarotherodon galilaeus*  (Linnaeus, 1758) | PC9/PC86^1^ | HE792775 | HE792792 | HE792792 |  | Senegal | Mendlová et al. (2012) |
| *Cichlidogyrus nshomboi* Muterezi Bukinga, Vanhove, Van Steenberge & Pariselle, 2012 | *Boulengerochromis microlepis* (Boulenger, 1899) | *AP368* | *XXXXXX* | *XXXXXX* | *XXXXXX* |  | DR Congo | This study |
| *Cichlidogyrus nshomboi* Muterezi Bukinga, Vanhove, Van Steenberge & Pariselle, 2012 | *Boulengerochromis microlepis* (Boulenger, 1899) | AP369 | XXXXXX |  |  |  | DR Congo | This study |
| *Cichlidogyrus papernastrema*  Price, Peebles & Bamford, 1969 | *Tilapia sparrmanii* Smith, 1840 | *80* | *XXXXXXXX* |  |  | *XXXXXX* | DR Congo | Jorissen et al. (2021); this study |
| *Cichlidogyrus papernastrema*  Price, Peebles & Bamford, 1969 | *Coptodon rendalli* (Boulenger, 1897) | AP381 | XXXXXX |  |  |  | Zambia | This study |
| *Cichlidogyrus philander* Douëllou, 1993 | *Pseudocrenilabrus philander*  (Weber, 1897) | CP11 | MG279691 |  | MG250200 | MG288503 | South Africa | Igeh et al. (2017) |
| *Cichlidogyrus pouyaudi* Pariselle & Euzet 1994 | *Tylochromis intermedius* (Boulenger, 1916) | PC69 | HQ010039 | HE792793 | HE792793 |  | Senegal | Mendlová et al. (2012; Mendlová et al.) |
| *Cichlidogyrus quaestio* Douëllou, 1993 | *Coptodon rendalli* (Boulenger, 1897) | *83* | *XXXXXXXX* |  |  | *XXXXXXXX* | DR Congo | Jorissen et al. (2021) |
| *Cichlidogyrus quaestio* Douëllou, 1993 | *Coptodon rendalli* (Boulenger, 1897) | 74 | XXXXXXXX |  |  |  | DR Congo | Jorissen et al. (2021) |
| *Cichlidogyrus sanseoi* Pariselle & Euzet 2004 | *Hemichromis fasciatus* Peters, 1857 | AP52 | XXXXXX |  |  |  | Senegal | This study |
| *Cichlidogyrus schreyenbrichardorum* Pariselle & Vanhove, 2015 | *Interochromis loocki* (Poll, 1949) | *AP358* | *XXXXXX* |  |  |  | Zambia | This study |
| *Cichlidogyrus schreyenbrichardorum* Pariselle & Vanhove, 2015 | *Interochromis loocki* (Poll, 1949) | AP362 | XXXXXX | XXXXXX | XXXXXX |  | Zambia | This study |
| *Cichlidogyrus sclerosus* Paperna & Thurston, 1969 | *Oreochromis niloticus* (Linnaeus, 1758) | *220* | *XXXXXXXX* |  |  | *XXXXXXXX* | DR Congo | Jorissen et al. (2021) |
| *Cichlidogyrus sclerosus* Paperna & Thurston, 1969 | *Oreochromis mossambicus* (Peters, 1852) | *AP249* | *XXXXXX* |  |  |  | DR Congo | This study |
| *Cichlidogyrus sclerosus* Paperna & Thurston, 1969 | *Oreochromis niloticus* (Linnaeus, 1758) | *AP124* | *XXXXXX* |  |  |  | France | This study |
| *Cichlidogyrus sclerosus* Paperna & Thurston, 1969 | *Oreochromis mossambicus* (Peters, 1852) | *AP149* | *XXXXXX* |  |  |  | Mozambique | This study |
| *Cichlidogyrus sclerosus* Paperna & Thurston, 1969 | *Oreochromis niloticus* (Linnaeus, 1758) |  | DQ157660 | DQ537359 | DQ537359 |  | China | Wu et al. (2006); Wu et al. (2007) |
| *Cichlidogyrus teugelsi* Pariselle & Euzet 2004 | *Hemichromis fasciatus*  Peters, 1857 | AP13 | XXXXXX |  |  |  | Ivory Coast | This study |
| *Cichlidogyrus thurstonae* Ergens, 1981 | *Oreochromis niloticus* (Linnaeus, 1758) | 214 | XXXXXXXX | XXXXXXXX | XXXXXXXX | XXXXXXXX | DR Congo | Jorissen et al. (2021) |
| *Cichlidogyrus tiberianus* Paperna, 1960 | *Coptodon rendalli* (Boulenger, 1897) | *AP382* | *XXXXXX* | *XXXXXX* | *XXXXXX* |  | Zambia | This study |
| *Cichlidogyrus tiberianus* Paperna, 1960 | *Coptodon guineensis* (Günther, 1862) | PC12/PC15^1^ | HE792776 | HE792796 | HE792796 |  | Senegal | Mendlová et al. (2012) |
| *Cichlidogyrus tilapiae* Paperna, 1960 | *Oreochromis niloticus* (Linnaeus, 1758) | *208* | *XXXXXXXX* | *XXXXXXXX* | *XXXXXXXX* | *XXXXXXXX* | DR Congo | Jorissen et al. (2021) |
| *Cichlidogyrus tilapiae* Paperna, 1960 | *Hemichromis fasciatus*  Peters, 1857 | PC43 | HQ010029 | HE792797 | HE792797 |  | Senegal | Mendlová et al. (2012); Mendlová et al. (2010) |
| *Cichlidogyrus vandekerkhovei* Vanhove, Volkaert & Pariselle, 2011 | *Ophthalmotilapia ventralis*  (Boulenger, 1898) | AP266 | XXXXXX |  |  |  | DR Congo | This study |
| *Cichlidogyrus vandekerkhovei* Vanhove, Volkaert & Pariselle, 2011 | *Ophthalmotilapia ventralis*  (Boulenger, 1898) | *AP267* | *XXXXXX* |  |  |  | DR Congo | This study |
| *Cichlidogyrus vealli* Pariselle & Vanhove, 2015 | *Interochromis loocki* (Poll, 1949) | *AP356* | *XXXXXX* | *XXXXXX* | *XXXXXX* |  | Zambia | This study |
| *Cichlidogyrus vealli* Pariselle & Vanhove, 2015 | *Interochromis loocki* (Poll, 1949) | AP357 | XXXXXX | XXXXXX | XXXXXX |  | Zambia | This study |
| *Cichlidogyrus yanni* Pariselle & Euzet, 1996 | *Sarotherodon melanotheron* Rüppel, 1852 | *AP155* | *XXXXXX* |  |  |  | France | This study |
| *Cichlidogyrus yanni* Pariselle & Euzet, 1996 | *Coptodon guineensis* (Günther, 1862) | PC14 | HE792777 | HE792798 | HE792798 |  | Senegal | Mendlová et al. (2012) |
| *Cichlidogyrus zambezensis*  Douëllou, 1993 | *Serranochromis macrocephalus* (Boulenger, 1899) | 269 | XXXXXXXX | XXXXXXXX | XXXXXXXX | XXXXXXXX | DR Congo | Jorissen et al. (2021) |
| *Cichlidogyrus zambezensis*  Douëllou, 1993 | *Serranochromis macrocephalus* (Boulenger, 1899) | *AP375* | *XXXXXX* | *XXXXXX* | *XXXXXX* |  | Zambia | This study |
| *Cichlidogyrus zambezensis*  Douëllou, 1993 | *Serranochromis macrocephalus* (Boulenger, 1899) | *AP376* | *XXXXXX* |  |  |  | Zambia | This study |
| *Scutogyrus bailloni* Pariselle & Euzet, 1995 | *Sarotherodon galilaeus*  (Linnaeus, 1758) | *AP133* | *XXXXXX* |  |  |  | Ivory Coast | This study |
| *Scutogyrus bailloni* Pariselle & Euzet, 1995 | *Sarotherodon galilaeus*  (Linnaeus, 1758) | AP1 | HE792778 | HE792799 | HE792799 |  | Ivory Coast | Mendlová et al. (2012) |
| *Scutogyrus gravivaginus* (Paperna & Thurston 1969) | *Oreochromis mweruensis* Trewavas, 1983 | 65 | XXXXXXXX |  |  | XXXXXX | DR Congo | Jorissen et al. (2021); this study |
| *Scutogyrus longicornis* (Paperna & Thurston 1969) | *Oreochromis niloticus* (Linnaeus, 1758) | *277* | *XXXXXXXX* |  |  | *XXXXXX* | DR Congo | Jorissen et al. (2021); this study |
| *Scutogyrus longicornis* (Paperna & Thurston 1969) | *Oreochromis niloticus* (Linnaeus, 1758) | AP99 | XXXXXX |  |  |  | Mali | This study |
| *Scutogyrus longicornis* (Paperna & Thurston 1969) | *Oreochromis niloticus* (Linnaeus, 1758) | *PC105* | *HQ010035* | *HE792800* | *HE792800* |  | Senegal | Mendlová et al. (2012); Mendlová et al. (2010) |
| *Scutogyrus minus* (Dossou, 1982) | *Sarotherodon melanotheron* Rüppel, 1852 | AP88 | XXXXXX |  |  |  | Ivory Coast | This study |
| *Scutogyrus minus* (Dossou, 1982) | *Sarotherodon melanotheron* Rüppel, 1852 | *AP2/AP123^1^* | *HE792779* | *HE792801* | *HE792801* |  | Ivory Coast | Mendlová et al. (2012) |
| *Scutogyrus vanhovei* Pariselle, Bitja Nyom & Bilong Bilong, 2013 | *Pelmatolapia mariae* (Boulenger, 1899) | AP385 | XXXXXX |  |  |  | Cameroon | This study |

^1^ isolate numbers differ on GenBank but sequences were concatenated in Mendlová et al. (2012).

**Appendix 4.** Substitution models of molecular evolution and partitions for Bayesian inference (BI) and maximum likelihood estimation (ML) of phylogeny of species of *Cichlidogyrus* and *Scutogyrus*. Models include the general time reversible model (GTR), the Kimura 1980 model (K80), the transitional model 3 with unequal base frequencies (TIM3e), the Tamura-Nei model (TN), and the three-parameter model 2 (TPM2) plus empirical base frequencies (+ F), a proportion of invariable sites (+ I), a discrete Γ model with four rate categories (Γ4), or a FreeRate model with three categories (+ R3). For model specification see the IQ-TREE ModelFinder manual (Kalyaanamoorthy et al., 2017).

| **Partition** | **Bayesian inference (BI)** | **Maximum likelihood estimation (ML)** |
| --- | --- | --- |
| 28S rDNA | GTR + F + I + Γ4 | TIM3e + F + R3 |
| 18S rDNA | K80 + I + Γ4 | K80 + I + Γ4 |
| ITS rDNA | GTR + F + Γ4 | TPM2 + F + Γ4 |
| CO1 mtDNA | GTR + F + I + Γ4 | TN + F + I + Γ4 |

**Appendix 5.** Species groups of *Cichlidogyrus* with species included, species potentially included, and the respective host ranges reported in the taxonomic literature including host species of candidate species.

| **Group** | **Species included** | **Candidate species** | **Hosts** | **Hosts of candidate species** |
| --- | --- | --- | --- | --- |
| *Bulb* | *C. papernastrema* Pariselle & Euzet, 1994  *C. philander* Pariselle & Euzet, 1994  *C. zambezensis* Pariselle & Euzet, 1994 | *C. maeander* Geraerts & Muterezi Bukinga in Geraerts *et al*., 2020  *C. pseudozambezensis* Geraerts & Muterezi Bukinga in Geraerts *et al*., 2020. | **Coptodonini**  *Coptodon rendalli* (Boulenger, 1897)  **Haplochromini**  *Pseudocrenilabrus philander* (Weber, 1897)  *Sargochromis mellandi* (Boulenger, 1905)  *Serranochromis angusticeps* (Boulenger, 1907)  *Serranochromis jallae* (Boulenger, 1896)  *Serranochromis macrocephalus* (Boulenger, 1899)  *Serranochromis thumbergi* (Castelnau, 1861)  **Oreochromini**  *Oreochromis mortimeri* (Trewavas, 1983)  *O.* *mweruensis* (Trewavas, 1983)*,*  **Tilapiini**  *Tilapia sparrmanii* Smith, 1840. | **Coptodonini**  *Coptodon camerunensis* (Lönnberg, 1903)  *C. dageti* (Thy van den Audenaerde, 1971)  *C. guineensis* (Günther, 1862)  *C. zillii* (Gervais, 1848)  **Haplochromini**  *Sargochromis codringtonii* (Boulenger, 1905)  *Serranochromis* cf. *macrocephalus* sensu Geraerts et al. (2020)  ‘*Orthochromis*’ sp. ‘Lomami’ sensu Geraerts et al. (2020)  **Oreochromini**  *Sarotherodon melanotheron* Rüppell, 1852  *Sarotherodon occidentalis* (Daget, 1962) |
| *Cop* | *C. berradae* Pariselle & Euzet, 2003  *C. digitatus* Dossou, 1982  *C. quaestio* Douëllou, 1993  *C. yanni* Pariselle & Euzet, 1996 | *C. berminensis* Pariselle, Bitja Nyom, & Bilong Bilong, 2013  *C. halinus* Paperna, 1969  *C. nuniezi* Pariselle & Euzet, 1998  *C. reversati* Pariselle & Euzet, 2003. | **Coptodonini**  *Coptodon camerunensis* (Lönnberg, 1903)  *C. coffea* (Thys van den Audenaerde, 1970)  *C. dageti* (Thys van den Audenaerde, 1971)  *C. discolor* (Günther, 1903)  *C. guineensis* (Günther, 1862)  *C. louka* (Thys van den Audenaerde, 1969)  *C. rendalli* (Boulenger, 1897)  *C. tholloni* (Sauvage, 1884)  *C. walteri* (Thys van den Audenaerde, 1968)  *C. zillii* (Gervais, 1848)  **Gobiocichlini**  *“Tilapia” brevimanus* Boulenger, 1911  **Haplochromini**  *Pseudocrenilabrus multicolor* (Schöller, 1903)  *Sargochromis codringtonii* (Boulenger, 1908)  *Serranochromis macrocephalus* (Boulenger, 1899)  *Thoracochromis wingatii* (Boulenger, 1902)*,*  **Oreochromini**  *Oreochromis niloticus* (Linnaeus, 1758)  **Pelmatolapiini**  *Pelmatolapia cabrae* (Boulenger, 1899)  *P. mariae* (Boulenger, 1899)  **Tilapiini**  *Tilapia sparrmanii* Smith, 1840 | **Coptodonini**  *Coptodon bakossiorum* (Stiassny, Schliewen, & Dominey, 1992)  *C. bemini* (Thys van den Audenaerde, 1972)  *C. gutturosa* (Stiassny, Schliewen, & Dominey, 1992)  *C. thysi* (Stiassny, Schliewen, & Dominey, 1992)  **Heterotilapiini**  *Heterotilapia buttikoferi* (Hubrecht, 1881)  *H. cessiana* (Thys van den Audenaerde, 1968)  **Oreochromini**  *Sarotherodon melanotheron* Rüppel, 1852 |
| *CPO* | *C. aegypticus* Ergens, 1981  *C. agnesi* Pariselle & Euzet, 1995  *C. arthracanthus* Paperna, 1960  *C. bilongi* Pariselle & Euzet, 1995  *C. cubitus* Dossou, 1982  *C. dossoui* Douëllou, 1993  *C. douellouae* Pariselle, Bilong Bilong, & Euzet, 2003  *C. ergensi* Dossou, 1982  *C. flexicolpos* Pariselle & Euzet, 1995  *C. gallus* Pariselle & Euzet, 1995  *C. thurstonae* Ergens, 1981  *C. tiberianus* Paperna, 1960 | *C. anthemocolpos* Dossou, 1982  *C. bonhommei* Pariselle & Euzet, 1998  *C. bouvii* Pariselle & Euzet, 1997  *C. gillesi* Pariselle, Bitja Nyom, & Bilong Bilong, 2013  *C. guirali* Pariselle & Euzet, 1997  *C. hemi* Pariselle & Euzet, 1998  *C. kouassii* N’Douba, Thys van den Audenaerde, & Pariselle, 1997  *C. legendrei* Pariselle & Euzet, 2003;  *C. lemoallei* Pariselle & Euzet, 2003  *C. louipaysani* Pariselle & Euzet, 1995  *C. microscutus* Pariselle & Euzet, 1996  *C. ouedraogoi* Pariselle & Euzet, 1996  *C. paganoi* Pariselle & Euzet, 1997  *C. testificatus* Dossou, 1982  *C. vexus* Pariselle & Euzet, 1995 | **Coptodonini**  *Coptodon bakossiorum* (Stiassny, Schliewen, & Dominey, 1992)  *C. camerunensis* (Lönnberg, 1903)  *C. coffea* (Thys van den Audenaerde, 1970)  *C. dageti* (Thys van den Audenaerde, 1971)  *C. deckerti* (Thys van den Audenaerde, 1967)  *C. guineensis* (Günther, 1862)  *C. gutturosa* (Stiassny, Schliewen, & Dominey, 1992)  *C. kottae* (Lönnberg, 1904)  *C. louka* (Thys van den Audenaerde, 1969)  *C. rendalli* (Boulenger, 1897)  *C. tholloni* (Sauvage, 1884)  *C. walteri* (Thys van den Audenaerde, 1968)  *C. zillii* (Gervais, 1848)*,*  **Gobiocichlini**  *“Tilapia” brevimanus* Boulenger, 1911  *“Tilapia” busumana* (Günther, 1903)  **Heterotilapiini**  *Heterotilapia buttikoferi* (Hubrecht, 1881)  **Oreochromini**  *Oreochromis aureus* (Steindachner, 1864)  *O. esculentus* (Graham, 1928)  *O. mortimeri* (Trewavas, 1966)  *O. mossambicus* (Peters, 1852)  *O. mweruensis* Trewavas, 1983  *O. niloticus* (Linneaus, 1758)  *O. variabilis* (Boulenger, 1906)  *Sarotherodon galilaeus* (Linneaus, 1785)  **Pelmatolapiini**  *Pelmatolapia cabrae* (Boulenger, 1899)  *P. mariae* (Boulenger, 1899)  **Tilapiini**  *Tilapia sparrmanii* Smith, 1840  **Haplochromini**  *Haplochromis longirostris* (Hilgendorf, 1888) and *Serranochromis macrocephalus* (Boulenger, 1899) (Paperna and Thurston 1969; Douëllou 1993) not verified.  **Invasive**  *Cichlidogyrus thurstonae* on the Malagasy cichlids *Paretroplus lamenabe* Sparks, 2008 and *Ptychochromis inornatus* Sparks, 2002 (Šimková et al., 2019)*. Cichlidogyrus arthracanthus* and *C. tiberianus* on Western Asian haplochromine and oreochromine cichlids including *Astatotilapia flaviijosephi* (Lortet, 1883)*, Tristramella sacra* (Günther, 1865)*,* and *Tristramella simonis* (Günther, 1864) (Paperna 1960). However, *T. sacra* is now considered extinct (Goren, 2014). | **Oreochromini**  *Sarotherodon occidentalis* (Daget, 1962) |
| *EAR* – *CT* subgroup | *C. brunnensis* Kmentová, Gelnar, Koblmüller & Vanhove, 2016  *C. milangelnari* Rahmouni, Vanhove, & Šimková, 2017 | *C. evikae* Rahmouni, Vanhove, & Šimková, 2017  *C. jeanloujustinei* Rahmouni, Vanhove, & Šimková, 2017  *C. koblmuelleri* Rahmouni, Vanhove, & Šimková, 2018 | **Cyprichromini**  *Cyprichromis microlepidotus* (Poll, 1956)  **Trematocarini**  *Trematocara unimaculatum* Boulenger, 1901 | **Eretmodini**  *Eretmodus marksmithii* Burgess, 2012 *Tanganicodus irsacae* Poll, 1950 |
| *EAR* – *Heel* subgroup | *C. casuarinus* Pariselle, Muterezi Bukinga, & Vanhove, 2015  *C. centesimus* Vanhove, Volkaert, & Pariselle, 2011  *C. nshomboi* Muterezi Bukinga, Vanhove, Van Steenberge, & Pariselle, 2012 | *C. aspiralis* Rahmouni, Vanhove, & Šimková, 2017  *C. habluetzeli* Rahmouni, Vanhove, & Šimková, 2018  *C. pseudoaspiralis* Rahmouni, Vanhove, & Šimková, 2017. | **Bathybatini**  *Bathybates fasciatus* Boulenger, 1901  *B. hornii* Steindachner, 1911  *B. leo* Poll, 1965  *B. minor* Boulenger, 1906  *B. vittatus* Boulenger, 1914  *Hemibates stenosoma* (Boulenger, 1901)  **Boulengerochromini**  *Boulengerochromis microlepis* (Boulenger, 1899)  **Ectodini**  *Ophthalmotilapia boops* (Boulenger, 1901)  *O. nasuta* (Poll & Matthes, 1962)  *O. ventralis* (Boulenger, 1898) | **Ectodini**  *Aulonocranus dewindti* (Boulenger, 1899)  **Cyphotilapiini**  *Cyphotilapia frontosa* (Boulenger, 1906) |
| *EAR*–*Troph* subgroup | *C. buescheri* Pariselle & Vanhove, 2015  *C. irenae* Gillardin, Vanhove, Pariselle, Snoeks, Huyse, & Volkaert, 2011  *C. schreyenbrichardorum* Pariselle & Vanhove, 2015  *C. vealli* Pariselle & Vanhove, 2015. | *C. antoineparisellei* Rahmouni, Vanhove, & Šimková, 2018  *C. banyankimbonai* Pariselle & Vanhove, 2015  *C. frankwillemsi* Pariselle & Vanhove, 2015  *C. franswittei* Pariselle & Vanhove, 2015  *C. georgesmertensi* Pariselle & Vanhove, 2015  *C. gistelincki* Gillardin, Vanhove, Pariselle, Huyse, & Volkaert, 2012  *C. masilyai* Rahmouni, Vanhove, & Šimková, 2018  *C. muterezii* Pariselle & Vanhove, 2015; *C. raeymaekersi* Pariselle & Vanhove, 2015  *C. salzburgeri* Rahmouni, Vanhove, & Šimková, 2018  *C. steenbergei* Gillardin, Vanhove, Pariselle, Huyse, & Volkaert, 2012. | **Haplochromini (Tropheini)**  ‘*Gnathochromis*’ *pfefferi* (Boulenger, 1898)  *Interochromis loocki* (Poll, 1949) | **Haplochromini (Tropheini)**  ‘*Ctenochromis*’ *horei* (Günther, 1894)  *Limnotilapia dardennii* (Boulenger, 1899)  *Simochromis diagramma* (Günther, 1894)  ‘*Petrochromis*’ *orthognathus* Matthes, 1959  *Petrochromis trewavasae* Poll, 1948  *Pseudosimochromis babaulti* (Pellegrin, 1927)  *P. curvifrons* (Poll, 1942)  *P. marginatus* (Poll, 1956)  *P. pleurospilus* (Nelissen, 1978). |

| *EAR* – Other species | *C. attenboroughi* Kmentová, Gelnar, Koblmüller & Vanhove, 2016  *C. consobrini* Jorissen, Pariselle & Vanhove in Jorissen et al. 2018b  *C. gillardinae* Muterezi Bukinga, Vanhove, Van Steenberge, & Pariselle, 2012. | *C. adkoningsi* Rahmouni, Vanhove, & Šimková, 2018  *C. discophonum* Rahmouni, Vanhove, & Šimková, 2017  *C. glacicremoratus* Rahmouni, Vanhove, & Šimková, 2017  *C. haplochromii* Paperna & Thurston, 1969  *C. longipenis* Paperna & Thurston, 1969  *C. makasai* Vanhove, Volkaert, & Pariselle, 2011  *C. rectangulus* Rahmouni, Vanhove, & Šimková, 2017  *C. sturmbaueri* Vanhove, Volkaert, & Pariselle, 2011  *C. vandekerkhovei* Vanhove, Volkaert, & Pariselle, 2011 | **Benthochromini**  *Benthochromis horii* Takahashi, 2008  **Ectodini**  *Ophthalmotilapia boops* (Boulenger, 1901)  *O. nasuta* (Poll & Matthes, 1962)  *O. ventralis* (Boulenger, 1898)  **Haplochromini**  *Astatotilapia burtoni* (Günther, 1894)  ‘*Orthochromis*’ *katumbii* Schedel, Vreven, Manda, Abwe, Manda, & Schliewen, 2018  *Sargochromis mellandi* (Boulenger, 1905).) | **Ectodini**  *Aulonocranus dewindti* (Boulenger, 1899)  *Cardiopharynx schoutedeni* Poll, 1942  *Tanganicodus irsacae* Poll, 1950  **Cyphotilapiini**  *Cyphotilapia frontosa* (Boulenger, 1906)  **Haplochromini**  *Astatoreochromis alluaudi* Pellegrin, 1904  *Haplochromis aeneocolor* Greenwood, 1973  *H. angustifrons* Boulenger, 1914  *H. bicolor* Boulenger, 1906  *H. degeni* (Boulenger, 1906)  *H. elegans* Trewavas, 1933  *H. guiarti* (Pellegrin, 194)  *H. longirostris* (Hilgendorf, 1888)*, H. macrognathus* Regan, 1922  *H. nigripinnis* Regan, 1921  *H. nubilus* (Boulenger, 1906)  *H. obesus* (Boulenger, 1906)  *H. obliquidens* (Hilgendorff, 1888)  *H. petronius* Greenwood, 1973  *H. retrodens* (Hilgendorff, 1888)  *H. schubotzi* Boulenger, 1914  *H. squamipinnis* Regan, 1921  *Pharyngochromis darlingi* (Boulenger, 1911)  *Pseudocrenilabrus multicolor* (Schöller, 1903)  *Thoracochromis wingatii* (Boulenger, 1902)  **Possibly invasive or misreported**  *C. bifurcatus* and *C. haplochromii* on *Oreochromis aureus* (Steindachner, 1864)*, O. leucostictus* (Trewavas, 1933)*, O. niloticus* (Linnaeus, 1758), and *Coptodon zillii* Gervais, 1848 (Paperna, 1960, 1979; Thurston, 1970; Jiménez-Garcia et al., 2001). |
| --- | --- | --- | --- | --- |
| *Halli* | Previous studies suggest that *Cichlidogyrus halli* Price & Kirk, 1967 is species complex (Douëllou 1993; Jorissen et al. 2018b) including following morphotypes:  *C. halli* Price & Kirk, 1967  *C. halli* ‘morphotype 2’ sensu Jorissen et al. (2018b). | **–** | **Coptodonini**  *Coptodon guineensis* (Günther, 1862)  *C. zillii* (Gervais, 1848)  **Oreochromini**  *Oreochromis aureus* (Steindachner, 1864)  *O. esculentus* (Graham, 1928)  *O. leucostictus* (Trewavas, 1933)  *O. mortimeri* (Trewavas, 1966)  *O. mossambicus* (Peters, 1852)  *O. mweruensis* Trewavas, 1983  *O. niloticus* (Linnaeus, 1758)  *O. shiranus* Boulenger, 1897  *O. spilurus* (Günther, 1894)  *O. tanganicae* (Günther, 1894)  *O. variabilis* (Boulenger, 1906)  *Sarotherodon galilaeus* (Linnaeus, 1758)  *S. melanotheron* Rüppel, 1852  *S. occidentalis* (Daget, 1962)  **Haplochromini**  *Serranochromis macrocephalus* (Boulenger, 1899) | **–** |
| *Hemi* | *Cichlidogyrus amieti* Birgi & Euzet, 1983  *C.* cf. *bychowskii* sensu Paperna, 1965 (see Jorissen et al. 2018a)  *C. dracolemma* Řehulková, Mendlová, & Šimková, 2013  *C. falcifer* Dossou & Birgi, 1984  *C. longicirrus* Paperna, 1965  *C. nandidae* Birgi & Lambert, 1986  *C. sanseoi* Pariselle & Euzet, 2004  *C. teugelsi* Pariselle & Euzet, 2004 | *C. calycinus* Kusters, Jorissen, Pariselle, & Vanhove in Jorissen et al. 2018a  *C. dageti* Dossou & Birgi, 1984  *C. dionchus* Paperna & Thurston, 1969  *C. euzeti* Dossou & Birgi, 1984  *C. inconsultans* Birgi & Lambert, 1987  *C. kmentovae* Jorissen, Pariselle, & Vanhove in Jorissen et al. 2018a  *C. polyenso* Jorissen, Pariselle, & Vanhove in Jorissen et al. 2018a. | **Hemichromini**  *Hemichromis bimaculatus* Gill, 1862  *H. elongatus* (Guichenot, 1861)  *H. fasciatus* Peters, 1857  *H. letourneuxi* Sauvage, 1880  **Chromidotilapiini**  *Chromidotilapia guntheri* (Sauvage, 1882)  **Nothibranchiidae**  *Aphyosemion cameronense* (Boulenger, 1903)  *A. exiguum* (Boulenger, 1911)  *A. obscurum* (Ahl, 1924)  **Polycentridae**  *Polycentropsis abbreviata* Boulenger, 1901 | **Hemichromini**  *Hemichromis stellifer* Loiselle, 1979 |
| *Oreo1* | *C. acerbus* Dossou, 1982  *C. cirratus* Paperna, 1964  *C. mbirizei* Muterezi Bukinga, Vanhove, Van Steenberge, & Pariselle, 2012  *C. nageus* Řehulková, Mendlová, & Šimková, 2013  *C. njinei* Pariselle, Bilong Bilong, & Euzet, 2003 | *C. giostrai* Pariselle, Bilong Bilong, & Euzet, 2003  *C. mvogoi* Pariselle, Bitja Nyom, & Bilong Bilong, 2014  *C. lagoonaris* Paperna, 1969  *C. ornatus* Pariselle & Euzet 1996  *C. slembroucki* Pariselle & Euzet, 1998 | **Coptodonini**  *Coptodon guineensis* (Günther, 1862)  *C. zillii* (Gervais, 1848)*,*  **Oreochromini**  *Oreochromis esculentus* (Graham, 1928)  *O. mossambicus* (Peters, 1852)  *O. mweruensis* Trewavas, 1983  *O. niloticus* (Linnaeus, 1758)  *O. tanganicae* (Günther, 1894)  *O. variabilis* (Boulenger, 1906)  *Sarotherodon galilaeus* (Linnaeus, 1758)  *S. melanotheron* Rüppel, 1852. | **Coptodonini**  *Coptodon camerunensis* (Lönnberg, 1903)  *C. dageti* (Thys van den Audenaerde, 1967)  **Oreochromini**  *Sarotherodon caudomarginatus* (Boulenger, 1916)  *S. mvogoi* (Thys van den Audenaerde, 1965)  **Heterotilapiini**  *Heterotilapia buttikoferi* (Hubrecht, 1881) |
| *Oreo2* | *C. amphoratus* Pariselle & Euzet, 1996  *C. sclerosus* Paperna & Thurston, 1969 | **–** | **Coptodonini**  *Coptodon guineensis* (Günther, 1862)  *C. louka* (Thys van den Audenaerde, 1969)  *C. rendalli* (Boulenger, 1897)  *C. tholloni* (Sauvage, 1884)  *C. zillii* (Gervais, 1848)  **Haplochromini**  *Serranochromis macrocephalus* (Boulenger, 1899)  **Oreochromini**  *Oreochromis aureus* (Steindachner, 1864)  *O. leucostictus* (Trewavas, 1933)  *O. mortimeri* (Trewavas, 1983)  *O. mossambicus* (Peters, 1852)  *O. mweruensis* (Trewavas, 1983)  *O. niloticus* (Linnaeus, 1758)  *O. spilurus* (Günther, 1894)  *O. urolepis* (Norman, 1922)  *Sarotherodon galilaeus* (Linnaeus, 1758)  **Invasive**  *Cichlidogyrus sclerosus* on *Vieja fenestrata* (Günther, 1860) (Cichlidae: Cichlasomatini) (Jiménez-Garcia et al. 2001) | **–** |
| *Tylo* | *Cichlidogyrus berrebii* Pariselle & Euzet, 1994  *C. kothiasi* Pariselle & Euzet, 1994  *C. pouyaudi* Pariselle & Euzet, 1994 | *Cichlidogyrus bixlerzavalai* Jorissen, Pariselle, & Vanhove, 2018  *C. chrysopiformis* Pariselle, Bitja Nyom, & Bilong Bilong, 2014  *C. dijetoi* Pariselle, Bitja Nyom, & Bilong Bilong, 2014  *C. mulimbwai* Muterezi Bukinga, Vanhove, Van Steenberge, & Pariselle, 2012  *C. muzumanii* Muterezi Bukinga, Vanhove, Van Steenberge, & Pariselle, 2012  *C. omari* Jorissen, Pariselle, & Vanhove, 2018  *C. sergemorandi* Rahmouni, Vanhove, & Šimková, 2018  *C. sigmocirrus* Pariselle, Bitja Nyom, & Bilong Bilong, 2014 | **Tylochromini**  *Tylochromis intermedius* (Boulenger, 1916)  *T. jentinki* (Steindachner, 1894). | **Tylochromini**  *Tylochromis polylepis* (Boulenger, 1900)  *T. praecox* Stiassny, 1989  *T. sudanensis* Daget, 1954. |
| *Scutogyrus* | *Scutogyrus bailloni* Pariselle & Euzet, 1995  *S. gravivaginus* (Pariselle & Thurston, 1969)  *S. longicornis* (Paperna & Thurston ,1969)  *S. minus* (Dossou, 1982)  *S. vanhovei* Pariselle, Bitja Nyom, & Bilong Bilong, 2014 | *S. chikii* Pariselle & Euzet, 1995  *S. ecoutini* Pariselle & Euzet, 1995 | **Oreochromini**  *Oreochromis aureus* (Steindachner, 1864)  *O. leucostictus* (Trewavas, 1933)  *O. mortimeri* (Trewavas, 1966)  *O. mweruensis* Trewavas, 1983  *O. niloticus* (Linnaeus, 1758)  *O. tanganicae* (Günther, 1894)  *O. variabilis* (Boulenger, 1906)  *Sarotherodon galilaeus* (Linnaeus, 1758)  *S. melanotheron* Rüppel, 1852  **Pelmatolapiini**  *Pelmatolapia mariae* (Boulenger, 1899)  **Invasive**  *S. longicornis* on *Vieja fenestrata* (Günther, 1860) (Jiménez-Garcia et al. 2001) | **Oreochromini**  *Oreochromis mossambicus* (Peters, 1852)  *Sarotherodon occidentalis* (Daget, 1965). |

**Appendix 6.** Ancestral states of continuous characters inferred from a literature survey and used for character mapping. Character IDs refer to numbers in Appendix 2.

| **Character ID** | ***Tylo*** | ***Oreo2*** | ***Halli*** | ***CPO*** | ***Hemi*** | ***Oreo1*** | ***EAR*** | ***EAR: Heel*** | ***EAR: Troph*** | ***EAR: CT*** | ***Cop*** | ***Scutogyrus*** | ***Bulb*** |
| --- | --- | --- | --- | --- | --- | --- | --- | --- | --- | --- | --- | --- | --- |
| 1 | 0.91 | 0.88 | 0.78 | 0.85 | 0.85 | 0.89 | 0.88 | 0.9 | 0.75 | 0.88 | 0.88 | 0.86 | 0.89 |
| 2 | 1.16 | 0.94 | 0.96 | 0.95 | 1.1 | 0.97 | 1.03 | 1.17 | 0.94 | 1.09 | 1.07 | 1.02 | 1.11 |
| 3 | 3.83 | 5.96 | 3.27 | 3.21 | 3.48 | 3.62 | 2.92 | 2.9 | 2.43 | 2.88 | 3.82 | 2.77 | 3.16 |
| 4 | 2.3 | 5.5 | 2.64 | 2.84 | 2.57 | 3.29 | 2.31 | 1.87 | 2.17 | 2.05 | 2.91 | 2.36 | 2.27 |
| 5 | 0.29 | 0.45 | 0.34 | 0.44 | 0.37 | 0.47 | 0.5 | 0.33 | 0.33 | 0.39 | 0.47 | 0.7 | 0.4 |
| 6 | 0.14 | 0.17 | 0.16 | 0.16 | 0.15 | 0.17 | 0.15 | 0.15 | 0.2 | 0.15 | 0.15 | 0.16 | 0.16 |
| 7 | 1.67 | 1.22 | 1.24 | 1.31 | 2.43 | 1.29 | 1.48 | 2.36 | 1.17 | 1.01 | 1.91 | 1.44 | 2.07 |
| 8 | 1.61 | 1.47 | 1.93 | 1.81 | 1.69 | 1.46 | 1.64 | 1.95 | 1.29 | 1.44 | 1.7 | 2.03 | 1.83 |
| 9 | 34.04 | 89.38 | 78.29 | 71.55 | 116 | 55.79 | 49.27 | 33.45 | 43.8 | 57.26 | 42.97 | 52.82 | 42.47 |
| 12 | 34.15 | 42.33 | 57.34 | 39.09 | 49.6 | 38.42 | 35.95 | 18.44 | 37.05 | 47.02 | 37.46 | 41.77 | 41.09 |
| 14 | 5.89 | 6.62 | 6.05 | 5.45 | 3.38 | 7.19 | 6.9 | 21.36 | 4.14 | 5.85 | 6.13 | 4.66 | 4.67 |
| 15 | 0.66 | 0.65 | 0.64 | 18.44 | 0.78 | 1.41 | 1.72 | 0.26 | 0.12 | 0.08 | 1.67 | 1.45 | 0.75 |
| 16 | 2.97 | 43.78 | 8.6 | 28.61 | 42.07 | 13.48 | 8.65 | 8.25 | 6.71 | 1 | 6.88 | 13.28 | 4.83 |
| 17 | 0.46 | 1.26 | 0.37 | 1.72 | 2.29 | 1.16 | 1.01 | 1.67 | 2.44 | 0.28 | 0.57 | 2.17 | 1.88 |

**Appendix 7.** Ancestral states of discrete characters inferred from a literature survey and used for character mapping. Values represent probabilities for different character states. Character IDs refer to numbers in Appendix 2.

| **Character ID** | ***Tylo*** | ***Oreo2*** | ***Halli*** | ***CPO*** | ***Hemi*** | ***Oreo1*** | ***EAR*** | ***EAR: Heel*** | ***EAR: Troph*** | ***EAR: CT*** | ***Cop*** | ***Scutogyrus*** | ***Bulb*** |
| --- | --- | --- | --- | --- | --- | --- | --- | --- | --- | --- | --- | --- | --- |
| 10a | 0 | 1 | 1 | 0 | 1 | 0 | 1 | 1 | 1 | 1 | 0 | 1 | 1 |
| 10b | 0 | 0 | 0 | 0 | 0 | 0 | 0 | 0 | 0 | 0 | 0 | 0 | 0 |
| 10c | 0 | 0 | 0 | 1 | 0 | 1 | 0 | 0 | 0 | 0 | 1 | 0 | 0 |
| 10d | 0 | 0 | 0 | 0 | 0 | 0 | 0 | 0 | 0 | 0 | 0 | 0 | 0 |
| 10e | 0 | 0 | 0 | 0 | 0 | 0 | 0 | 0 | 0 | 0 | 0 | 0 | 0 |
| 10f | 0 | 0 | 0 | 0 | 0 | 0 | 0 | 0 | 0 | 0 | 0 | 0 | 0 |
| 10g | 1 | 0 | 0 | 0 | 0 | 0 | 0 | 0 | 0 | 0 | 0 | 0 | 0 |
| 11a | 1 | 1 | 0 | 1 | 1 | 1 | 1 | 1 | 1 | 1 | 1 | 1 | 0 |
| 11b | 0 | 0 | 1 | 0 | 0 | 0 | 0 | 0 | 0 | 0 | 0 | 0 | 0 |
| 11c | 0 | 0 | 0 | 0 | 0 | 0 | 0 | 0 | 0 | 0 | 0 | 0 | 1 |
| 13a | 0 | 0 | 1 | 0 | 0 | 0 | 0.25 | 0 | 0 | 0 | 0 | 0 | 0 |
| 13b | 0 | 0 | 0 | 0.33 | 0 | 0 | 0.25 | 0 | 1 | 1 | 0 | 0 | 0 |
| 13c | 0 | 0 | 0 | 0.33 | 0 | 1 | 0.25 | 0 | 0 | 0 | 1 | 1 | 1 |
| 13d | 0 | 0 | 0 | 0 | 0 | 0 | 0 | 0 | 0 | 0 | 0 | 0 | 0 |
| 13e | 0 | 0 | 0 | 0 | 0 | 0 | 0 | 0 | 0 | 0 | 0 | 0 | 0 |
| 13f | 1 | 0 | 0 | 0 | 0 | 0 | 0 | 0 | 0 | 0 | 0 | 0 | 0 |
| 13g | 0 | 0 | 0 | 0 | 0 | 0 | 0 | 0 | 0 | 0 | 0 | 0 | 0 |
| 13i | 0 | 0 | 0 | 0 | 0 | 0 | 0.25 | 1 | 0 | 0 | 0 | 0 | 0 |
| 13j | 0 | 0 | 0 | 0.33 | 0 | 0 | 0 | 0 | 0 | 0 | 0 | 0 | 0 |
| 13k | 0 | 1 | 0 | 0 | 0 | 0 | 0 | 0 | 0 | 0 | 0 | 0 | 0 |
| 13l | 0 | 0 | 0 | 0 | 1 | 0 | 0 | 0 | 0 | 0 | 0 | 0 | 0 |
| 13m | 0 | 0 | 0 | 0 | 0 | 0 | 0 | 0 | 0 | 0 | 0 | 0 | 0 |
| 13n | 0 | 0 | 0 | 0 | 0 | 0 | 0 | 0 | 0 | 0 | 0 | 0 | 0 |
| 18a | 1 | 0.5 | 1 | 0 | 0 | 0 | 1 | 0 | 1 | 1 | 1 | 1 | 1 |
| 18b | 0 | 0.5 | 0 | 1 | 1 | 1 | 0 | 0 | 0 | 0 | 0 | 0 | 0 |
| 18c | 0 | 0 | 0 | 0 | 0 | 0 | 0 | 1 | 0 | 0 | 0 | 0 | 0 |
| 18d | 0 | 0 | 0 | 0 | 0 | 0 | 0 | 0 | 0 | 0 | 0 | 0 | 0 |
